# In-vivo MRI of noradrenergic and dopaminergic nuclei predicts future cognition and Alzheimer’s disease progression

**DOI:** 10.64898/2026.10.29.762942

**Authors:** Wenhui Zhao, Samuel J. Guzman, Hyun Joo Yoo, Kacey Ruiz, Keran Chen, Ariel Moheban, Daniel A. Nation, Alzheimer’s Disease Neuroimaging Initiative (ADNI), Martin J. Dahl, Mara Mather

## Abstract

Noradrenergic and dopaminergic nuclei modulate cognition, and post-mortem data indicate they are affected early in the neurodegenerative cascade of Alzheimer’s disease. We derived in-vivo locus coeruleus (LC) and substantia nigra-ventral tegmental area (SN-VTA) MRI integrity measures from 739 Alzheimer’s Disease Neuroimaging Initiative (ADNI) participants, examined MRI-histology correspondence in individuals with both in-vivo imaging and post-mortem data, and tested whether these measures reflect early MRI-indexed degeneration predicting later cognitive decline. LC and SN-VTA contrast predicted cognitive and behavioral function 10 years later, despite only weak associations with current function. LC- and SN-VTA-associated brain-wide atrophy patterns aligned with PET-derived norepinephrine-transporter and dopamine-receptor maps, respectively. A one-standard-deviation decrease in baseline LC contrast was associated with 84% higher five-year odds of MCI-to-AD progression, and predictive effects of the catecholaminergic nuclei were mediated by their corresponding atrophy patterns. The SN-VTA pattern was associated with less amyloid PET-related memory decline. Together, these findings establish LC and SN-VTA MRI contrast as markers of early neurodegeneration that forecast cognitive decline years before clinical onset.

## Introduction

Postmortem studies suggest that the locus coeruleus (LC), a few millimeter-sized noradrenergic structure located deep in the brainstem, is the first place Alzheimer’s disease (AD)-associated tau pathology develops, and that the pathology spreads from there to the cortex^1,2^. In line with this role, higher LC neuronal density assessed after death is associated with prior slower late-life cognitive decline, and post-mortem studies revealed LC neuronal loss in amnestic mild cognitive impairment (MCI), with further loss in AD^3,4^. However, most of our knowledge about the role of the human LC in late-life memory decline and AD relies on static post-mortem assessments, reflecting long-standing challenges with non-invasive measurements^5–7^. Furthermore, dopaminergic degeneration is a characteristic symptom in Lewy body dementias, including dementia with Lewy bodies and Parkinson’s disease^8,9^. Lewy body pathology commonly coexists with AD pathology and is associated with faster cognitive decline^10,11^. Recent studies also revealed dopaminergic abnormalities in AD, including lower striatal dopamine transporter binding in amyloid-positive individuals with MCI or AD dementia^12^. In cognitively unimpaired older adults, amyloid and tau pathology were associated with a weakening or reversal of the positive relationship between dopamine synthesis and memory performance^13^. Recently developed specialized magnetic resonance imaging (MRI) sequences can visualize small catecholaminergic (i.e., noradrenergic or dopaminergic) nuclei in living humans^14–24^, and the resulting MRI contrast in these nuclei has been shown to relate to concurrent cognition and Alzheimer’s diagnosis in older adults^25–27^. However, the postmortem data suggest that early damage in these regions should predict subsequent cortical degeneration and cognitive decline, as these regions are affected long before cortical regions^1,4,28^. To test this proposition, large multimodal longitudinal studies including both AD conversion information and scans showing contrast in catecholaminergic nuclei are needed. Informed by recent MRI physics work on the contrast mechanisms of catecholaminergic nuclei^29,30^, we identified dual-echo MRI scans that show a clear signal for noradrenergic and dopaminergic nuclei in the first wave of the Alzheimer’s Disease Neuroimaging Initiative (ADNI)^31,32^. Crucially, these scans were collected in a large cohort with many follow-ups (739 participants with 3,782 dual-echo MRI scans, with follow-up up to 20.03 years) that included postmortem LC and SN assessments and comprehensive AD biomarker and cognitive assessments, allowing us to contrast how in-vivo markers of these two neuromodulatory systems contribute to cognitive changes and AD development.

The LC is the principal source of norepinephrine for the forebrain^33^. LC neurons regulate attentional prioritization^34,35^, and LC input to the hippocampus can support synaptic plasticity and memory through noradrenergic and dopamine-sensitive mechanisms^36–40^. Its widely-branching axonal architecture may contribute to its selective vulnerability, providing a potential substrate for tau propagation^41–43^. In addition, age-related changes in tonic locus coeruleus activity may contribute to spread of tau pathology^44–46^. Eventually, loss of LC-derived norepinephrine may impair microglial amyloid-β clearance and anti-inflammatory regulation^47^. Dopaminergic populations in the SN-VTA innervate striatal, limbic, and cortical circuits involved in learning and working memory^48,49^, while VTA projections can support hippocampal memory persistence^50,51^. Dopaminergic signaling may also influence AD pathology. Increasing VTA dopamine release enhanced amyloid-β degradation and reduced prefrontal amyloid deposition^52^. Together, these findings suggest that alterations in catecholaminergic nuclei could relate to cognitive decline through both disrupted neuromodulation and disease-related changes in their brain-wide projection targets^35^.

Mechanistic animal studies identify candidate pathways, but longitudinal human studies are needed to establish how in-vivo measurements of these nuclei relate to cognitive and clinical trajectories. Functional MRI studies found that lower novelty-related LC activity and weaker LC-medial temporal connectivity were associated with subsequent amyloid-related cognitive decline in cognitively unimpaired older adults^53^, while reduced baseline VTA connectivity distinguished participants with MCI who progressed to AD dementia over 24 months^54^. Structural MRI provides complementary evidence. In older adults, LC MRI contrast has been associated with episodic memory^24,25,55^, and SN-VTA contrast with working memory^24^, supporting potentially distinguishable cognitive associations^24^. Lower T1-derived VTA gray-matter measures preceded worsening on the Clinical Dementia Rating^56^, and declining LC contrast was associated with poorer episodic memory at a later assessment^24^. LC MRI measures have also been associated with progression from amnestic MCI to dementia and clinical worsening in initially asymptomatic individuals^57,58^. A recent study revealed declining LC integrity during early AD, which was also associated with memory performance^59^. In symptomatic AD, lower LC and SN contrast were each associated with faster longitudinal cognitive decline, even when accounting for one another^60^. Despite these advances, it remains unclear how LC and SN-VTA in-vivo MRI measures associated with cognition and brain evolve over many years, and how these link to clinical progression.

The widespread projections and disease-related mechanisms of LC and SN-VTA suggest that their cognitive associations may depend on the integrity of distributed brain systems^43,61–63^. In autosomal-dominant AD, lower LC integrity was associated with greater tau burden and poorer memory^64^. In a longitudinal study, lower baseline LC integrity was associated with greater medial temporal lobe tau burden approximately three years later, and tau burden at that follow-up was associated with poorer cognitive performance^65^. Complementary multivariate work has related neuromodulatory nuclei microstructure to distributed white-matter patterns associated with tau and apolipoprotein E ε4 status^66^. These findings motivate examining how LC and SN-VTA contrast covaries with brain-wide regional-atrophy trajectories, and whether expression of the resulting structural patterns is associated with clinical progression and amyloid-related cognitive decline.

In-vivo MRI can address these questions by localizing catecholaminergic regions and linking them to longitudinal brain, molecular, and behavioral measurements. With specialized sequences, LC contrast co-localizes with neuromelanin-containing and tyrosine-hydroxylase-positive tissue in postmortem specimens^67^, while SN contrast relates to neuromelanin concentration in postmortem tissue and nigrostriatal dopamine function in-vivo^68^. The exact physical and biological sources of these signals remain uncertain. Neuromelanin-metal interactions, mobile-water content, and macromolecular properties may contribute differently depending on the acquisition protocol^69,70^. Experimental studies have demonstrated MRI contrast in noradrenergic cell groups even in the absence of neuromelanin^27,28^ and proton-density-weighted LC and SN-VTA contrast have also shown convergence with established catecholamine-sensitive acquisitions^24^. These findings motivate our use of proton-density-weighted (PDw) scans from ADNI dual-echo MRI sequence while evaluating the biological correspondence of the resulting measurements through antemortem MRI and later regional neuropathology from the same individuals.

Here, we used in-vivo dual-echo MRI scans from ADNI to quantify LC and SN-VTA contrast and evaluated longitudinal stability and correspondence with later neuropathology in the same individuals. We examined associations with concurrent and subsequent cognition over MRI-to-assessment intervals extending to approximately twenty years. We then identified brain-wide atrophy patterns associated with catecholaminergic nuclei and correlated these patterns with PET-derived catecholaminergic receptor and transporter distributions. Finally, we evaluated associations with MCI-to-AD progression, whether progression effects were mediated by brain-wide atrophy patterns, and moderation of amyloid-related memory decline.

## Results

### Overview of the cohort

We drew on 3,782 dual-echo PD-T2 MRI scans with matched T1-weighted images from 739 ADNI participants, including cognitively normal (CN) individuals and those with MCI or AD dementia. After quality control and extreme-value screening, 3,428 scans from 726 participants contributed to the LC or SN-VTA contrast ratio (CR), an in-vivo proxy for the integrity of these nuclei. Overall cohort follow-up extended to 20.03 years. Detailed sample characteristics, scan retention and repeated-imaging intervals are reported in the Supplementary Note and Supplementary Table 1; the processing workflow is shown in Supplementary Fig. 1.

**Fig. 1:**
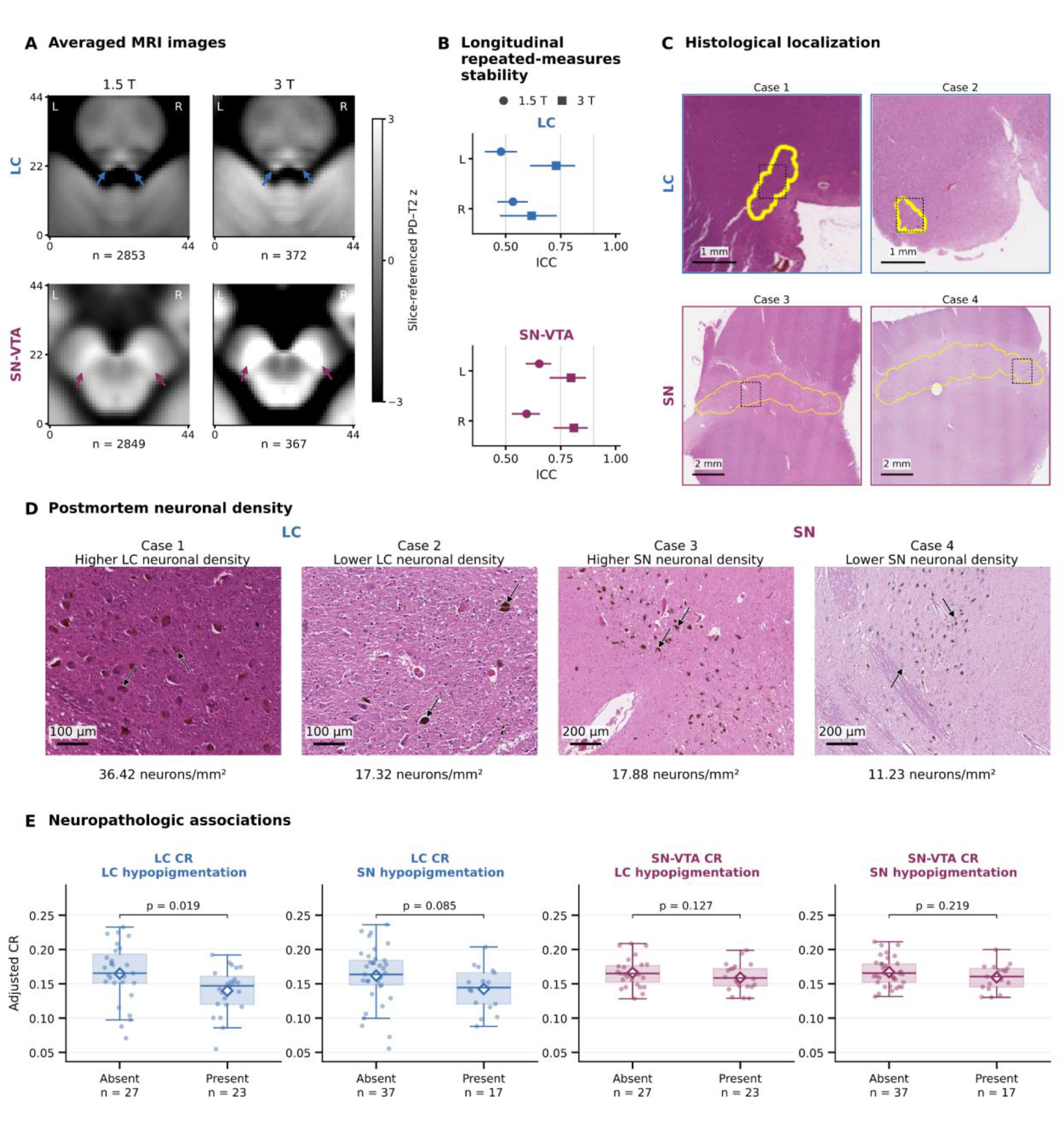
LC and SN-VTA MRI contrast characteristics, longitudinal stability, and neuropathologic associations. **a**, Native-space PD-T2 images, averaged across scans. Mean images for LC and SN-VTA are shown separately for 1.5 T and 3 T. For each scan, a 45 × 45-voxel native-space axial patch was centered on the bilateral LC or SN-VTA mask centroid. The color scale denotes slice-wise standardized intensity estimates. The n beneath each image denotes the number of scans. Blue and plum arrows indicate the hyperintense LC and SN–VTA locations, respectively. **b**, Longitudinal repeated-measures stability. Intraclass correlation coefficient (ICC) point estimates and 95% confidence intervals from 5,000 bootstrap simulations are shown for LC and SN-VTA CR, separately by field strength and hemisphere. Vertical reference lines denote ICC values of 0.50, 0.75, and 0.90. **c**, Histological localization. Hematoxylin and eosin (H&E)-stained tissue sections show LC localization in Cases 1 and 2 and SN localization in Cases 3 and 4. Yellow outlines indicate the localized regions. Scale bars, 1 mm for LC and 2 mm for SN. Black dashed rectangles indicate locations of the higher-magnification fields in d, which are displayed in a different orientation. **d**, Postmortem neuronal density. Representative tissue images show higher and lower neuronal density within the LC pair (Cases 1 and 2) and the SN pair (Cases 3 and 4). Scale bars, 100 µm for LC and 200 µm for SN. Arrows indicate illustrative neuronal cell bodies. **e**, Neuropathologic associations. Covariate-adjusted bilateral LC or SN-VTA MRI CR values are shown split by the presence or absence of gross hypopigmentation in the LC or SN. Boxes show the interquartile range, center lines show the median, whiskers extend to 1.5 times the interquartile range, points represent participants, and white diamonds show group means. Displayed CR values were obtained from the same fitted models used to prepare CR for inference. Participant-specific variation was retained, and the corresponding mean unadjusted CR was added back for display. CR, contrast ratio; ICC, intraclass correlation coefficient; LC, locus coeruleus; MRI, magnetic resonance imaging; PD-T2, proton-density-minus-T2-weighted difference signal; SN, substantia nigra; SN-VTA, substantia nigra-ventral tegmental area.

**Fig. 2:**
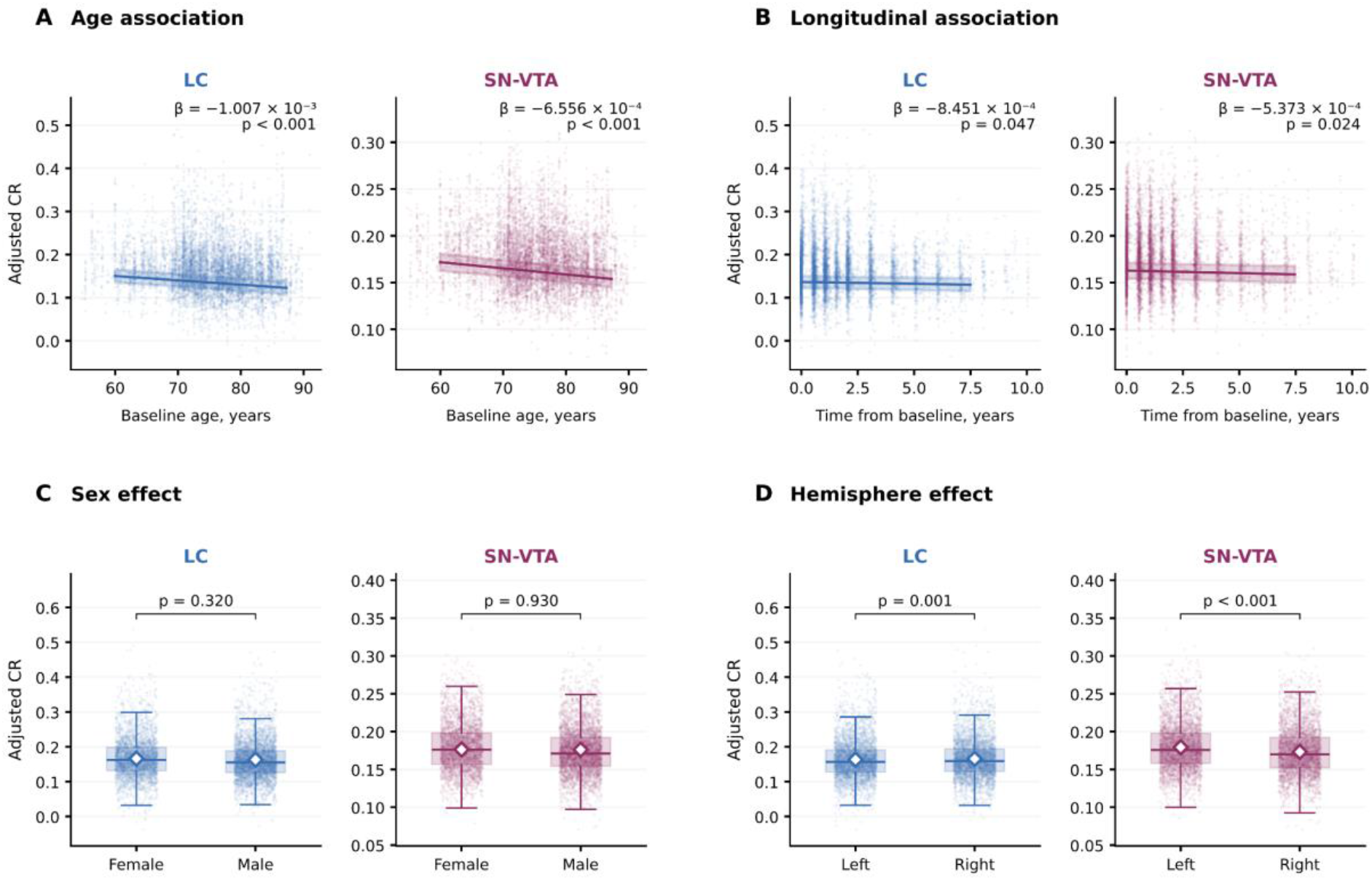
Associations of age, longitudinal time, sex and hemisphere with LC and SN-VTA CR. **a**, Age association. Covariate-adjusted CR and fixed-effect predictions are shown across baseline age. **b**, Longitudinal association. Covariate-adjusted CR and fixed-effect predictions are shown across time since baseline. Solid lines show model-predicted CR and shaded bands show 95% confidence intervals. The displayed β coefficients denote adjusted changes in CR per year. **c**, Sex contrast. Covariate-adjusted CR distributions are shown for female and male participants. **d**, Hemisphere contrast. Covariate-adjusted CR distributions are shown for the left and right hemispheres. In c and d, boxes show the interquartile range, center lines show the median, whiskers extend to 1.5 times the interquartile range, and points show repeated hemisphere-specific observations. White diamonds and vertical bars denote estimated marginal means and 95% confidence intervals. P values correspond to the male-minus-female and right-minus-left model contrasts, respectively. CR, contrast ratio; LC, locus coeruleus; SN-VTA, substantia nigra-ventral tegmental area.

### Catecholaminergic CR characteristics, longitudinal stability and neuropathologic associations

Before examining associations with cognition and AD progression, we characterized the anatomical localization of the LC and SN-VTA MRI signal, the stability of catecholaminergic CR across repeated scans, and its correspondence with postmortem neuropathology.

The availability of scans at both field strengths allowed us to examine cross-field correspondence. CRs were positively associated between 1.5 T and 3 T for both LC (β = 0.0169, P = 7.54 × 10⁻⁴) and SN-VTA (β = 0.0128, P = 9.45 × 10⁻⁶; Supplementary Fig. 3), indicating assessments of both field strengths capture shared information.

Averaged PD-T2 images showed a bilateral focal hyperintensity at the anatomical locations of the LC and SN-VTA^71^ across 1.5 T and 3 T acquisitions (Fig. 1a; Supplementary Fig. 4). The averages included 2,853 LC and 2,849 SN-VTA scans at 1.5 T, and 372 LC and 367 SN-VTA scans at 3 T. Left- and right-hemisphere CR distributions largely overlapped at 1.5 T and 3 T (Supplementary Fig. 2).

We next assessed the temporal stability of individual differences in CR across repeated MRI examinations as a reliability proxy^72^. At 1.5 T, the LC intraclass correlation coefficient (ICC) was 0.478 for the left hemisphere (95% CI, 0.409-0.547) and 0.533 for the right hemisphere (95% CI, 0.466-0.596). The corresponding SN-VTA ICCs were 0.653 on the left (95% CI, 0.595-0.703) and 0.595 on the right (95% CI, 0.532-0.652). At 3 T, LC ICCs were 0.729 on the left (95% CI, 0.615-0.812) and 0.618 on the right (95% CI, 0.477-0.728). The corresponding SN-VTA ICCs were 0.797 on the left (95% CI, 0.704-0.860) and 0.810 on the right (95% CI, 0.721-0.869) (Fig. 1b; Supplementary Table 2), together indicating reliable MRI-based assessments of catecholaminergic integrity.

De-pigmentation of catecholaminergic nuclei (i.e., a loss of their characteristic neuromelanin-driven color) is a prominent feature of several neurodegenerative diseases evident in neuropathological investigations^72^. We related in-vivo MRI-derived CR to loss of pigmentation observed at autopsy in the same participants (assessed on average 4.06 years later; SD 3.36 years; range 0.15-11.29 years). Representative LC and SN tissue images and examples of higher and lower neuronal density are shown in Fig. 1c,d and Supplementary Fig. 6. Among 50 participants with LC assessments, those with LC hypopigmentation had lower LC CR than those without hypopigmentation (β = −0.036; unadjusted P = 0.019; Fig. 1e), suggesting the validity of the imaging approach. LC CR was also lower in participants with post-mortem substantia nigra hypopigmentation, but the association was not nominally significant (β = −0.021; P = 0.085; 37 absent and 17 present). Neither post-mortem LC hypopigmentation nor SN hypopigmentation was nominally associated with SN-VTA CR (LC hypopigmentation: β = −0.010; P = 0.127; substantia nigra hypopigmentation: β = −0.008; P = 0.219; Supplementary Table 3), while numerical differences pointed in the same direction.

We also examined how pigmentation preservation related to microscopic measures of neuronal density and pathology. In the LC, pigmentation preservation showed a positive association with neuron density (r = 0.531, P = 0.006). It showed non-significant positive associations with amyloid absence (r = 0.351, P = 0.116), tau absence (r = 0.060, P = 1.000), and α-synuclein/Lewy-body absence (r = 0.226, P = 0.376). In the substantia nigra, pigmentation preservation was not significantly associated with amyloid absence (r = 0.067, P = 1.000) and tau absence (r = 0.271, P = 0.274), but showed an association with α-synuclein/Lewy-body absence (r = 0.542, P = 0.011). Pigmentation preservation was not significantly associated with neuron density (r = −0.094, P = 0.635). (Supplementary Table 4 and Supplementary Fig. 5)

Partial least squares correlation (PLSC) analyses relating adjusted CR to all five neuropathological measures did not identify a significant overall association for LC (n = 20; P = 0.452) or SN-VTA (n = 23; P = 0.302; Supplementary Fig. 5 and Supplementary Tables 5 and 6). Taken together, higher in-vivo LC CR was associated with greater LC pigmentation preservation at autopsy, which was in turn associated with higher neuronal density. In contrast, SN-VTA CR was not significantly associated with SN pigmentation, potentially reflecting differences in anatomical coverage between the MRI-defined SN-VTA region and the SN examined at autopsy.

### Demographic, longitudinal and hemispheric associations with catecholaminergic CR

We next examined how LC and SN-VTA CR varied with age and longitudinal time. Older baseline age was associated with lower LC and SN-VTA CR (LC: β = −1.007 × 10^−3^, P < 0.001; SN-VTA: β = −6.556 × 10^−4^, P < 0.001; Fig. 2a). After accounting for these between-participant age differences, CR also decreased with time since baseline (LC: β = −8.451 × 10^−4^ per year, P = 0.047; SN-VTA: β = −5.373 × 10^−4^ per year, P = 0.024; Fig. 2b). Sex was not significantly associated with either CR measure (Fig. 2c). Right LC CR was higher than left LC CR (β = 0.002, P = 0.001), whereas right SN-VTA CR was lower than left SN-VTA CR (β = −0.006, P < 0.001; Fig. 2d). APOE ε4 allele count was not significantly associated with LC CR (β = 0.001, P = 0.569) or SN-VTA CR (β = 0.001, P = 0.335) after adjustment for covariates. Complete fixed-effect estimates are reported in Supplementary Table 7. LC and SN-VTA CR were positively associated with each other (β = 0.011; P = 6.527 × 10^−23^; Supplementary Fig. 7). Separate analyses by scanner manufacturer and field strength consistently showed higher left than right SN-VTA CR, whereas LC CR was higher on the right in Siemens 1.5-T scans but higher on the left in Siemens 3-T scans (Supplementary Table 8).

### Concurrent, prospective and lag-dependent catecholaminergic CR associations with cognitive, clinical and AD-related measures

We first examined associations of LC and SN-VTA CR with cognitive performance, AD-related biological measures, clinical impairment and everyday functioning assessed within 30 days of MRI scans. Nine concurrent associations from eight of the outcomes survived false-discovery-rate (FDR) correction for the two catecholaminergic nuclei (Fig. 3a; Supplementary Table 9). No association with cognitive measures survived correction. Higher SPARE-AD scores, indicating an AD-like pattern of brain atrophy, were associated with lower LC CR (β = −0.006; P_FDR_ < 0.001) and lower SN-VTA CR (β = −0.005; P_FDR_ < 0.001), and an older-appearing brain, measured by the SPARE-BA age gap score, was associated with lower SN-VTA CR (β = −0.003; P_FDR_ < 0.001). Larger right and left hippocampal volumes (both β = 0.002; both P_FDR_ = 0.040) and right and left entorhinal volumes (both β = 0.003; both P_FDR_ = 0.018) were associated with higher SN-VTA CR. Greater impairment in the Clinical Dementia Rating (CDR) Home and Hobbies domain was associated with lower SN-VTA CR (β = −0.002; P_FDR_ = 0.031), whereas greater difficulty on the Functional Activities Questionnaire Current Events item was associated with lower LC CR (β = −0.003; P_FDR_ = 0.008). Thus, higher CRs for both nuclei were associated with less age- and AD-related brain and behavior patterns. Sensitivity test results, including a model that tested LC and SN-VTA unique effects, are reported in Supplementary Note, Supplementary Fig. 8 and Supplementary Table 10.

In addition to evaluating how catecholaminergic integrity is associated with concurrent cognition, we leveraged the longitudinal nature of the dataset to test if baseline LC and SN-VTA CR is linked to performance at subsequent time points (180-1,095 days after the initial scan). Specifically, we tested whether LC and SN-VTA CR showed associations with subsequent outcomes that explained variance beyond the variance explained by associations with concurrent status by adjusting for the corresponding baseline assessment in the analysis. Twenty prospective associations reflecting 18 outcomes survived FDR correction for LC or SN-VTA (Fig. 3b; Supplementary Table 11). Higher LC CR was associated with better subsequent performance on the memory composite (β = 0.028; P_FDR_ = 0.014) and language composite (β = 0.035; P_FDR_ = 0.011). Test-level associations included Rey Auditory Verbal Learning Test (RAVLT) immediate learning (β = 0.055; P_FDR_ = 0.002), 30-min delayed recall (β = 0.051; P_FDR_ = 0.004), and recognition discrimination (β = 0.054; P_FDR_ = 0.010); together with animal fluency (β = 0.058; P_FDR_ = 0.004), and vegetable fluency (β = 0.078; P_FDR_ = 0.002). Together, these findings are consistent with noradrenaline’s known role in memory processing^24,73–75^. By contrast, higher SN-VTA CR was associated with better subsequent Digit Span Backward performance (β = 0.079; P_FDR_ = 0.025) and Boston Naming Test performance (β = 0.058; P_FDR_ = 0.025), in line with a dopaminergic role in working memory and processing speed^24,76^. For AD-related biological, clinical, and functional measures, higher LC CR was associated with larger subsequent right and left entorhinal volumes (β = 0.017 and 0.015; both P_FDR_ = 0.044), one of the first cortical targets of AD-related tau accumulation^77,78^. Higher LC and SN-VTA CR were each associated with less subsequent impairment in CDR Home and Hobbies (LC: β = −0.037, P_FDR_ = 0.050; SN-VTA: β = −0.036, P_FDR_ = 0.043) and CDR Sum of Boxes (LC: β = −0.027, P_FDR_ = 0.050; SN-VTA: β = −0.027, P_FDR_ = 0.043). Higher SN-VTA CR was additionally associated with less subsequent impairment in CDR Community Affairs (β = −0.051; P_FDR_ = 0.012) and the Functional Activities Questionnaire items assessing Games or Hobbies (β = −0.045; P_FDR_ = 0.043), Current Events (β = −0.037; P_FDR_ = 0.049), Appointments or Medications (β = −0.033; P_FDR_ = 0.049) and Travel or Transportation (β = −0.047; P_FDR_ = 0.018).

Association estimates were similar when removing the upper follow-up limits, indicating the associations were not substantially affected by including assessments with higher uncertainty. After accounting for the respective other catecholaminergic system, selected LC associations with memory, language and verbal fluency and SN-VTA associations with community functioning and travel or transportation remained FDR significant (Supplementary Note; Supplementary Figs. 8 and 9; Supplementary Tables 10 and 11), pointing to unique associations.

We then examined how the CR-outcome associations varied with the interval between MRI and cognitive/functioning assessment. Lag-dependent models were fitted for all 38 measures. Figure 3C displays the 36 LC and SN-VTA curves for the 18 outcomes with the highest proportion of observed follow-up time covered by statistically significant intervals (see Methods); the remaining outcomes are shown in Supplementary Figs. 10-11. Among the displayed results, all 18 outcomes and 32 curves contained at least one interval in which the 95% simultaneous confidence band excluded zero and that met the display-duration criterion. Higher LC CR showed positive associations with memory, RAVLT immediate learning, delayed recall, recognition discrimination, animal fluency, and bilateral entorhinal volumes at later MRI-to-assessment intervals. Higher SN-VTA CR was associated with better memory and RAVLT performance and less subsequent clinical and functional impairment at later intervals. For the memory composite, the zero-excluding intervals were 7.85-18.80 years for LC and 8.96-18.80 years for SN-VTA. Brain-imaging outcomes also showed associations at later intervals. Higher LC CR was associated with larger bilateral hippocampal and entorhinal volumes and lower SPARE-AD scores, indicating a less AD-like atrophy pattern. Higher SN-VTA CR was associated with larger left hippocampal volume. Both catecholaminergic CR measures were associated with lower white-matter hyperintensity (WMH) burden, an MRI marker of cerebral small-vessel disease^79^. Some zero-excluding intervals extended to approximately 18.8 years for regional volumes and SPARE-AD and 20.0 years for WMH burden (Fig. 3c; Supplementary Fig. 11; Supplementary Table 12). Thus, positive associations between the initial structure of catecholaminergic nuclei and selected cognitive and functional outcomes as well as brain atrophy were found at MRI-to-assessment intervals extended beyond a decade, even when controlling for baseline associations.

In sensitivity analyses, we assessed whether the late-follow-up findings depended on assessment selection, diagnosis at MRI or adjustment for the other CR measurement by running separate models with a minimum 180-day MRI-to-assessment interval, restricting analyses to participants with CN or MCI at MRI, or including both CR measurements. Across these three full-range specifications, 25-26 LC and 21-25 SN-VTA curves among the 38 outcomes contained at least one interval in which the simultaneous 95% confidence band excluded zero. Positive memory-composite associations at later follow-up were supported across all evaluated specifications, meaning that, for instance, catecholaminergic nuclei CRs predicted subsequent outcomes for both those who were cognitively normal and those with MCI or AD at baseline, even though the power was weaker for these subgroups.

After mutual adjustment for the respective other catecholaminergic system, 25 LC and 24 SN-VTA curves contained a zero-excluding interval, including positive associations with the memory composite and RAVLT immediate learning, delayed recall and recognition for both CRs (Supplementary Figs. 12-23; Supplementary Tables 12 and 13). Thus, LC and SN-VTA CR showed associations with future memory outcomes after mutual adjustment. The model curves evaluated over the 5th-95th-percentile lag range to control the extreme values were and provided in Supplementary Figs. 24-27.

**Fig. 3:**
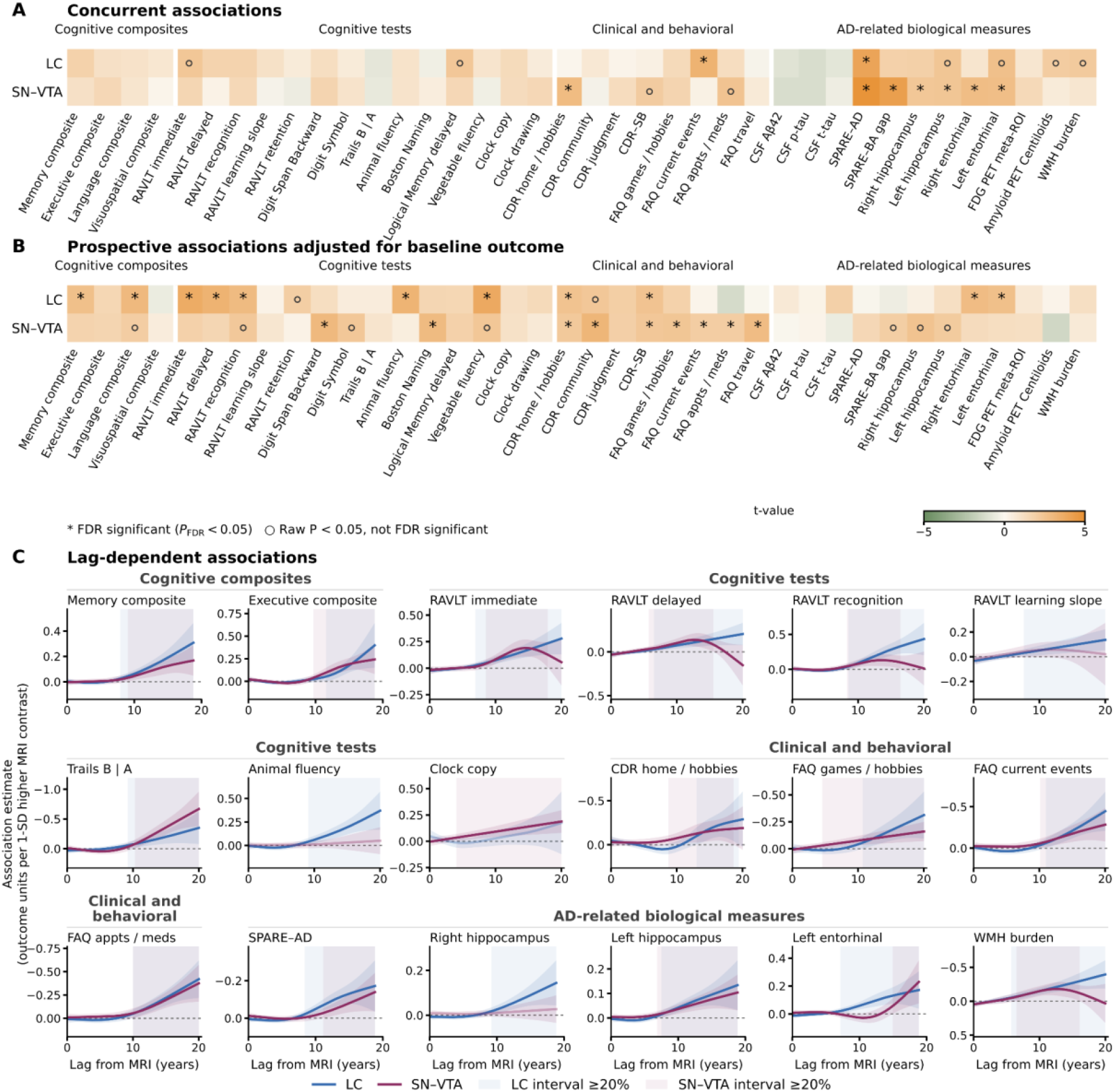
Concurrent, prospective and lag-dependent associations of LC and SN-VTA CR with cognitive, clinical and AD-related measures. **a**, Concurrent associations. Heatmaps show t-statistics from separate LC and SN-VTA mixed-effects models relating each phenotype or biomarker to LC or SN-VTA CR. Assessments and MRI scans were collected within 30 days. **b**, Prospective associations adjusted for baseline outcomes. Heatmaps show t-statistics for associations between bilateral LC or SN-VTA CR and a subsequent phenotype or biomarker, adjusted for its corresponding baseline value. Future assessments occurred 180-1,095 days after the MRI scan. For the prospective and lag-dependent analyses in b and c, bilateral CR was standardized separately for each nucleus and outcome within each MRI field strength. In a and b, columns are grouped into four prespecified outcome families: harmonized cognitive composites, test-level or derived cognitive measures, AD-related biological measures, and clinical or behavioral measures. Cell color shows t-values: Signs were reversed for color mapping for measures where lower values indicate better outcomes. Therefore, for all measures, orange indicates an association of higher CR with better outcomes and green the opposite association. FDR-adjusted P values are denoted P_FDR_. Asterisks indicate P_FDR_ < 0.05; hollow circles indicate nominal P < 0.05 but P_FDR_ ≥ 0.05. **c**, Lag-dependent associations. Generalized additive model (GAM) curves are shown for the 18 outcomes with the highest proportion of observed follow-up time covered by statistically significant intervals. Curves show the estimated association with future outcome per one-standard-deviation higher bilateral LC or SN-VTA CR as a function of elapsed time after MRI. LC curves are shown in blue and SN-VTA curves in plum. Bands indicate 95% simultaneous confidence intervals within each curve, derived from 10,000 simulations. Pale vertical shading denotes individual zero-excluding intervals lasting at least 20% of the corresponding curve’s observed lag span. Shorter intervals remain reported in Supplementary Table 12. Curves without a zero-excluding interval use reduced line opacity (0.35). For measures where lower values indicate better outcomes, y-axes are reversed so that negative coefficients appear above zero; the original coefficient values and signs are retained. On the axes, upward positions indicate associations with better outcomes. For analyses including both catecholaminergic systems in a common model, see Supplementary Figs. 8 and 20-23. Aβ42, amyloid-β42; AD, Alzheimer’s disease; CDR-SB, Clinical Dementia Rating Sum of Boxes; CR, contrast ratio; CSF, cerebrospinal fluid; FDG, fluorodeoxyglucose; FDR, false discovery rate; FAQ, Functional Activities Questionnaire; GAM, generalized additive model; LC, locus coeruleus; MRI, magnetic resonance imaging; PET, positron emission tomography; p-tau, phosphorylated tau; RAVLT, Rey Auditory Verbal Learning Test; SD, standard deviation; SN-VTA, substantia nigra-ventral tegmental area. SPARE-AD, Spatial Pattern of Abnormality for Recognition of Early Alzheimer’s disease; SPARE-BA, Spatial Pattern of Abnormality for Recognition of Brain Aging; t-tau, total tau; WMH, white-matter hyperintensity.

### Brain-wide atrophy patterns associated with LC and SN-VTA trajectories

To examine how LC and SN-VTA CR covaried with distributed brain atrophy, a marker for late-life neurodegeneration^80^, we used multivariate statistics (PLSC) to relate each nucleus’s CR to MRI-derived volumes across 145 brain regions. Separate analyses related CR levels to regional-volume levels, represented by model intercepts, and CR rates of change to regional-volume rates of change. Corresponding left- and right-hemisphere CR estimates were averaged for each analysis. PLSC assigns each region a weight, or salience; a participant’s brain score summarizes the resulting weighted combination of regional-volume estimates. All four analyses identified a significant pattern of covariation (Fig. 4a,b).

For CR levels (intercept), the correlation between the brain score and bilateral CR was r = 0.228 for the LC (n = 697; P_perm_ < 0.001) and r = 0.266 for SN-VTA (P_perm_ < 0.001). For rates of change, the corresponding correlations were r = 0.178 for LC (N = 538; P_perm_ = 0.007) and r = 0.178 for SN-VTA (n = 539; P_perm_ = 0.006). This suggests that individual differences in the level as well as rate of change of catecholaminergic systems are consistently associated with markers of brain-wide atrophy.

Bootstrap ratios (BSRs) summarize the stability of regional weights across participant resamples and can be interpreted akin to *z*-values. The displayed maps and anatomical summaries (Fig. 4) highlight regions with |BSR| ≥ 2.56. The largest positive weights among regions meeting this stability threshold in the LC intercept pattern were located in the bilateral posterior insula, left inferior temporal cortex, left central operculum, left anterior insula, and left parahippocampal cortex. Positive subcortical contributions included the left amygdala and hippocampus, whereas the caudate nuclei showed negative saliences. Thus, higher fitted LC CR was associated with a multiregional pattern of larger insular, opercular, and medial-temporal volumes together with relatively smaller caudate volumes. The SN-VTA intercept pattern was dominated by positive saliences in bilateral anterior and posterior insula, right frontal operculum and right planum temporale. The largest subcortical contributions were bilateral pallidum and putamen. Higher SN-VTA CR was therefore associated with larger volumes across a pattern involving insular and opercular cortex, pallidum and putamen, regions with prominent dopaminergic modulation^81^. The contributors to the LC rate-of-change pattern were cortical, including bilateral cuneus, left inferior occipital cortex, right medial precentral cortex and right lingual cortex. The corresponding SN-VTA contributors included right medial frontal, medial precentral, superior occipital, anterior cingulate, entorhinal, and superior temporal cortices (Fig. 4a,b, Supplementary Fig. 28; Supplementary Tables 14 and 15). The BSR vectors for intercept and change patterns showed modest positive correlations (LC: r = 0.281; SN-VTA: r = 0.360), suggesting some spatial consistency between the two patterns.

The intercept and rate-of-change associations for both catecholaminergic nuclei remained significant after accounting for the corresponding CR parameter of the other nucleus (Supplementary Note; Supplementary Fig. 29; Supplementary Tables 15 and 16). These results support unique structural associations for both CR measurements after accounting for their shared variation.

### CR-related brain-wide atrophy patterns show spatial correspondence with catecholaminergic receptor and transporter maps

To test whether the cortical atrophy patterns associated with LC and SN-VTA CR aligned with the spatial distributions of norepinephrine transporters and dopamine receptors and transporters, we compared their cortical weights (see Fig. 4 A/B) with six PET maps^82^. The LC intercept LV1 showed one FDR-significant association across these six maps. Its cortical salience distribution was positively correlated with the [^11^C]MRB norepinephrine-transporter (NET) map from Hesse et al.^83^ (ρ = 0.403; P_FDR_ = 0.028), suggesting that brain regions whose atrophy was related to LC CR also housed more transporters. The other MRB map, dopamine transporter (DAT), dopamine D1-like receptor (D1), and two dopamine D2-family receptor (D2) maps did not survive FDR correction for LC LV1. Additional exploratory analyses also tested additional maps, none of which survived FDR correction for LC LV1 (Supplementary Fig. 30). The SN-VTA intercept LV1 showed positive spatial correspondence with three dopaminergic maps: D1-like receptor availability measured with [^11^C]SCH23390 (ρ = 0.491; P_FDR_ = 0.001), D2-family receptor availability measured with [^11^C]FLB457 (ρ = 0.597; P_FDR_ = 0.003), and D2/3 receptor availability measured with [^18^F]fallypride (ρ = 0.572; P_FDR_ = 0.003). Neither the NET map nor the DAT map survived FDR correction for SN-VTA LV1. These findings link the LC-related cortical atrophy pattern to the norepinephrine-transporter distribution and the SN-VTA-related pattern to dopamine-receptor distributions (Fig. 4c), consistent with a potential neuroprotective catecholaminergic role^47,84,85^. Among the 16 additional receptor, transporter or binding-site maps examined in exploratory analyses, eight showed positive correspondence with the SN-VTA intercept pattern after FDR correction, whereas none survived correction for the LC pattern (Supplementary Fig. 30). Simultaneous adjustment for the three dopamine-receptor maps reduced the magnitudes of all eight initially significant exploratory correlations, and none remained significant (Supplementary Table 17). This indicates that their spatial variations are shared with dopamine-receptor distributions.

After accounting for the respective other catecholaminergic nucleus’s CR, all three SN-VTA dopamine-receptor correspondences remained FDR-significant. The unique LC pattern retained a positive correspondence with the Hesse et al. NET map, but did not meet the FDR threshold (ρ = 0.361; P = 0.008; P_FDR_ = 0.050; Supplementary Note; Supplementary Fig. 29; Supplementary Tables 16 and 17).

**Fig. 4:**
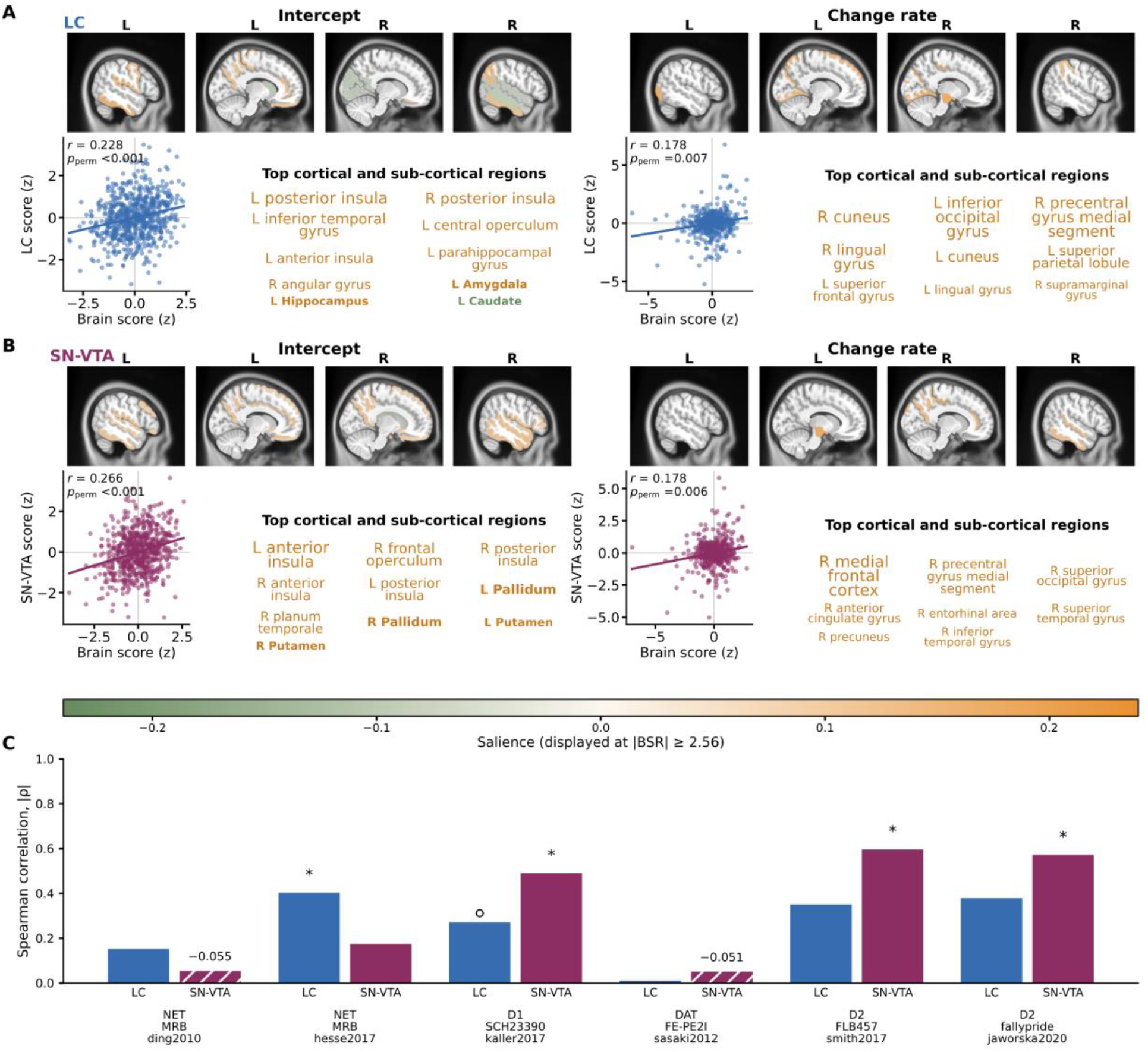
Whole-brain structural atrophy patterns associated with LC and SN-VTA trajectories and their spatial correspondence with PET-derived neurotransmitter maps. **a**, LC structural patterns. PLSC results for the bilateral LC trajectory intercept and rate of change. Brain maps show salience values for regions satisfying |BSR| ≥ 2.56 (i.e., stable contributions). Positive saliences indicate volume parameters that covaried in the same direction as the LC parameter; negative saliences indicate inverse covariation. Scatterplots show standardized participant brain scores against standardized LC scores. Each point represents one participant; lines are descriptive ordinary-least-squares fits. Pearson’s r summarizes the score association, whereas P_perm_ is the permutation P value. **b**, SN-VTA structural patterns. Corresponding PLSC results for the bilateral SN-VTA trajectory intercept and rate of change. Display conventions are the same as in a. In a and b, the anatomical-name summaries show up to ten parcels satisfying |BSR| ≥ 2.56, selected by absolute salience, with at most four subcortical names and additional eligible cortical names filling unused places. Font size increases with absolute salience, text color indicates salience sign, and subcortical names are bold. **c**, Spatial correspondence with neurotransmitter receptor and transporter maps. Bars show the absolute Spearman correlation (|ρ|) between the unthresholded cortical saliences from the LC or SN-VTA intercept LV1 and six PET maps. Hatched bars and signed numerical labels indicate negative correlations. Asterisks indicate P_FDR_ < 0.05; hollow circles indicate nominal P < 0.05 without passing the FDR. Two-sided P values were estimated from 10,000 Moran-spectral surrogate maps. BSR, bootstrap ratio; CR, contrast ratio; D1, dopamine D1-like receptor; D2, dopamine D2-family receptor; DAT, dopamine transporter; FDR, false discovery rate; LC, locus coeruleus; LV, latent variable; NET, norepinephrine transporter; PET, positron emission tomography; PLSC, partial least squares correlation; SN-VTA, substantia nigra-ventral tegmental area.

### Lower LC CR was associated with higher five-year AD progression odds

We next examined whether catecholaminergic CRs at the baseline MRI assessment were associated with diagnostic progression over the following five years. The MCI-to-AD analysis included 232 participants with MCI and a paired baseline LC or SN-VTA scan, of whom 166 progressed to AD within five years, and 66 remained free of dementia with at least five years of follow-up. One SD lower LC CR was associated with approximately 84% higher odds of progression (OR = 1.835; 95% CI, 1.222-2.754; P_FDR_ = 0.007). SN-VTA CR was not significantly associated with MCI-to-AD progression (OR = 1.223; 95% CI, 0.849-1.761; P_FDR_ = 0.279; Fig. 5a, upper row).

The any-progression analysis included 335 participants classified as CN or MCI with a matched baseline LC or SN-VTA scan, of whom 180 experienced a diagnostic transition from CN to MCI or dementia, or from MCI to dementia, within five years; 155 remained free of progression with at least five years of follow-up. Per one SD lower CR, the LC model gave OR = 1.460; 95% CI, 1.072-1.989 (P_FDR_ = 0.033); the SN-VTA model gave OR = 1.366; 95% CI, 0.994-1.877 (P_FDR_ = 0.054; Fig. 5a, lower row). Complete model results are reported in Supplementary Table 18.

The LC associations with both progression outcomes persisted after accounting for SN-VTA CR in the same model (Supplementary Note; Supplementary Fig. 31; Supplementary Tables 18 and 19). Together, these findings suggest that individual differences in MRI markers of catecholaminergic integrity have high utility for predicting future cognitive status and the development of AD.

### Catecholaminergic CR showed indirect associations with AD progression through their brain-atrophy patterns

We next tested whether the catecholaminergic nuclei-related brain-atrophy patterns identified by PLSC (see Fig. 4 a,b) statistically linked CR to diagnostic progression (Fig. 5b). The mediation analyses included 232 participants for MCI-to-AD and 335 for any progression. The mediator was the brain score for the corresponding catecholaminergic CR intercept pattern. Models included the covariates used in the corresponding progression analyses, together with total parenchymal volume (TPV; total gray- and white-matter volume) to account for overall brain atrophy.

For MCI-to-AD, the association between LC CR and the brain score was positive (a = 0.246) and the association between the brain score and progression, conditional on LC CR was negative (b = −0.600). The LC indirect effect was −0.148 on the latent-probit scale (95% CI, −0.243 to −0.052; P_FDR_ = 0.005). In the SN-VTA model, a = 0.123 and b = −0.692, yielding an indirect effect of −0.085 (95% CI, −0.168 to −0.002; P_FDR_ = 0.044; Fig. 5b, left column).

For any progression, the indirect effect was −0.091 for LC (95% CI, −0.149 to −0.034; P_FDR_ = 0.004) and −0.062 for SN-VTA (95% CI, −0.120 to −0.004; P_FDR_ = 0.035; Fig. 5b, middle column).

After adjustment for the brain score and TPV, the direct LC path remained negative in the MCI-to-AD model (c′ = −0.214; 95% CI, −0.410 to −0.018; P = 0.033), but the other direct-effect intervals included zero. We also fitted models without TPV adjustment and found that 57% of the mediation effect for SN-VTA was explained by whole brain volume change (Fig. 5b, right column; and detailed in the Supplementary Note). All four indirect effects met their FDR criterion. Complete effects are reported in Supplementary Table 20 and Supplementary Figs. 32-33. Re-estimating the PLSC patterns in participants classified as CN or MCI retained all eight indirect associations across the two regions, two progression outcomes and models with or without TPV adjustment (Supplementary Table 21). Together, these findings suggest that distributed brain-volume patterns consistently link catecholaminergic CR with subsequent diagnostic progression, even when the patterns are identified in participants without AD and after accounting for overall brain volume.

In sensitivity analyses requiring a T1 scan after the LC/SN-VTA MRI and excluding progression events occurring at or before that scan, the LC indirect associations remained FDR significant, whereas the corresponding SN-VTA confidence intervals included zero. The LC association was consistent when using an LC-associated brain volume pattern established without information from AD patients.(Supplementary Note; Supplementary Tables 19 and 22).

### The SN-VTA-related volume pattern was associated with less amyloid-related memory decline

Finally, we tested whether catecholaminergic CR or their corresponding brain-atrophy patterns modified the association between amyloid PET burden and memory change over time. The amyloid-related memory analyses included 122 participants contributing 562 memory assessments. After accounting for CR, the corresponding brain-atrophy pattern score, and TPV, amyloid-positive participants showed faster annual memory decline than amyloid-negative participants (LC model: β = −0.137; SN-VTA model: β = −0.136; both P < 0.001). In the continuous-burden models, each one-SD increase in amyloid burden, measured on the Centiloid scale, was associated with more negative annual memory change (β = −0.076 in both models; both P < 0.001).

Neither catecholaminergic CR measurement significantly modified the association between binary amyloid status and annual memory change after FDR correction (LC: β = 0.020; P_FDR_ = 0.175; SN-VTA: β = −0.021; P_FDR_ = 0.175). The corresponding continuous-Centiloid interactions were also not significant after FDR correction (LC: β = 0.011; P_FDR_ = 0.336; SN-VTA: β = 0.005; P_FDR_ = 0.565; Fig. 5c, forest plots; full results in Supplementary Fig. 34 and Supplementary Table 23).

The LC-related brain-wide atrophy pattern (quantified through brain scores) also showed no significant interaction with amyloid burden and time after FDR correction. Its interaction estimate was 0.029 for binary amyloid status (P_FDR_ = 0.132) and 0.015 for continuous Centiloids (P_FDR_ = 0.148; Fig. 5c, forest plots).

For the SN-VTA-related brain scores, the amyloid-associated difference in annual memory change was less negative at higher scores (i.e., less atrophy). The brain-score-by-amyloid-status-by-time interaction was positive (β = 0.088; P_FDR_ = 0.001). The corresponding interaction was also positive when amyloid burden was measured continuously in Centiloids (β = 0.045; P_FDR_ = 0.002; Fig. 5c, forest plots; Supplementary Tables 19 and 23).

The model-based trajectories illustrated a smaller amyloid-associated difference in memory decline at higher SN-VTA-related brain scores, whereas amyloid-positive participants with high scores showed comparatively attenuated decline. The continuous response surface similarly showed that a higher SN-VTA-related PLSC LV1 brain scores were associated with a less negative adjusted annual memory change at higher Centiloid burden (Fig. 5c). These associations remained significant after adjustment for SN-VTA CR and TPV. The corresponding LC interactions did not survive FDR correction. In sum, these findings suggest that the brain-volume pattern associated with SN-VTA CR may reflect greater resilience to amyloid-related memory decline.

**Fig. 5:**
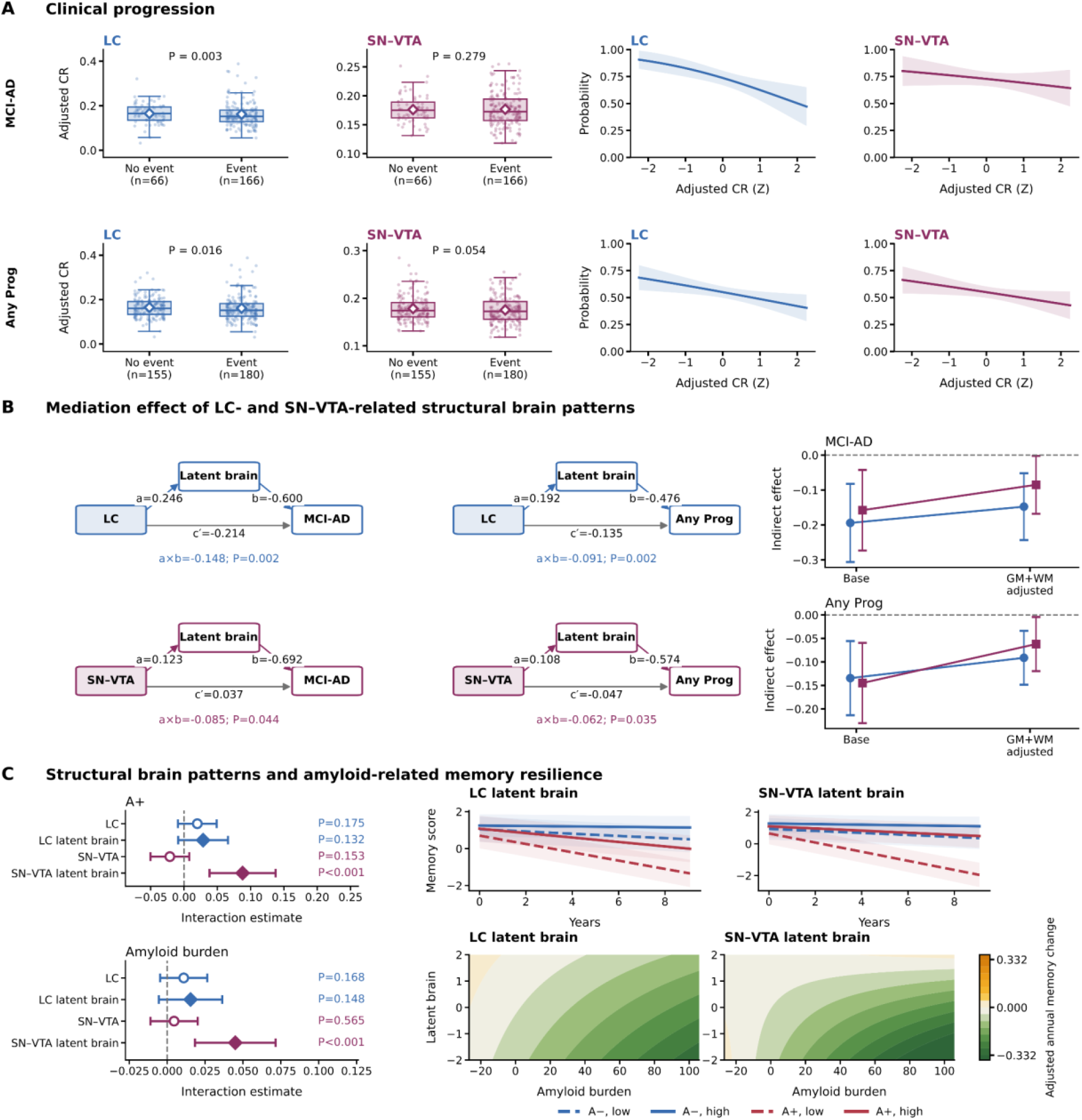
LC and SN-VTA CR in five-year AD progression, distributed structural atrophy pathways and amyloid-related memory resilience. **a**, Five-year clinical progression. Covariate-adjusted bilateral LC or SN-VTA CR is shown according to five-year progression status for MCI to AD (MCI-AD; top row) and any qualifying clinical progression (Any Prog; bottom row). Boxes show the interquartile range, center lines show medians, whiskers extend to 1.5 times the interquartile range, jittered points represent participants, and white diamonds indicate group means. P values are from the corresponding two-sided logistic mixed-effects models. Curves show marginal standardized five-year progression probabilities across standardized LC or SN-VTA CR; shaded bands indicate 95% confidence intervals. **b**, Associational indirect effects through distributed structural patterns. Path diagrams show the total-parenchymal-volume-adjusted models for MCI-AD and Any Prog. The standardized LC or SN-VTA CR is shown on the left; the corresponding PLSC intercept LV1 brain score, labeled “Latent brain” in the figure, is shown above; and progression status is shown on the right. Path a represents the association between LC or SN-VTA CR and the PLSC LV1 brain score; path b represents the association between the brain score and the latent progression response conditional on CR; and c′ is the remaining direct association. The displayed a × b value is the indirect effect. Its displayed P value is unadjusted; P_FDR_ values are reported in the Results. Attenuation plots compare indirect effects estimated in the same sample before and after adjustment for total parenchymal volume, labeled GM+WM in the figure. Points indicate indirect-effect estimates, vertical bars indicate 95% confidence interval. Attenuation percentages were descriptive. **c**, Amyloid-related longitudinal memory resilience. Forest plots show interactions of amyloid burden and time with LC or SN-VTA CR and with the corresponding PLSC LV1 brain score for binary amyloid positivity (A+; upper forest) and continuous Centiloid burden (lower forest) from the total-parenchymal-volume-adjusted models. Open circles denote CR interactions, filled diamonds denote PLSC LV1 brain-score interactions, and horizontal lines indicate 95% confidence intervals. Displayed P values are unadjusted; P_FDR_ values are reported in the Results. Binary-amyloid trajectory plots show fixed-effect predictions for amyloid-negative and amyloid-positive participants at low and high PLSC LV1 brain scores, defined as −1 and +1 SD, respectively. Solid lines indicate high scores, dashed lines indicate low scores, and shaded bands indicate 95% confidence intervals. Continuous-amyloid response surfaces show adjusted fixed-effect memory change over one year across raw Centiloid burden and PLSC LV1 brain scores. The Centiloid axis spans the observed 5th-95th-percentile range, and PLSC LV1 brain scores span −2 to +2 SD. More positive colors indicate more favorable annual memory change. A−, amyloid negative; A+, amyloid positive; Any Prog, any qualifying clinical progression; CR, contrast ratio; FDR, false-discovery rate; GM+WM, total gray- and white-matter volume; LC, locus coeruleus; LV1, latent variable 1; MCI-AD, mild cognitive impairment to AD; PLSC, partial least squares correlation; SD, standard deviation; SN-VTA, substantia nigra-ventral tegmental area; TPV, total parenchymal volume.

## Discussion

Using 3,782 legacy dual-echo MRI scans of 739 unique older participants with up to 20.03 years of follow-up, we derived in-vivo LC and SN-VTA CR measures and examined their distinct association profiles with cognition, regional brain atrophy and clinical progression. Regarding the noradrenergic system, higher LC CR was associated with preserved LC pigmentation at autopsy, better later memory, language and verbal fluency. It was also associated with larger entorhinal, hippocampal volumes, lower WMH burden, and less AD-like pattern of brain atrophy. These LC CR associations were significant over follow-up intervals beginning approximately 5.7-9.2 years after MRI and extending to 17.7-20.0 years, and independent of SN-VTA CR. LC CR was further associated with a distributed brain atrophy pattern, co-localized with the distribution of norepinephrine transporter, that involved insular and medial temporal regions. Evaluating the association between in-vivo catecholaminergic markers and subsequent AD development, each standard-deviation decrease in baseline LC CR was associated with 84% higher five-year odds of MCI-to-AD progression. Regarding the dopaminergic system, higher SN-VTA contrast was associated with better memory, executive function, and everyday functioning, including community activities, hobbies, and transportation, and lower WMH burden. These associations were mainly significant approximately 7-10 years through the latest evaluated time points, independent of LC CR. The everyday-function findings are compatible with dopamine’s role in motivation and effort allocation^86,87^. Individual differences in SN-VTA CR also covaried with a brain-wide atrophy pattern involving the insula, putamen and pallidum, which co-localized with dopamine receptors. Higher scores for this pattern (i.e., less SN-VTA-related atrophy) were associated with less amyloid-related memory decline.

In-vivo imaging of small catecholaminergic nuclei in humans remains challenging. The LC is a narrow structure bordering the fourth ventricle and is represented by only a few voxels at typical in-vivo MRI resolutions, making its estimated signal sensitive to partial-volume mixing, ventricular CSF contamination, small registration errors, and reference-region definition^19,88,89^. SN-VTA measurements are similarly susceptible to partial-volume effects and signal contributions from adjacent midbrain tissue, including paramagnetic-ion-rich nigral regions^69,90,91^. In the multisite ADNI dataset, these anatomical constraints were compounded by variation in field strength, acquisition geometry, echo time, scanner, manufacturer, motion and other image-quality features, which were addressed through quality control, outlier removal and technical adjustment. Despite all of these challenges, our findings indicate that MRI measures of LC and SN-VTA structure carry meaningful information about the future trajectories of cognitive and brain decline.

The physical basis of catecholaminergic MRI contrast also remains debated. Neuromelanin-metal interactions, mobile-water content, and macromolecular properties may contribute differently depending on the acquisition protocol^30,69,92,93^. These uncertainties make empirical qualification of the derived phenotypes essential. The ADNI dataset provides the rare opportunity to compare in-vivo MRI to post-mortem histology, which can help contribute to better understanding of the biological basis of the MRI contrast.

Anatomical localization, repeated scans and autopsy comparisons provided complementary evidence about the catecholaminergic imaging measures. The PD-T2 difference images used to guide localization showed clear bilateral hyperintense signals at the expected LC and SN-VTA locations. ICC measures indicated that between-participant differences in CR were preserved across repeated scans, suggesting temporal stability (Figure 1). Lower LC CR before death was associated with gross LC pigmentation loss at autopsy. Separate tissue analyses linked reduced LC pigmentation to lower neuronal density^94^ and reduced SN pigmentation to α-synuclein/Lewy-body pathology^95^ consistent with prior neuropathologic observations^96,97^. However, gross SN hypopigmentation was not associated with the SN-VTA region combined CR. The weaker SN correspondence may reflect the mismatch between SN-only tissue ratings and an in-vivo mask combining SN and VTA, the larger and more heterogeneous midbrain territory, the high-signal voxel-sampling procedure, small autopsy samples, and biological change between MRI and death.

Older baseline age was associated with lower LC and SN-VTA CR, and both phenotypes showed modest additional longitudinal decline during follow-up. The direction aligns with prior evidence that LC and SN-VTA change becomes more negative with older age and with reported late-life declines in LC and nigral contrast^24,71,98^. Sex was not significantly associated with either measure. Hemispheric differences were also found; previous LC studies indicate that such comparisons can depend on the acquisition and signal-extraction method^88,99^.

It was interesting that despite only weak concurrent associations between the nuclei and cognitive and behavioral measures, LC and SN-VTA contrast substantially predicted future functioning while controlling for baseline levels. Concurrent cognitive associations were largely null, whereas lower LC and SN-VTA CR covaried with selected brain atrophy and everyday-functional measures, including a more AD-like structural brain pattern. Prospective models identified broader associations with subsequent cognition and function after adjustment for baseline outcome level. This temporal difference indicates that catecholaminergic system alterations precede or accompany changes that become measurable only later, although it does not establish a causal relationship.

Within the prospective analyses, higher LC CR was associated with better subsequent memory and language performance, including learning, delayed recall, recognition and category fluency, as well as larger later entorhinal volumes and less clinical impairment. For the memory composite and bilateral entorhinal volumes, higher LC CR was associated with better memory performance or larger volumes over significant follow-up intervals of 7.85-18.80 years for memory, 7.08-18.84 years for left entorhinal volume, and 8.75-17.69 years for right entorhinal volume after MRI. The findings across learning, delayed recall and recognition are consistent with previous human LC imaging results^24^, and may result from a noradrenergic modulation of hippocampal synaptic plasticity and memory^38^. Specifically, experimental rodent studies identified the importance of norepinephrine-sensitive β-adrenergic receptors^38^ and dopamine D1/D5 receptors^37,39^ in LC-dependent hippocampal plasticity and memory. Better performance on the category fluency task depends on controlled retrieval and flexible search, providing a potential link to noradrenergic support of task engagement and memory retrieval^73,100,101^.

Higher dopaminergic imaging measures were associated with better subsequent working-memory and naming performance and with reduced subsequent clinical and functional impairment. The Digit Span Backward finding agrees with prior work relating SN-VTA contrast to working memory and with experimental evidence that VTA dopamine-neuron activity contributes to short-term-memory maintenance^24,102,103^. Associations with complex everyday activities may also be compatible with dopaminergic contributions to motivated, organized, and goal-directed behavior^104^. The lag-dependent analyses indicated that several SN-VTA associations with memory and functional outcomes were most evident at longer follow-up intervals.

Multivariate analyses linked LC and SN-VTA CR levels and rates of change to patterns of regional brain atrophy. The patterns associated with CR levels overlapped in insular and opercular cortex, with additional prominent medial-temporal contributions for LC and striatal and basal ganglia contributions for SN-VTA. These are broadly compatible with known LC connectivity and cortical-basal-ganglia-midbrain organization^61,105,106^. The rate-of-change patterns differed from the intercept patterns, suggesting that stable between-person differences and longitudinal covariation need not share the same topography^107^. The molecular-map associations linking CR-related atrophy to neurotransmitter maps provided complementary spatial context. The LC intercept pattern uniquely corresponded with the cortical noradrenergic transporter map, whereas the SN-VTA pattern uniquely corresponded with D1- and D2-family dopamine-receptor maps. While correlational, these system-concordant associations support the biological substrate of the cortical topographies. The SN-VTA pattern also correlated with several exploratory neurotransmitter maps, but these correspondences were attenuated and no longer significant after joint adjustment for the three dopamine-receptor maps. This attenuation is consistent with shared spatial variation among neurotransmitter systems^63^.

Among participants with MCI at their first MRI assessment, each 1-SD lower LC CR was associated with 84% higher odds of progression to AD and 46% higher odds of any diagnostic progression within five years; both LC associations persisted after accounting for SN-VTA CR, pointing to unique noradrenergic contributions. These findings extend earlier LC MRI studies linking lower contrast to clinical worsening or conversion to dementia^57,58^. SN-VTA CR was not significantly associated with either progression outcome after correction for multiple comparisons. This contrast does not imply that the dopaminergic midbrain was clinically irrelevant - and we note that previous positive studies using different phenotypes, including conventional T1-MRI-derived VTA signal or functional connectivity, found relevant relationships^54,56^.

Higher LC or SN-VTA CR was associated with greater expression of its corresponding structural brain pattern (i.e., lower atrophy), and greater pattern expression in turn was associated with lower clinical progression propensity. All four associational indirect effects remained significant after adjustment for total parenchymal volume, but their magnitudes were attenuated by 24-32% for LC and 47-57% for SN-VTA. Thus, part of each pathway reflected global tissue preservation, especially for the SN-VTA. The direct LC path remained significant in the MCI-to-AD model after adjustment for the LC-related distributed latent brain pattern and total parenchymal volume, whereas the SN-VTA result was expressed principally through the distributed latent brain pattern. Together with the concurrent and prospective associations with cognition and behavior, these observational findings of mediation of prospective findings by structural brain patterns suggest that the biggest impact of declines in LC and SN-VTA structural integrity occurs over time, as the impaired function in these nuclei leads to atrophy in their target cortical brain regions, which in turn impairs cognition and behavior^57,84,108,109^.

Finally, the amyloid analyses further separated catecholaminergic phenotypes from distributed latent brain expression. Amyloid-positive status and higher continuous amyloid burden were associated with more adverse memory trajectories, but neither LC nor SN-VTA CR significantly modified this relationship, and the LC-related structural pattern showed no significant moderation. By contrast, greater expression of the SN-VTA-related pattern (i.e., less atrophy) was associated with less adverse amyloid-related memory decline in both binary and continuous amyloid models after adjustment for SN-VTA CR and total parenchymal volume. At comparable amyloid burden, participants expressing this distributed pattern more strongly showed a less negative memory trajectory. This can be described as resilience-consistent moderation, referring to maintaining better cognitive or functional outcomes in the presence of adverse aging- or disease-related factors^110–112^. Taken together, the findings suggest that clinically relevant catecholaminergic information may be expressed both at the local catecholaminergic system and through distributed brain organization, with LC CR carrying the direct progression signal and the SN-VTA-related structural pattern carrying information about cognitive resilience to amyloid burden.

Several limitations define the scope of these conclusions. The catecholaminergic phenotypes were reconstructed from legacy 1.5-T and 3-T scans acquired across multiple sites, scanner platforms, vendors, and protocols. Although field-strength-specific processing, quality control, image-quality covariates, and technical and site adjustments were used, acquisition heterogeneity likely increased measurement error and may have attenuated the observed associations. Thus, the here reported associations may represent a lower bound of the true effect sizes, replication using a consistent measurement protocol is important. Second, although CSF phosphorylated tau and limited postmortem tau measures were available, regional tau PET was not, precluding direct tests of whether LC variation tracked or preceded spatially specific medial-temporal and cortical tau accumulation, which is important given the roles of the LC in AD-related tau accumulation^1,57,65^. Purpose-designed studies with sufficiently large samples and longitudinal measurements of LC and SN-VTA integrity, whole-brain structure, regional tau PET, and clinical outcomes will be needed in future studies.

Nevertheless, the current study indicates that catecholaminergic-nucleus phenotypes and their related distributed structural volume patterns capture complementary levels of late-life brain and clinical variation. Legacy dual-echo PD-T2 MRI can be used to recover anatomically localized and post-mortem validated LC and SN-VTA CR measures with measurable multi-year stability and modest age- and time-related decline. Their associations were more extensive for subsequent than concurrent cognition and everyday function, with several relationships becoming most evident only after multi-year delays from the catecholaminergic CR assessment. The two neuromodulatory systems showed different association profiles. Lower ante-mortem LC CR was associated with gross postmortem LC hypopigmentation, and a one SD lower LC CR was associated with 84% higher five-year odds of progression from MCI to AD whereas SN-VTA CR was not significantly associated with progression. At the systems level, both phenotypes covaried with distributed structural atrophy patterns whose cortical topographies showed system-concordant correspondence with the norepinephrine-transporter map for LC and D1- and D2-family dopamine-receptor maps for SN-VTA. These catecholaminergic nuclei-associated atrophy patterns carried progression-related information that was not fully summarized by total parenchymal volume, and the SN-VTA-related pattern additionally moderated amyloid-associated memory decline in a resilience-consistent direction. Thus, clinically relevant information may be expressed both in the integrity of LC or SN-VTA and through their related brain systems.

## Data availability

The ADNI data analyzed in this study are available through the LONI Image & Data Archive (https://ida.loni.usc.edu/) following approval of a data-access application and acceptance of the ADNI Data Use Agreement.

## Code availability

Scripts used to conduct the analyses are available at https://github.com/Wenhui-Zhao/adni-lc-snvta/.

## Acknowledgments

We sincerely thank the participants, investigators, and staff of the ADNI for their contributions to data collection and sharing.

## Author contributions

M.M. and M.J.D. designed and supervised the research. W.Z. performed the research and analyzed the data. S.J.G. annotated the postmortem data. H.J.Y. replicated the analytical pipeline. W.Z., K.R., K.C., and A.M. performed image quality control. W.Z., M.M., and M.J.D. wrote the manuscript. W.Z., M.M., M.J.D., H.J.Y., and D.A.N. revised the manuscript.

## Methods

### Study design and data source

Data used in the preparation of this article were obtained from the ADNI database (https://adni.loni.usc.edu/). ADNI is a longitudinal, multisite study initiated in 2004, and detailed recruitment, inclusion, and exclusion criteria have been described previously^32,113^. The present study analyzed matched T1-weighted and dual-echo PD-T2 MRI scan sets together with linked longitudinal phenotypic and biomarker data to quantify LC and SN-VTA CR and evaluate their demographic, clinical, cognitive, biomarker, neuropathologic, and whole-brain structural associations. ADNI study protocols were approved by the institutional review boards of the participating sites. Written informed consent was obtained from participants and/or their authorized representatives, as applicable.

### T1 and dual-echo PD-T2 magnetic resonance images from the ADNI dataset

T1-weighted MRI acquisition protocols in ADNI have been described previously^31^. The ADNI dual-echo fast/turbo spin-echo sequence produces two image contrasts within a single acquisition: a PD-weighted image at the shorter echo time and a T2-weighted image at the longer echo time^31^. Each PD-T2 pair used in the current study was matched to a T1-weighted MRI scan from the same participant acquired within ±2 days and at the same field strength. When multiple T1 scans satisfied these criteria, we retained the scan closest in acquisition date to the PD-T2 scan. T1-weighted images served as the anatomical reference for spatial standardization. The PD-weighted image provided the primary LC- and SN-VTA contrast, in line with prior research showing high intracellular water content for catecholaminergic neurons^29,30^, whereas the paired T2-weighted image was used to support registration and to derive 4^th^ ventricle and brainstem inclusion masks for downstream PD-T2 difference-map processing. Information regarding the sequence parameters for dual-echo fast/turbo spin-echo sequence have been described previously^31,114^.

### Scan inclusion and quality control

To identify images with quality degradation severe enough to compromise LC or SN-VTA measurements, each matched scan set underwent a manual visual quality check using a standardized six-panel montage generated from the PD-weighted image. The first row displayed mid-sagittal, mid-coronal, and mid-axial whole-brain views, whereas the second row displayed pons-level sagittal and coronal views and an enlarged axial view centered on the brainstem. Reviewers assessed both global image integrity and visibility of the target regions, with more attention to severe motion-related blurring, ghosting, zipper or banding artifacts, radiofrequency interference, signal dropout, incomplete anatomical coverage, reconstruction corruption, and artifacts affecting the dorsal pons and fourth-ventricle region or the midbrain SN-VTA region. Mild imperfections were retained when the relevant anatomy remained interpretable. Scans that could not be classified confidently from the montage were inspected in the full Neuroimaging Informatics Technology Initiative (NIfTI) volume using FSLeyes.

The 3,782 dual-echo PD-T2 scans were divided among four trained raters, and each received an initial binary pass-or-fail rating. A random sample of 100 scans was embedded in the routine review batches and rated independently by all four raters to assess interrater agreement. W.Z. additionally performed second-pass adjudication of uncertain or discordant production ratings after discussion with the other raters. An adjudicated rating replaced the corresponding initial production rating and was not treated as an additional vote. Interrater agreement for the 100-scan overlap sample was calculated from the four independent ratings. A scan was classified as failed when at least three of four ratings indicated failure and as passed otherwise. Because failure ratings were infrequent and the binary distribution was strongly imbalanced, Gwet’s AC1 was used as the primary chance-corrected agreement coefficient^115^. Overall pairwise agreement was 89.7% (95% bootstrap CI, 85.2%-93.8%), and Gwet’s AC1 was 0.880 (95% CI, 0.816-0.933). All four raters agreed on the pass-or-fail classification for 82 of the 100 scans, at least three raters agreed for 92 scans, and eight scans had a two-versus-two split.

Following adjudication and application of the prespecified voting rule, 325 of 3,782 scans failed quality control and were excluded (8.6%), whereas 3,457 scans from 731 participants passed and were retained (91.4%). Among the 1.5-T scans, 285 of 3,273 failed (8.7%) and 2,988 passed; among the 3-T scans, 40 of 509 failed (7.9%) and 469 passed. Within the 100-set overlap sample, two matched scans met the three-of-four failure threshold and 98 passed. Matched scans classified as failed were excluded from subsequent LC and SN-VTA feature extraction and statistical analyses. Feature-level extreme-value screening was applied after CR calculation and is described below.

### Diagnosis and covariates

Harmonized demographic and longitudinal diagnostic information was obtained from the Alzheimer’s Disease Sequencing Project Phenotype Harmonization Consortium (ADSP-PHC) Demographics and Diagnosis dataset. The ADSP-PHC maps clinical research diagnoses assigned by the originating cohort to common categories. The harmonized categories were Unaffected, MCI, AD dementia, dementia not attributed to AD, Other Cognitive diagnosis, and Missing; Unaffected was coded as cognitively normal (CN) in the present analyses.

Diagnosis at imaging was defined as the nearest valid diagnosis within ±30 days of the date assigned to the matched MRI scan set. When no valid diagnosis was available within this interval, the nearest valid assessments before and after the MRI scan were identified, and diagnosis was assigned only when the two bracketing classifications agreed. Otherwise, diagnosis at imaging was classified as missing.

Baseline age and baseline diagnosis were taken from the earliest harmonized record designated as the ADNI baseline visit. Sex and years of education were taken from the first harmonized record. Apolipoprotein E (APOE) ε4 allele count was derived from the ADNI APOE genotype results and coded as 0, 1, or 2.

### Phenotypic and biomarker preprocessing

Cognitive, clinical, functional, fluid-biomarker, and derived imaging measures were prepared before linkage to LC and SN-VTA CR measures. Prespecified ADNI numeric missing-value and error codes (−4, −1, −9, −99, −999, −9999, and 999) were recorded as missing. Other continuous analysis variables were standardized using the mean and standard deviation:

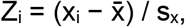

where x_i_ is the data value for observation i, and x̄ and sₓ are the mean and standard deviation, respectively.

Each matched MRI scan set was linked to the nearest assessment of the same participant within ±30 days. The amyloid PET-memory analysis used a separate MRI-PET anchoring procedure due to the available sample size and is described under PET measures and in the corresponding statistical subsection.

### Harmonized cognitive-domain composites

Four harmonized longitudinal cognitive-domain scores, memory, executive function, language, and visuospatial function, were obtained from the ADSP-PHC Cognitive data. The scores were generated by co-calibrating neuropsychological items across studies and test batteries, placing each domain on a common metric^116^. Each score was accompanied by an individual standard error of measurement (SEM). A score was retained only when SEM was ≤0.6, the default precision criterion in the ADSP-PHC documentation. Higher scores indicated better cognitive performance. In ADNI, the harmonized memory score incorporated RAVLT and Logical Memory measures, memory and orientation items from the Alzheimer’s Disease Assessment Scale-Cognitive Subscale (ADAS-Cog) and Mini-Mental State Examination (MMSE), and memory items from the Montreal Cognitive Assessment (MoCA). The executive function incorporated Digit Span, Trail Making, Digit Symbol, clock-drawing components, ADAS-Cog number cancellation, and selected MMSE and MoCA items. Language incorporated Boston Naming, animal and vegetable fluency, and selected language-related items from the ADAS-Cog, MMSE and MoCA. Visuospatial function incorporated clock-copy components, ADAS-Cog constructional praxis and MMSE pentagon copying ^116,117^.

### Test-level and study-derived cognitive measures

Fourteen additional measures were obtained or derived from the ADNI neuropsychological battery. Nine directly recorded outcomes were Rey Auditory Verbal Learning Test (RAVLT) 30-min delayed recall, Digit Span Backward, Digit Symbol, animal and vegetable category fluency, Boston Naming Test total, Logical Memory delayed recall, clock-copy score, and clock-drawing score. Five additional measures were derived from trial-level scores:

RAVLT immediate learning was defined as the sum of correctly recalled words across the five acquisition trials^118^: RAVLT_immediate,i_ = Σ_k=1_^5^ T_ik_,, where Tᵢₖ is the number of words recalled by participant i on acquisition trial k. All five trial scores were required.

The within-assessment RAVLT learning slope was estimated using a regression-based approach^119^:T_ik_ = α_i_ + β_learning,i_k + ε_ik_, k = 1,…,5. The fitted slope coefficient was retained. All five acquisition trials were required; larger positive values indicated greater improvement across successive trials.

RAVLT recognition score was calculated as the original recognition-total field minus the corresponding recognition-error field^120^: R_recognition,i_ = AVDELTOT_i_ − AVDELERR2_i_. Higher values indicated better performance.

To characterize 30-min delayed-recall performance conditional on the acquisition level reached at the end of learning, the following regression was estimated^121,122^: D_30,i_ = α + β T_5,i_ + ε_i_,, where D₃₀,ᵢ is 30-min delayed recall and T₅,ᵢ is Trial 5 recall. The acquisition-adjusted RAVLT delayed-recall score was r_delayed,i_ = D_30,i_ − Ê(D_30,i_ | T_5,i_). Positive values indicated better delayed recall than expected given Trial 5 performance.

To reduce Trail Making Test (TMT) Part B variation shared with Part A, including visual-search, visuomotor, and processing-speed demands, we constructed a regression-adjusted Part B measure. Residual adjustment of Part B for Part A has been described as an alternative to subtraction and ratio scores^123^. Positive Part A and Part B completion times were log-transformed to reduce distributional skewness, and the following regression was estimated:

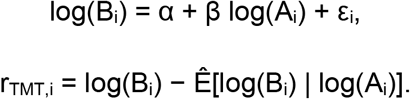

Higher residual values indicated slower Part B completion than expected given Part A performance.

The Trail Making and delayed-recall residual models were estimated in the complete raw dataset before matching to LC and SN-VTA features so that their coefficients did not depend on the composition of a particular imaging-analysis sample. Each model was fitted only when at least 10 complete records and at least three distinct predictor values were available. All 14 test-level or derived measures were standardized before matching with LC and SN-VTA features. Higher values indicated better performance for every measure except the Trail Making residual, for which higher values indicated slower performance.

### Clinical and behavioral measures

Eight clinical and behavioral outcomes were included: the Clinical Dementia Rating (CDR) Judgment and Problem Solving, Community Affairs and Home and Hobbies box scores; the CDR Sum of Boxes; and four Functional Activities Questionnaire (FAQ) items assessing games or hobbies, current events, appointments or medications, and travel or transportation. The CDR box scores measure domain-specific cognitive or functional impairment, whereas the CDR Sum of Boxes summarizes severity across all six CDR domains^124,125^. The FAQ is an informant-rated measure of independence in cognitively demanding instrumental activities of daily living^126^. The four FAQ items were analyzed separately. ADNI raw response codes were mapped to the 0-3 impairment scale: raw codes 0 or 1 were mapped to 0, codes 2 or 3 to 1, code 4 to 2, and code 5 to 3. On the analysis scale, 0 indicated normal independent performance, 1 indicated difficulty with retained independence, 2 indicated a need for assistance, and 3 indicated dependence. Higher values indicated greater impairment^127^.

Within the available ADNI clinical and functional battery, the item-level outcomes were selected a priori to sample distinct clinically relevant domains whose component demands overlap with broad functions modulated by the LC-noradrenergic and dopaminergic midbrain systems, including task engagement and adaptive response control, motivated memory and goal-directed behavior^128–130^.

### AD-related molecular, metabolic, structural, and vascular measures

To characterize associations between LC and SN-VTA CR and AD-related biological processes, the current study used a set of 12 continuous measures selected based on biological coverage and data availability. The variables were organized with reference to the amyloid, tau, and neurodegeneration or neuronal-injury dimensions of the amyloid, tau, and neurodegeneration/neuronal-injury (AT(N)) framework; SPARE brain-age gap was treated as a structural-aging measure, and white-matter hyperintensity (WMH) burden as a separate vascular imaging measure^80^. WMH burden was included because cerebrovascular disease frequently co-occurs with AD and WMHs are commonly interpreted as imaging markers of cerebral small-vessel disease^131,132^. The variable set comprised cerebrospinal fluid amyloid-β42, phosphorylated tau at threonine 181 and total tau; SPARE-AD and SPARE brain-age gap; right and left hippocampal and entorhinal-cortex volumes; fluorodeoxyglucose positron emission tomography (FDG PET) metabolism; amyloid PET Centiloids; and WMH burden.

### Cerebrospinal-fluid biomarkers

Cerebrospinal-fluid (CSF) amyloid-β42, phosphorylated tau181, and total tau were obtained from the harmonized scores from the ADSP Phenotype Harmonization Consortium (ADSP-PHC). The contributing ADNI Elecsys measurements were obtained using fully automated electrochemiluminescence immunoassays at the UPenn/ADNI Biomarker Laboratory^133,134^. Details regarding the lumbar punctures can be found in the ADNI procedures manual (http://www.adni-info.org/).

Higher amyloid-β42 values generally indicate less cerebral amyloid pathology, whereas higher phosphorylated tau and total tau indicate greater tau-related pathology and neuronal injury, respectively.

### Structural MRI summary measures

Six structural summary measures were obtained from the harmonized ADSP-PHC MUSE dataset: SPARE-AD, SPARE brain-age gap, right and left hippocampal volumes, and right and left entorhinal-cortex volumes. In the upstream release, intracranial volume was estimated using DeepMRSeg^135^; T1-weighted images underwent intensity inhomogeneity correction; anatomical regions were segmented with MUSE; longitudinal scans were processed using a pseudo-4D procedure; and segmentations underwent automated and manual quality control. Released MUSE volumes were harmonized using a two-stage ComBat-GAM that modeled scanner- and site-level effects while preserving specified biological covariates^136–138^.

SPARE-AD summarizes expression of a multivariate Alzheimer-like structural atrophy pattern, with higher values indicating a more AD-like pattern^139^. SPARE brain age represents a multivariate estimate of brain age; the brain-age gap was the difference between predicted brain age and chronological age, with higher values indicating an older-appearing brain^140^. The hippocampal measures corresponded to MUSE labels 47 and 48, and the entorhinal measures to labels 116 and 117. Larger values indicated larger regional volumes. The first available ADSP-PHC intracranial volume value for each participant was used as the participant baseline intracranial volume and as a covariate in the MUSE regional-volume analyses described below, including concurrent, prospective and lag-dependent associations and estimation of regional-volume trajectories.

### PET measures

FDG PET metabolism and amyloid burden were evaluated using measures provided by ADNI. The AD-sensitive FDG PET meta-region ratio and global amyloid PET Centiloid value were included in the AD-related variable set. The longitudinal memory-resilience analysis separately used global amyloid status as the primary exposure and the corresponding continuous Centiloid value as the sensitivity exposure.

For each eligible FDG PET scan, the UC Berkeley dataset provided the mean FDG intensity within one AD-sensitive composite meta-region and the mean of the highest 50% of voxel intensities within a pons/cerebellar-vermis reference region. The composite combines the left and right angular gyri, bilateral posterior cingulate gyrus and left and right inferior temporal gyri, which were identified through a literature-based coordinate analysis of FDG PET abnormalities in AD and mild cognitive impairment and combined into one mask^141^. The FDG PET meta-region ratio was calculated as

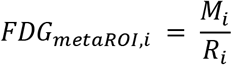

where Mi denotes the mean FDG intensity in the composite meta-region and Ri denotes the mean of the highest 50% of FDG intensities in the pons/cerebellar-vermis reference region. One FDG PET ratio was included per eligible scan; higher values indicated greater relative glucose metabolism in the AD-sensitive cortical composite.

Amyloid burden was represented by the harmonized global Centiloid measure from the ADSP-PHC amyloid PET dataset. In the MRI-free processing framework, amyloid PET images were aligned to standard space, nonlinearly registered to a universal amyloid PET template, summarized within a global cortical target region, and converted from tracer-specific standardized uptake value ratios to Centiloids^142,143^. Centiloids express measurements obtained with eligible amyloid tracers on a common quantitative scale. The AD-related variable set included one continuous global Centiloid value per eligible PET scan.

For the longitudinal memory-resilience analysis, one eligible amyloid PET scan was paired with one LC and SN-VTA MRI session within 90 days. Amyloid status provided by ADSP-PHC was coded as amyloid-negative (A−, 0) or amyloid-positive (A+, 1) using the 25-Centiloid cutoff. A+/A− status was the primary amyloid exposure in separate LC and SN-VTA longitudinal memory models. The corresponding continuous global Centiloid value replaced binary amyloid status in separate sensitivity models.

### White-matter hyperintensity burden

WMH burden was obtained from the combined University of California, Davis ADNI1 and subsequent ADNI releases. Data marked as failed were excluded. The whole-brain WMH volume, expressed in cubic centimeters, was transformed as

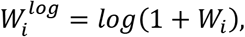

where *W_i_* denotes total WMH volume in the original units. For the WMH measure used in the current study, higher values indicated greater WMH burden.

### Whole-brain MUSE volumes and longitudinal trajectories

The whole-brain analysis used harmonized ADSP-PHC MUSE volume measures. In the ADSP-PHC MUSE processing pipeline, longitudinal scans were processed using a pseudo-four-dimensional procedure that propagated the baseline atlas-registration pathway to follow-up scans, and the resulting segmentations underwent automated and manual quality control. The resulting volumes were subsequently harmonized using a two-stage ComBat-GAM procedure that reduced scanner, site, and field-strength-associated variation. MUSE combines multiple atlases, registration methods, and locally optimized label fusion, and previous evaluations have demonstrated improved segmentation accuracy and interscanner consistency relative to conventional single-atlas approaches^136–138,144^.

The project MUSE dictionary contained 145 directly segmented regions and 114 derived or composite volume measures. The whole-brain PLSC analyses used only the 145 directly segmented regions.

Participant-specific intercept and slope were estimated separately for each MUSE measure and separately for left and right LC or SN-VTA contrast. For participant *i*, MRI session *j*, and MUSE measure *r*, the regional model was

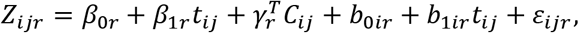

where *Z_ijr_* is standardized regional volume, *t_ij_* is follow-up time in years, *C_ij_* is the fixed-effect covariate vector, *b*_0*ir*_and *b*_1*ir*_are the participant random intercept and random time slope.

MUSE models included participant-baseline intracranial volume (ICV), centered baseline age, sex, centered education, APOE-ε4 allele count, and baseline diagnosis. Because the MUSE volumes had already undergone scanner- and site-level harmonization, no additional batch term or site random intercept was included in these models. The left and right LC and SN-VTA trajectory models used the same participant covariates, omitted ICV, and additionally included field strength, centered entropy focus criterion (EFC), an image-quality measure sensitive to blurring and ghosting^145^, and echo-time category; a site random intercept was retained.

Models were fitted by restricted maximum likelihood using lmer in lme4^146^. Participant-specific parameters were calculated from the fixed effect and participant random effect,

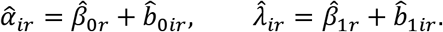

Because time was centered, α̂*_ir_* represents the fitted value at the centered time origin. Intercepts were retained for participants with at least one eligible observation. Slopes were retained only for participants with at least three eligible scans and at least one year of follow-up.

Left and right LC or SN-VTA trajectories were fitted separately. Participants were retained for a bilateral intercept or slope only when the corresponding parameter was available from both hemispheres. Raw side-specific parameters were then averaged,

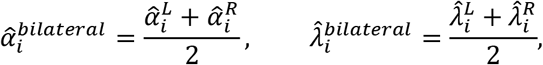

The bilateral parameter was standardized after averaging and then entered into the PLSC analysis.

### Neurotransmitter receptor and transporter maps

Twenty-two PET maps representing 20 molecular targets across 9 neurotransmitter systems were obtained through the neuromaps toolbox and the PET-derived neurotransmitter atlas of Hansen et al.^63,82^. The primary catecholaminergic set comprised two norepinephrine-transporter maps measured with [^11^C]MRB^83,147^, a dopamine-transporter map measured with [¹⁸F]FE-PE2I ^148^, a dopamine D1-like receptor map measured with [^11^C]SCH23390^149^, and two dopamine D2-family receptor maps measured with [^11^C]FLB457 and [¹⁸F]fallypride^150,151^.

The exploratory set comprised maps of serotonin 5-HT_1_A, 5-HT_1_B, 5-HT_2_A, 5-HT_4_ and 5-HT_6_ receptors^152–155^; histamine H_3_ receptors^156^; α4β2 nicotinic acetylcholine receptors, muscarinic M1 receptors and the vesicular acetylcholine transporter^63,157–160^; cannabinoid CB1 receptors^161^; GABA_A_ α5-enriched and broad GABA_A_ benzodiazepine sites^162,163^; κ- and μ-opioid receptors ^164,165^; metabotropic glutamate receptor 5^166^; and NMDA receptors^167^.

The PET maps were parcellated within the cortical gray-matter subset of the fixed primary MUSE atlas for spatial comparison with the PLSC salience maps (see “Partial least squares correlation analysis of whole-brain structural images”). The primary MUSE atlas comprised 145 nonoverlapping anatomical labels, of which 98 were classified as cortical gray-matter parcels; subcortical, cerebellar, white-matter, ventricular and brainstem labels were excluded from the spatial analyses^136^.

### Neuropathologic assessment

Neuropathology data were obtained from the ADNI Neuropathology Core^168^. ADNI neuropathologic assessments follow established diagnostic criteria and are distributed using the National Alzheimer’s Coordinating Center (NACC) Neuropathology Data Forms, Versions 10 and 11. Under the standardized ADNI-NPC sampling protocol, fixed brainstem tissue is sampled axially, including a midbrain block and a pons block containing the SN and LC. Histologic samples are paraffin-embedded, sectioned at 6 μm, and stained with hematoxylin and eosin (H&E) and by immunohistochemistry for β-amyloid, phosphorylated tau, α-synuclein, and phosphorylated TAR DNA-binding protein 43 (TDP-43). The present study used two complementary sources of neuropathologic information: NACC gross hypopigmentation ratings and raw slice-based microscopic annotations of LC-containing pons and SN-containing midbrain sections.

The available data form records the gross LC and SN hypopigmentation on a four-level semiquantitative scale: 0, none; 1, mild; 2, moderate; and 3, severe. The LC-rating sample comprised 27 participants with no hypopigmentation, 18 with mild, 4 with moderate, and 1 with severe hypopigmentation. The SN-rating sample comprised 37 with no hypopigmentation and 17 with mild hypopigmentation; none had moderate or severe ratings. Because moderate and severe categories were sparse, the primary analyses compared no hypopigmentation (rating 0) with any hypopigmentation (ratings 1-3). The LC hypopigmentation analysis included 50 participants with complete model data, including 23 with hypopigmentation present and 27 with hypopigmentation absent. The SN hypopigmentation analysis included 54 participants with complete model data, including 17 with hypopigmentation present and 37 with hypopigmentation absent.

These NACC variables represent macroscopic assessments of regional pigmentation at autopsy. The SN hypopigmentation rating specifically relates to the substantia nigra and does not separately characterize the ventral tegmental area.

Additional quantitative annotations were generated manually from the available LC-containing pons and SN-containing midbrain histologic sections. For each section, the analyzed tissue area was recorded in square millimeters, and H&E-defined neurons, β-amyloid plaques, tau tangles, and α-synuclein-positive Lewy bodies were counted when the corresponding stain was available.

Feature availability differed across stains and regions. In the LC/pons annotation set used for the present analyses, pigmentation ratings were available for 29 participants, amyloid status and log-transformed neuron density for 30, tau status for 29, and α-synuclein/Lewy-body status for 26. In the SN/midbrain annotation set, pigmentation ratings were available for 28 participants, amyloid status and log-transformed neuron density for 32, tau status for 31, and α-synuclein/Lewy-body status for 29. Pairwise correlation sample sizes ranged from 22 to 30 in LC and from 25 to 32 in SN, whereas the complete five-variable PLSC samples included 20 LC participants and 23 SN-VTA participants.

For each pathology family, a binary presence variable was defined as 1 when the lesion count was greater than zero and 0 when the participant was assessed and the count was zero. Neuron density was calculated as neuron count divided by positive annotated area and was natural-log transformed. Associations among the five health-oriented neuropathology measures were estimated separately for LC and SN using pairwise-complete observations. PLSC was conducted separately for LC and SN-VTA using one five-variable annotation block in each region: pigmentation preservation, amyloid absence, tau absence, α-synuclein/Lewy-body absence, and log-transformed neuron density. Global significance was evaluated using 5,000 permutations of the adjusted MRI outcome. Feature stability was evaluated using 5,000 participant-level bootstrap resamples. Supplementary Tables 4-6 report the P values and feature summaries; the global test and bootstrap procedures are detailed in Supplementary Methods.

### MRI group template construction and standardization

MRI processing was performed separately for 1.5T and 3T data, but was otherwise identical unless stated. All image bias field corrections, within-subject registrations, template constructions, and template-to-Montreal Neurological Institute (MNI) registrations were performed with Advanced Normalization Tools (ANTs) version 2.6.2. T1-weighted and T2-weighted images were first corrected for low-frequency intensity non-uniformity using N4BiasFieldCorrection. The N4-corrected T2-weighted image was then affinely registered to the corresponding N4-corrected T1-weighted image using antsRegistrationSyN.sh with affine-only mode (-t a), with T1-weighted MRI as the fixed image. The resulting T2-to-T1 affine transform was subsequently applied to the matched raw PD-weighted image with antsApplyTransforms using linear interpolation, which propagated the PD-weighted image into T1 space while retaining information from the T2-weighted image as the directly registered moving contrast.

Field-strength-specific multivariate group templates were then constructed from the co-registered T1-anchored PD-weighted images. Multivariate template construction was run with bias correction enabled (-b 1), the GR update model (-t GR), cross-correlation similarity (-s CC), -m 50x90x50, -g 0.15, and rigid initialization was enabled (-r 1). The final 1.5T and 3T templates were created by a total of 6 iterations.

The PD-weighted MNI template used as the reference for final group template-to-MNI registration was extracted from Statistical Parametric Mapping version 8 (SPM8) and was resampled to 0.5-mm isotropic resolution with ResampleImage. The final 1.5T and 3T PD-weighted templates were then registered to this 0.5-mm PD-weighted MNI reference with antsRegistration. Registration used winsorization of image intensities to the 0.5th-99.5th percentile, histogram matching, and each PD-weighted template was treated as the moving image and registered to the 0.5-mm PD-weighted MNI template as the fixed image, using rigid, affine and nonlinear stages. The rigid and affine stages used mutual information (32 bins; regular sampling, 0.25), convergence 1000x500x250x100, shrink factors 8x4x2x1, and smoothing sigmas 3x2x1x0vox. The deformable stage used SyN[0.1,3,0] with cross-correlation radius 4, convergence 100x70x50x20, shrink factors 4x2x1x1, and smoothing sigmas 2x1x0.5x0vox. Forward and inverse transforms were retained and concatenated across all transformation steps for subsequent single-shot subject-to-template normalization and inverse warping of LC and SN-VTA priors and reference masks.

### Native LC and SN-VTA priors, reference regions, and anatomical constraints

Left and right LC priors, together with a central pontine reference mask, were taken from the high-confidence LC meta-mask of Dahl et al.^64^ in MNI space and mapped to each subject’s native scan space by concatenating the inverse transforms across the full registration chain (ANTs 2.6.2). Nearest-neighbor interpolation was used to maintain mask boundaries, and the corresponding native-space image was specified as the reference grid. SN-VTA priors and the crus cerebri reference mask were derived from the work of Cassidy et al.^68^ and were mapped to native scan space using the same registration method.

Additional anatomical masks used to constrain LC and SN-VTA processing were derived from whole-brain SynthSeg segmentations of the native T2-weighted images. A brainstem/midbrain inclusion mask was generated from SynthSeg labels 16, 28, and 60 and was used for both LC and SN-VTA. Fourth-ventricle masks were generated from SynthSeg label 15 and were used for LC-specific near-ventricle contamination control. Because the SynthSeg outputs used in this pipeline were on a different grid from the native T2-weighted images, the masks were resampled to the native T2 grid using FMRIB Linear Image Registration Tool (FLIRT) in the FMRIB Software Library (FSL) v6.0.7.10 with nearest-neighbor interpolation.

### PD-T2 MRI difference maps

The dual-echo MRI sequence provided spatially matched PD-weighted and T2-weighted images for each scan. In the acquired ADNI images, the two echoes demonstrated distinct relative signal behavior between CSF and catecholaminergic nuclei. CSF-containing spaces, such as the 4^th^ ventricle, show hyperintensity on the PD-weighted image near the LC and are dark compared to the surrounding tissues on the paired T2-weighted image. This distinction was important because the final PD-based contrast-ratio features summarized the upper portion of the within-mask voxel-wise contrast-ratio distribution. Therefore, if CSF-contaminated voxels were included in the regional mask and had stronger PD contrast than the biological LC or SN-VTA signal, the extracted feature could be dominated by CSF rather than the catecholaminergic-nucleus signal.

We therefore computed voxelwise PD-T2 difference maps in native space as:

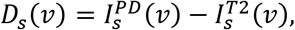

where*D_s_*(*v*) denotes the difference value at voxel v for scan s, and 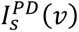 and 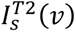 denote the corresponding native-space signal intensities from PD-weighted and T2-weighted images. The difference map was used as an auxiliary image for LC and SN-VTA mask engineering, including contrast-to-noise ratio (CNR)-based mask expansion based on the native LC and SN-VTA priors and CNR-based mask purification. Final contrast-ratio features were extracted from the native PD-weighted image rather than from the PD-T2 difference map.

### LC and SN-VTA mask customization on PD-T2 difference maps

LC and SN-VTA search masks were generated in native space from anatomical priors. Because LC and SN-VTA locations vary across individuals and small registration errors can shift these nuclei relative to atlas-based priors, the PD-T2 difference image was used to refine the spatial support of each prior. Specifically, voxels near the anatomical prior were added when they showed higher PD-T2 difference-image intensity than the corresponding slice-matched regional reference tissue, whereas poorly supported voxels and likely CSF-contaminated voxels were removed in subsequent refinement steps.

For each scan s, region r, and hemisphere h, let *A_s_*_,*r*,ℎ_ denote the native anatomical prior mask. Let *R_s_*_,*r*_ denote the region-specific reference mask, with the central pontine reference mask used for LC and the crus cerebri reference mask used for SN-VTA. Let *B_s_* denote the SynthSeg-derived brainstem and midbrain inclusion mask.

First, the PD-T2 difference image was converted to a slice-referenced scale using the region-specific reference mask. For each axial slice z, finite voxel intensities were extracted from the corresponding slice of the reference mask, denoted *R_s_*_,*r*,*z*_, on the native PD-T2 difference image. These within-slice reference intensities were restricted to values between the 2nd and 98th percentiles to reduce the influence of outlier voxels. If at least 10 finite reference voxels were available on a slice, the mean of the retained values was used as the raw slice-specific reference intensity. Slices with fewer than 10 finite reference voxels were treated as missing. Missing slice-specific reference intensities were filled by linear interpolation across axial slices.

Next, for every reference-mask voxel, the reference mean for that voxel’s slice was subtracted from its PD-T2 difference-image intensity. All resulting residual values from all reference voxels and all slices were combined into a single one-dimensional set. Residuals below the 2nd percentile and above the 98th percentile were excluded, and one standard deviation was calculated from the remaining residual values. Each voxel was then assigned a slice-referenced standardized PD-T2 difference value,

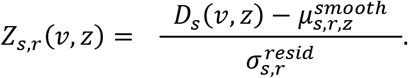

Here, *D_s_*(*v*, *z*) denotes the PD-T2 difference-image intensity at voxel v on slice z; 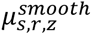 denotes the slice-specific reference mean after interpolation of missing slice means and weighted averaging across the current and immediately neighboring slices; and 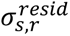 denotes the single standard deviation calculated from the trimmed residual values of all reference voxels across all slices.

The anatomical prior was then expanded within the brainstem/midbrain inclusion mask *B_s_*. A voxel was eligible for addition to the prior only if it satisfied all of the following criteria: it was inside *B_s_*, was within a 1.25-mm in-plane neighborhood of the anatomical prior^89^, was located on an axial slice already occupied by the anatomical prior, and had *Z_s_*_,*r*_(*v*, *z*) ≥ 0.25. Thus, the expansion step permitted local in-plane adjustment around the prior but did not allow the search mask to extend beyond the prior’s original rostrocaudal slice coverage.

After mask expansion, the expanded mask was required to remain spatially close to the anatomical prior. Masks were rejected and replaced by the original anatomical prior if their through-plane centroid shifted by more than 0.75 slices, their in-plane centroid shifted by more than 2.0 mm, any expanded voxels fell outside the prior’s original slice range, fewer than 75% of the prior’s original slices were retained, or fewer than 50% of the prior’s original voxels were retained.

For the SN-VTA mask customization, crus cerebri reference voxels were excluded from the search domain to prevent overlap between the target and reference regions. For the LC mask customization, an additional fourth-ventricle contamination-control step was applied during mask engineering. This step was applied after mask expansion to the provisional LC search mask, so that voxels from both the original LC anatomical prior and the locally added voxels were screened if they were within 1.0 mm of the fourth-ventricle mask. Voxels outside this near-fourth-ventricle zone were not affected. Within the provisional LC search mask, z scores were computed separately for the PD-T2 difference image and the native PD-weighted image using a robust center and scale. Specifically, the center was defined as the median of the finite voxel values within the provisional LC search mask, and the scale was defined as 1.4826 times the median absolute deviation from that median^169^. We used robust z scores because the provisional LC search mask might contain a small number of CSF-contaminated or misregistered high-intensity voxels near the fourth ventricle; median-based scaling reduces the influence of these extreme voxels on the normalization itself, whereas ordinary mean/SD scaling could be biased toward the same outlying voxels that the guard was designed to detect. The resulting robust z scores were used to penalize near-ventricle voxels that showed disproportionately high PD-weighted intensity relative to their PD-T2 difference-image support, because such voxels were more likely to reflect CSF-related signal than LC tissue.

The LC near-ventricular risk score was defined as

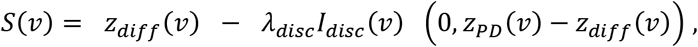

Here, *z_diff_*(*v*) and *z_PD_*(*v*) denote robust within-mask z scores from the PD-T2 difference image and the native PD-weighted image, respectively. *I_disc_*(*v*) indicates near-fourth-ventricle voxels selected for discrepancy penalization. In the primary analysis, *λ_disc_* =0.75, the near-ventricle zone was defined as voxels within 1.0 mm of the fourth ventricle. Near-ventricle voxels with finite *S*(*v*) ≥ 0 were retained. This score is designed for the LC boundary near the fourth ventricle, because true LC-like voxels should be bright on both PD and on the PD-T2 difference image, whereas CSF-contaminated voxels may appear bright on PD but show weaker PD-T2 difference signal. After mask expansion and LC-specific near-ventricle screening, only connected components overlapping the original anatomical prior were retained. The resulting mask was defined as the expanded anatomical search mask, denoted *E_s_*_,*r*,ℎ_.

After mask expansion and LC-specific near-ventricle screening, a ranked voxel-removal step was applied to remove poorly supported voxels from *E_s_*_,*r*,ℎ_. The same slice-referenced PD-T2 value used for mask expansion was used for this removal step. Within *E_s_*_,*r*,ℎ_, voxels with *Z_s_*_,*r*_(*v*, *z*) < 0 were considered poorly supported because their PD-T2 difference-image intensity was lower than the corresponding slice-specific reference baseline. These voxels were ranked for removal. Lower retention priority was assigned to voxels near the boundary of *E_s_*_,*r*,ℎ_, voxels not connected to any non-negative-Z seed through a permissive bridge, and boundary voxels on the first or last occupied slice of the mask. Higher retention priority was assigned to deep interior voxels and to voxels on slices where a large fraction of mask voxels had *Z_s_*_,*r*_(*v*, *z*) < 0, preventing the removal step from aggressively deleting an entire globally low-contrast slice. Voxels with the lowest retention priority were removed first. For the LC, the number of removed voxels was capped by the number of voxels added during local expansion. For the SN-VTA, the primary final mask used the ranked-removal variant capped at 50% of the expanded search-mask voxels. The resulting mask, denoted *M_s_*_,*r*,ℎ_, was used as the final analysis mask for LC and SN-VTA feature extraction.

LC and SN-VTA quantification was performed on native PD-weighted images using the derived analysis masks *M_s_*_,*r*,ℎ_. Feature extraction was performed separately for each scan, region, hemisphere, and field strength. For the LC, the central pontine reference mask was used as the reference region to standardize contrast values for between-participant analyses. For the SN-VTA, the crus cerebri reference mask was used as the reference region. Before feature extraction, each final analysis mask was intersected with the SynthSeg-derived brainstem/midbrain inclusion mask, and voxels belonging to the corresponding reference mask were excluded from the target mask to prevent overlap between target and reference regions. Scans were excluded if the target mask or the corresponding reference mask was empty after these operations (n = 3 after manual QC).

The primary imaging phenotype was a voxel-wise, same-slice, reference-normalized PD CR measure. For each axial slice z, the median PD-weighted intensity was computed within the corresponding slice of the region-specific reference mask *R_s_*_,*r*,*z*_. For each valid target voxel v in the final analysis mask on slice z, the voxel-wise PD contrast ratio was calculated as

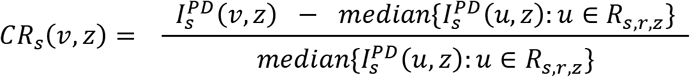

Here, 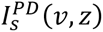 denotes the native PD-weighted intensity at target voxel v on axial slice z, and *R_s_*_,*r*,*z*_denotes the slice-matched reference mask for scan s and region r.

For each hemisphere-specific mask, all valid voxel-wise CR values were ranked, and the feature used in the current study was defined as the mean of the highest 50% of voxel-wise CR values. This emphasis on higher-contrast voxels is consistent with previous studies using peak or high-intensity MRI signals as proxies for catecholaminergic nuclear integrity^24,94^.

Before statistical analysis, CR features underwent an extreme-value screen. Screening was performed separately by region, field strength, and hemisphere. For observation *x_k_*, the robust z score was calculated as

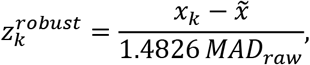

where *x̃* is the median within the corresponding stratum and *MAD_raw_* is the median of the absolute deviations from that median. The scaling factor 1.4826 makes the median absolute deviation consistent for the standard deviation under a Gaussian distribution. Observations with an absolute robust z score greater than 4 were excluded. Median- and median-absolute-deviation-based estimates were used because they are less influenced by extreme observations than their mean- and standard-deviation-based counterparts^170^. After screening, 3,428 scan sets from 726 participants contributed at least one retained hemisphere-specific CR measure, and 3,390 scan sets from 724 participants had bilateral CR measurements for both nuclei.

### Longitudinal repeated-measures stability of LC and SN-VTA CR

Longitudinal repeated-measures stability of CR was evaluated separately for LC and SN-VTA, at each field strength and for each hemisphere and interpreted as a reliability proxy. Because repeated scans were acquired during longitudinal follow-ups likely including true biological change and not in a prespecified short-interval test-retest design, the coefficients were interpreted as measures of longitudinal stability. Within-participant variance in such models could reflect measurement variability, occasion-specific variation, and genuine biological change over follow-up^171,172^. Analyses included matched MRI scan sets retained after manual quality control and feature-level extreme-value screening. Scan sets with AD dementia at imaging or missing diagnosis at imaging were excluded. To limit acquisition-related heterogeneity over follow-ups, only scans with contiguous series preceding the first detected change in PD-image manufacturer, scanner, or protocol were retained. The scan at which the first change occurred and all subsequent scans were excluded.

Participants were required to contribute at least two retained observations for a given LC or SN-VTA, field-strength, and hemisphere combination. For each combination, a participant-random-intercept linear mixed-effects model was fitted by restricted maximum likelihood using lmer in lme4^146^:

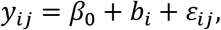

where *y_ij_* is the CR for participant *i* at observation *j*, *b_i_* is the participant-specific random intercept with variance 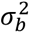, and *ε_ij_* is the within-participant residual with variance 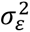. The intraclass correlation coefficient (ICC) was calculated as

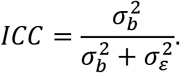

Under this model, the ICC represents the proportion of total observed variance attributable to between-participant differences^172^.

Ninety-five percent confidence intervals were estimated using 5,000 parametric-bootstrap simulations. Confidence limits were defined as the 2.5th and 97.5th percentiles of the finite simulated ICC estimates. Complete ICC estimates are reported in Supplementary Table 2.

### Scan average across 1.5 T and 3 T scans

Group-average images were generated separately for sections including the LC and SN-VTA and for each field-strength to visualize the spatial signal intensity pattern underlying the CR measurements (Fig. 1a). For each native-space scan, the nearest nonempty axial slice to the rounded three-dimensional centroid of the bilateral LC or SN-VTA mask was selected, and a 45 × 45-voxel native-space patch was extracted. Slice-referenced PD-T2 values were first averaged voxelwise within each acquisition geometry. For the 1.5-T images displayed in Fig. 1a, the two geometry-specific averages were combined using their pixel sums and voxel counts on the common 45 × 45-voxel grid with 0.9375 × 0.9375-mm in-plane spacing, the acquisition-geometry-specific averages are provided in Supplementary Fig. 4.

### Statistical analysis

Analyses were performed in R version 4.4.1. The hemisphere was effect-coded as −0.5 for left and +0.5 for right. The EFC, which reflects image entropy associated with motion-related blurring and ghosting, and the foreground-to-background energy ratio (FBER), which calculates the mean energy of the image within the head relative to outside the head, were computed from each T2-weighted image and used as technical image-quality covariates^145,173,174^. Sex, echo-time, and field strength were coded as categorical factor covariates. Sex was modeled with female as the reference category. Echo-time values greater than 1 were converted from milliseconds to seconds, rounded to three decimal places, and modeled categorically. Echo time was modeled categorically in R, so each coefficient represented the CR difference between an echo-time category and the reference category. The category comparisons and reference are reported in Supplementary Table 7.

### Neuropathology associations

LC and SN-VTA CR were adjusted for technical factors separately for each nucleus, field strength and hemisphere using linear mixed-effects models (LME). These models included EFC, FBER, their interaction and echo-time category as fixed effects, and site and participant as random intercepts. The same fitted technical-adjustment models, including FBER and the participant random intercept, were used to prepare CR values for statistical analysis and the corresponding visualization. Adjusted CR values retained participant-specific variation. Left and right values were averaged, and each participant contributed the bilateral value from the MRI acquired closest to death to the neuropathology analyses. Then, four associations were tested: LC CR with LC hypopigmentation, LC CR with substantia nigra hypopigmentation, SN-VTA CR with LC hypopigmentation, and SN-VTA CR with substantia nigra hypopigmentation.

For participant *i*, the model was

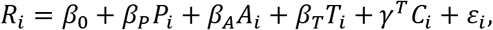

where *R_i_* is the bilateral residual CR, *P_i_* is the binary hypopigmentation, *A_i_* is centered age at death, *T_i_* is the centered MRI-to-death interval, and *C_i_* contains sex, diagnosis at the selected MRI, and field strength. The four associations are reported in Supplementary Table 3.

### Demographic, longitudinal, diagnostic, and hemispheric associations

Associations of CR with baseline age, longitudinal time, sex, and hemisphere were estimated separately for LC and SN-VTA using LME models. For participant i, matched MRI scan set j and hemisphere h, the model for LC or SN-VTA was

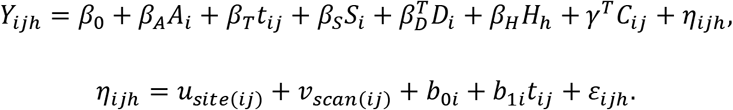

where *Y_ij_*_ℎ_ is the LC or SN-VTA CR; A_i_ is centered baseline age, t_ij_ is centered time from baseline, S_i_ is sex, D_i_ represents baseline diagnosis, H_h_ is hemisphere, and C_ij_ contains field strength, centered education, APOE ε4 allele count, and centered EFC and echo-time category. The model included a site random intercept, a scan-set random intercept to account for the paired hemispheres, and an uncorrelated participant intercept and time slope. Sex and hemisphere contrasts were obtained from estimated marginal means. Full results are reported in Supplementary Table 7. We additionally estimated hemispheric differences separately by scanner manufacturer, both across available field strengths and within each field strength (Supplementary Methods; Supplementary Table 8). To quantify covariance between the two CR measurements, bilateral LC CR was regressed on bilateral SN-VTA CR with the corresponding covariate adjustment (see Supplementary Methods, LC-SN-VTA CR association section).

### Concurrent, prospective and lag-dependent associations with cognitive, clinical and AD-related measures

To determine whether LC or SN-VTA CR was associated with concurrent or subsequent outcomes, 38 cognitive, clinical, behavioral and AD-related biological measures were evaluated using complementary concurrent and prospective mixed-effects models. Concurrent models used assessments obtained within 30 days of the matched MRI scan set. Prospective models related bilateral CR to a later assessment after adjustment for the corresponding baseline value. The 38 measures comprised four harmonized cognitive-domain composites, 14 test-level or study-derived cognitive measures, 12 AD-related molecular, metabolic, structural, and vascular measures, and eight clinical or behavioral measures.

Since LC and SN-VTA demonstrated different CR profiles in the clinically cognitively impaired group (Fig. 5a), and considering cognitive level and longitudinal decline can differ across clinical stage and future progression, including heterogeneity among people with MCI and domain-specific changes approaching AD progression^175,176^, cognitive models used three prespecified diagnosis-stage groups: (1) CN or MCI with no observed diagnostic worsening within available follow-up during the subsequent five years; (2) CN or MCI followed within five years by a higher-stage diagnosis, defined as CN-to-MCI or dementia and MCI-to-dementia worsening; and (3) current AD dementia. All three groups contributed observations. The primary cognitive coefficient was the association in group 1. For AD-related biological and clinical or behavioral measures, one common association coefficient was estimated across the three groups after adjustment for diagnosis stage. Concurrent and prospective models were fitted using LME models.

FDR correction was applied separately for the concurrent and prospective analyses, separately within each of the four outcome families, and separately for LC and SN-VTA^177^. Adjusted *P* values are denoted by *P_FDR_* . t-values were used for display because the concurrent and prospective models had different dependent variables and outcome scales. For visualization only, t-value signs were reversed for measures where lower values indicate better outcomes, so that orange indicates an association of higher CR with better outcomes and green the opposite association. Numerical reports retain the original model signs. Asterisks indicate P_FDR_ < 0.05; hollow circles indicate nominal P < 0.05 but P_FDR_ ≥ 0.05.

### Concurrent associations

Concurrent models related a phenotype or biomarker obtained within 30 days of MRI to the hemisphere-specific LC or SN-VTA CR from the matched MRI scan set.

For the cognitive composites and test-level or study-derived cognitive measures, the phenotype-CR association was allowed to vary across the three diagnosis-stage groups:

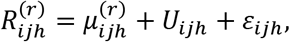

with

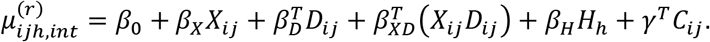

For the AD-related biological and clinical or behavioral measures, diagnosis stage was retained as an adjustment factor, but the phenotype-CR association was constrained to be common across stages:

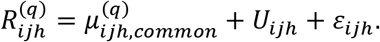

with

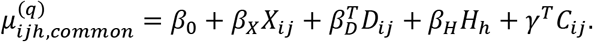

Here, i indexes participants, j indexes matched MRI scan sets, h indexes hemisphere, and q denotes LC or SN-VTA. R_ijh_^q^ is hemisphere-specific CR, X_ij_ is the phenotype or biomarker, D_ij_ is the diagnosis-stage vector, and H_h_ is the effect-coded hemisphere term. C_ij_ contains centered baseline age, centered time from imaging baseline, sex, field strength, centered education, APOE ε4 allele count, centered EFC, and echo-time category. Participant-baseline intracranial volume was additionally included for MUSE structural-volume measures.

In the cognitive interaction model, β_X_ is the phenotype-CR association in group 1, and β_XD_ contains the slope differences for the diagnostic-worsening and current-AD groups; β_X_ was therefore the primary cognitive coefficient in Fig. 3a. In the model without interaction terms, β_X_ is the common association across diagnosis stages and so serves as the primary coefficient for AD-related biological and clinical or behavioral measures. Complete results for the primary concurrent associations are provided in Supplementary Table 9.

The concurrent random-effects component was

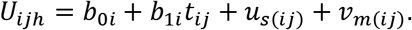

where b_0i_ and b_1i_ are the uncorrelated participant random intercept and time slope, u_s_(ij) is the site random intercept, and v_m_(ij) is the matched-scan-set random intercept that accounts for paired hemispheric observations. ε_ijh_ is the residual error.

### Prospective associations

Prospective models related the bilateral mean LC or SN-VTA CR to a later value of the phenotype or biomarker. The baseline assessment was linked to MRI within 30 days. A future assessment was eligible when it was distinct from the baseline assessment, occurred after both MRI and the baseline assessment, and was obtained 180-1,095 days after MRI. When multiple assessments were eligible, the one closest to 365 days was retained. Bilateral CR was standardized separately for each nucleus and outcome within each MRI field strength, by subtracting the corresponding mean and dividing by the corresponding standard deviation.

For each cognitive outcome, the same model included its baseline value and LC or SN-VTA CR as predictors of the later value. Interactions with diagnosis-stage group allowed the associations of both predictors with the later outcome to differ across groups:

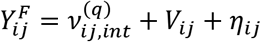

with

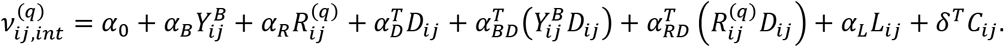

For the AD-related biological and clinical or behavioral outcomes, diagnosis stage was included as an adjustment factor without interaction terms:

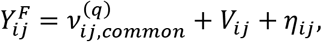

with

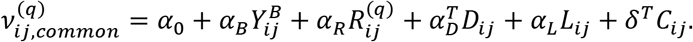

Y_ij_^B^ and Y_ij_^F^ are the baseline and future outcome values, respectively; R_ij_^q^ is the bilateral LC or SN-VTA CR; L_ij_ is the centered interval between MRI and the future assessment; D_ij_ is the diagnosis-stage; and C_ij_ contains the concurrent-model covariates other than hemisphere. Participant-baseline intracranial volume was included for MUSE structural-volume outcomes.

In the cognitive interaction model, α_R_ is the association between CR and the future cognitive outcome in group 1 after adjustment for baseline cognition, and α_RD_ contains the corresponding slope differences for the two nonreference groups; α_R_ was the primary cognitive coefficient in Fig. 3b. In the model without interaction terms, α_R_ is the common prospective association across diagnosis stages after adjustment for the baseline outcome and diagnosis stage. Complete focal prospective results are provided in Supplementary Table 11.

The prospective random-effects component was

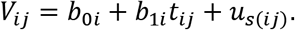

where b_0i_ and b_1i_ are the uncorrelated participant random intercept and time slope and *u_s_*_(_*_ij_*_)_ is the site random intercept. A scan-set random intercept was not included because each matched MRI scan set contributed one bilateral CR value. η_ij_ is the residual error. Sensitivity analyses assessed dependence on response direction, the prospective assessment window, inclusion of both CR measures, and restriction to CN/MCI at MRI. (Details in Supplementary Methods, Concurrent and prospective sensitivity analyses and Analyses restricted to CN and MCI at scan; Supplementary Figs. 8-9; Supplementary Tables 9-11).

### Lag-dependent generalized additive model

All 38 phenotype and biomarker outcomes were examined using lag-dependent generalized additive models (GAMs) that included all eligible post-MRI assessments. Figure 3c shows the LC and SN-VTA curves for the 18 outcomes with the highest significant-interval coverage. For each nucleus and outcome, coverage was calculated as the total duration of intervals where the confidence band excluded zero and each interval lasted at least 20% of the full observed lag range, divided by that range. Outcomes were ranked by the higher coverage value of LC or SN-VTA. The remaining 20 outcomes are shown in Supplementary Figs. 10-11. These follow-up models characterized whether an association was early, delayed, sustained, or changed direction over longer follow-up durations. Observations with current AD dementia were excluded to focus on pre-dementia trajectories, given the nonlinear and stage-dependent evolution of cognition, brain structure, and AD-related biomarkers across the disease continuum^175,178,179^.

For future assessment k occurring *l_ijk_* years after MRI, the centered lag was defined as 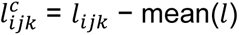. Before fitting the lag-dependent models, bilateral CR was standardized separately for each nucleus and outcome within each MRI field strength. Separate models were fitted for LC and SN-VTA:

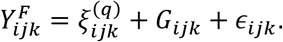

with

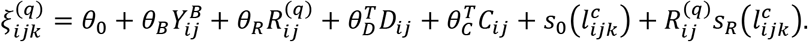

Y_ijk_^F^ is the future outcome, Y_ij_^B^ is the baseline outcome linked to MRI, R_ij_^q^ is bilateral LC or SN-VTA CR, D_ij_ distinguishes group 1 from subsequent diagnostic worsening, and C_ij_ contains the prospective-model covariates; participant-baseline intracranial volume was included for MUSE structural-volume outcomes. The smooth s_0_(*l^c^*) represents the average nonlinear association between elapsed time and the outcome at mean CR. The varying-coefficient term R ^q^s (*l^c^*) allows the CR association to vary with elapsed time.

The lag-specific association of LC or SN-VTA CR was therefore:

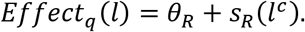

The varying-coefficient smooth was centered to zero at mean follow-up, so θ_R_ is the CR association at the reference lag and s_R_(*l^c^*) is its lag-specific deviation.

The random-effects component was

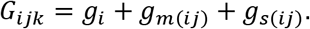

where *g_i_*, *g_m_*_(*ij*)_ and *g_s_*_(*ij*)_ are participant, matched-scan-set and site random intercepts implemented as random-effect smooths; ε_ijk_ is the residual error.

Models were fitted using mgcv::bam with fast restricted maximum likelihood, discrete computation and thin-plate regression splines with fixed basis dimension k = 5^180,181^. Curves were evaluated at 201 points spanning the full observed follow-up interval for each outcome and LC or SN-VTA model. Curve-specific 95% simultaneous confidence bands were estimated from 10,000 simulations using the unconditional Bayesian covariance matrix^182–184^. Shaded intervals identify lag ranges over which the corresponding band excludes zero. For measures where lower values indicate better outcomes, y-axes were reversed for display. Sensitivity analyses examined the effects of follow-up interval restrictions, limiting the sample to CN/MCI participants, and adjusting for the other CR measure using additional lag models (Supplementary Methods, “Lag-dependent sensitivity analyses”; Supplementary Figs. 12-23; Supplementary Tables 12-13). To exclude extreme follow-up intervals from display, the model curves were also evaluated over the 5th-95th percentile range of the lags (Supplementary Figs. 24-27).

### Partial least squares correlation analysis of whole-brain structural images

PLSC^185^ was used to identify multivariate patterns of covariation between LC or SN-VTA trajectories and trajectories of 145 whole-brain regional volumes segmented from the MUSE atlas.

Participant-specific MUSE parameters were obtained from linear mixed-effects models fitted separately to each harmonized MUSE volume. The models included centered follow-up time, participant-baseline intracranial volume, centered baseline age, sex, centered education, APOE ε4 allele count, and baseline diagnosis, together with uncorrelated participant random intercepts and time slopes. Left and right LC and SN-VTA trajectories were estimated separately. These models included centered follow-up time, centered baseline age, sex, centered education, APOE ε4 allele count, baseline diagnosis, field strength, centered EFC and echo-time category, together with uncorrelated participant random intercepts and time slopes and a site random intercept. Raw left- and right-hemisphere participant-specific parameters were averaged after model fitting, and the bilateral intercept or rate-of-change parameter was then standardized across participants. Rates of change required at least three eligible scan sets, at least two distinct time values, and at least one year of follow-up in each hemisphere.

For each primary analysis, the participant-by-parcel matrix X contained MUSE intercepts or rates of change for the 145 parcels, and the vector y contained the corresponding bilateral LC or SN-VTA parameter. PLSC decomposed the cross-block covariance as

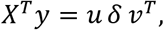

where u contains the MUSE parcel saliences, v is the LC or SN-VTA salience, and δ is the latent-variable singular value. Because the LC or SN-VTA block contained one bilateral parameter, each direct analysis yielded one latent variable (LV1). Participant MUSE brain scores and LC or SN-VTA scores were calculated as

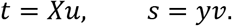

Latent-variable significance was assessed using 5,000 permutations. Parcel-salience stability was estimated from 5,000 participant bootstrap resamples and summarized by the bootstrap ratio of the scaled salience

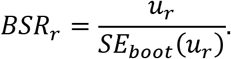

Permutation testing evaluates the complete latent covariance pattern, whereas bootstrap ratios describe the stability of individual parcel contributions^185^.

Latent-variable orientation was fixed independently of the molecular PET analyses. For each direct analysis, the LC or SN-VTA salience was oriented to be positive, and the same orientation factor was applied to regional saliences, bootstrap ratios, and participant scores. Positive regional saliences therefore indicate that a higher regional-volume intercept or a more positive regional-volume rate of change covaried with a higher LC or SN-VTA intercept or rate of change; negative saliences indicate inverse covariation. Brain maps display continuous, saliences with |BSR| ≥ 2.56. At most ten eligible anatomical names were displayed, including at most four subcortical names, with unused places filled by eligible cortical names. All 145 oriented weights were retained for participant scores and subsequent analyses.

Sensitivity analyses examined LC- and SN-VTA-specific covariance patterns after accounting for the corresponding parameter of the other region. For the unique analyses, LC and SN-VTA parameters were standardized on the same complete participant set and residualized against one another separately for intercept and rate of change:

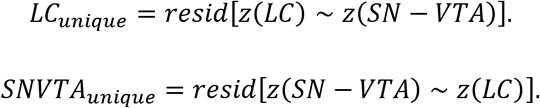

The residualized parameters were then entered into otherwise identical PLSC analyses. Molecular correspondence of the unique intercept patterns was evaluated to assess whether the primary spatial findings persisted after accounting for covariance between the two CR parameters (see Supplementary Methods, Unique structural patterns and molecular correspondence section; Supplementary Fig. 29; Supplementary Tables 15-17).

### Spatial correspondence with neurotransmitter receptor and transporter maps

Spatial correspondence analyses used the continuous, unthresholded cortical saliences from the LC and SN-VTA intercept LV1s. Twenty-two group-average PET maps, comprising six primary catecholaminergic maps and 16 exploratory receptor, transporter, or binding-site maps, were obtained through neuromaps and the PET-derived neurotransmitter atlas and summarized within the cortical gray-matter parcels of the same MUSE atlas used for PLSC.

For PLSC map *m* and PET map *k*, spatial correspondence was quantified using Spearman’s rank correlation:

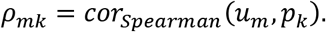

For each PLSC map, 10,000 Moran-spectral surrogate maps were generated from the Euclidean distance matrix between MUSE parcel centroids using a Gaussian spatial-weight kernel and ten nearest neighbors^186,187^.

The two-sided empirical spatial probability was calculated as

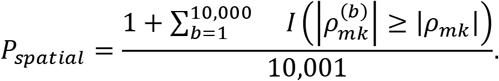

FDR correction was applied separately within each PLSC map and PET set. The intercept analysis therefore comprised four correction families: LC-primary, LC-exploratory, SN-VTA-primary and SN-VTA-exploratory. As a sensitivity analysis, we calculated partial Spearman correlations for the 16 exploratory PET maps, controlling for LC correspondences for the Hesse MRB NET map and SN-VTA correspondences for the SCH23390, FLB457, and fallypride dopamine-receptor maps separately and jointly. Procedural details are provided in Supplementary Methods. Complete results are reported in Supplementary Table 17; additional details are given in Supplementary Methods.

### Five-year clinical progression

Associations of LC and SN-VTA CR with five-year clinical progression were evaluated using logistic mixed-effects models. The index MRI set was the first eligible LC- or SN-VTA CR for each participant. The exposure was the standardized bilateral technically adjusted CR described under Neuropathologic associations. Technical adjustment was performed separately for LC and SN-VTA, field strength, and hemisphere; the left- and right-hemisphere adjusted values were then averaged. Participant-specific variation retained during technical adjustment was not removed since we only used baseline scans. For each nucleus, the CR was standardized by subtracting its mean and dividing by its SD across all eligible MRI examinations at both field strengths, including repeated examinations. These means and SDs were calculated before selecting the progression samples and were used for both progression outcomes.

Two outcomes were evaluated. The MCI-to-AD analysis included participants with MCI at the LC-SN-VTA baseline scan and defined progression as a subsequent diagnosis of AD. The any-progression analysis included participants who were CN or had MCI at the MRI LC-SN-VTA scan and defined progression as CN to MCI or dementia, or MCI to AD. Qualifying transitions had to occur more than 30 days and no later than five years after that MRI. Participants without an event were required to have at least five years of diagnostic follow-up. For participant i at site s, the model was

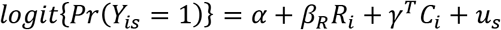

where *Y_is_* is the five-year progression indicator, *R_i_* is the residual CR, *C_i_* is the covariate vector, and *u_s_* is a site random intercept. The site random intercept was included because the logit model is nonlinear and it may still change the coefficient of interest or its uncertainty. For the covariates, MCI-to-AD models adjusted for centered age at MRI, sex, centered years of education, APOE-*ε*4 allele count, and field strength. The any-progression models additionally adjusted for diagnosis at the earliest eligible scan.

Models were fitted with a binomial distribution and logit link using lme4::glmer^146^. For the primary separate LC and SN-VTA models, FDR correction was applied across the two nuclei within each progression endpoint.

Progression-probability curves were generated from the fitted fixed effects. At each CR value, predicted probabilities were averaged over the observed covariate distribution to obtain marginal standardized five-year progression probabilities and 95% confidence intervals. Both CR measurements were included to assess their conditional associations with progression (see Supplementary Methods, Clinical progression with both LC and SN-VTA CR section; Supplementary Fig. 31; Supplementary Tables 18 and 19).

### Associational indirect effects through distributed structural brain patterns

Mediation analyses evaluated whether the statistical association between LC or SN-VTA CR and five-year progression included a pathway through the corresponding oriented PLSC intercept LV1 brain score. Higher brain scores indicate greater expression of the whole-brain volume pattern that covaried positively with LC or SN-VTA CR.

The total-parenchymal-volume-adjusted model included the intercept of total parenchymal volume to assess whether the indirect associations persisted after accounting for overall brain volume, defined as the sum of harmonized total gray- and white-matter volumes. For LC and SN-VTA separately, the two-level model was

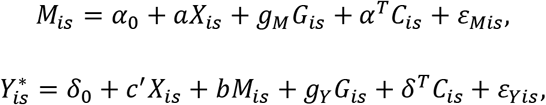

where *X_is_* is LC or SN-VTA CR, *M_is_* is the corresponding distributed structural score, *G_is_* is total parenchymal volume, and 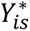 is the latent probit-scale propensity for progression. The observed progression outcome equaled 1 when 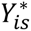 exceeded the estimated threshold and 0 otherwise. The base model omitted *G_is_* from both equations. Covariates matched those used in the corresponding progression model. Models were fitted in lavaan 0.7-2 as two-level structural equation models^188^. The defined effects were:

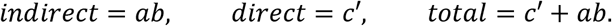

FDR correction was applied across the LC and SN-VTA indirect effects within each endpoint and model variant. Attenuation after total-parenchymal-volume adjustment was summarized descriptively as:

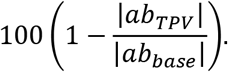

Three sensitivity analyses assessed dependence on the PLSC training population and models with temporal orders. Detailed definitions are provided in Supplementary Methods (Mediation using CN/MCI-trained PLSC patterns and Mediation analyses with temporal order sections; Supplementary Tables 19, 21 and 22).

### Amyloid-related longitudinal memory resilience

Amyloid-related moderation of longitudinal memory decline was evaluated among participants with CN or MCI status at baseline. Amyloid PET eligibility, quality control, and 90-day MRI-PET pairing followed the procedures described above. The outcome was the harmonized memory composite from ADSP-PHC. Participants were required to contribute at least two retained memory assessments spanning at least 180 days.

Binary amyloid positivity status was the primary pathology measure. Dependence on this binary classification was assessed by repeating the complete models with continuous Centiloid burden. For each amyloid measure, separate LC and SN-VTA models included three potential modifiers of the amyloid-associated memory trajectory: LC or SN-VTA CR, the corresponding PLSC intercept LV1 brain score, and total parenchymal volume. For participant i, memory assessment j, site s, and LC or SN-VTA model q, the primary total-parenchymal-volume-adjusted model was:

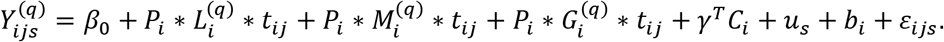

Y_ijs_^q^ is memory performance; P_i_ is binary amyloid status or Centiloid burden; L ^q^, M ^q^ and G ^q^ are LC or SN-VTA CR, PLSC LV1 brain score and total parenchymal volume; and t_ij_ is centered time from the scan. In the formula, * denotes inclusion of all constituent main effects, two-way interactions, and the corresponding three-way interaction. C_i_ contains centered baseline age, sex, diagnosis at index, centered education, APOE ε4 allele count, centered EFC and echo-time category; u_s_ and b_i_ are site and participant random intercepts. Field strength was omitted because all matched MRI scans were acquired at 1.5 T. Models were fitted using LME.

The prespecified coefficients of interest were the amyloid-by-CR-by-time and amyloid-by-PLSC-score-by-time interactions. FDR correction was applied separately for binary amyloid status and continuous Centiloids and separately for LC and SN-VTA CR and PLSC-score interactions.

A positive amyloid-by-brain-measure-by-time coefficient was described as resilience-consistent moderation only when amyloid burden was associated with a more adverse memory trajectory at the reference levels of the modifiers^110–112^.

Binary-amyloid trajectory plots were generated from the fixed effects of the fitted models at low and high PLSC LV-pattern expression, defined as −1 and +1 SD, respectively. Continuous-amyloid response surfaces varied raw Centiloid burden over its observed 5th-95th-percentile range and PLSC-pattern expression from −2 to +2 SD. Surface values represent model-estimated annual memory change, using all longitudinal memory assessments. Complete interactions are reported in Supplementary Table 23, and CR-specific predictions are illustrated in Supplementary Fig. 34. Additional definitions of the participant/reference scales are given in Supplementary Methods.

