## Supplementary Information for "In-vivo MRI of noradrenergic and dopaminergic nuclei predicts future cognition and Alzheimer’s disease progression"

**Contents**


Supplementary Methods

Supplementary Note

Supplementary Figure 1. Scan-processing flowchart

Supplementary Figure 2. Left-right LC and SN-VTA CR distributions by field strength

Supplementary Figure 3. LC and SN-VTA CR correspondence between 1.5 T and 3 T

Supplementary Figure 4. Acquisition-geometry-specific averaged LC and SN-VTA images

Supplementary Figure 5. Neuropathological associations with LC and SN-VTA CR

Supplementary Figure 6. In vivo MRI and postmortem brainstem histology from the same participants

Supplementary Figure 7. LC-SN-VTA CR association

Supplementary Figure 8. Concurrent and prospective LC and SN-VTA model comparisons

Supplementary Figure 9. Prospective associations across alternative follow-up windows

Supplementary Figure 10. Lag-dependent associations with additional cognitive and clinical measures

Supplementary Figure 11. Lag-dependent associations with additional functional, molecular and brain-imaging measures

Supplementary Figure 12. Lag-dependent associations from 180 days after MRI with cognitive composites and cognitive tests

Supplementary Figure 13. Lag-dependent associations from 180 days after MRI with additional cognitive, clinical and functional measures

Supplementary Figure 14. Lag-dependent associations from 180 days after MRI with clinical, functional and biological measures

Supplementary Figure 15. Lag-dependent associations from 180 days after MRI with amyloid PET and white matter hyperintensity burden

Supplementary Figure 16. Lag-dependent associations with cognitive composites and cognitive tests in participants with CN or MCI at MRI

Supplementary Figure 17. Lag-dependent associations with additional cognitive, clinical and functional measures in participants with CN or MCI at MRI

Supplementary Figure 18. Lag-dependent associations with clinical, functional and biological measures in participants with CN or MCI at MRI

Supplementary Figure 19. Lag-dependent associations with amyloid PET and white matter hyperintensity burden in participants with CN or MCI at MRI

Supplementary Figure 20. Mutually adjusted LC and SN-VTA lag-dependent associations with cognitive composites and cognitive tests

Supplementary Figure 21. Mutually adjusted LC and SN-VTA lag-dependent associations with additional cognitive, clinical and functional measures

Supplementary Figure 22. Mutually adjusted LC and SN-VTA lag-dependent associations with clinical, functional and biological measures

Supplementary Figure 23. Mutually adjusted LC and SN-VTA lag-dependent associations with amyloid PET and white matter hyperintensity burden

Supplementary Figure 24. Lag models over 5th-95th percentile lag range: cognitive composites and cognitive tests
Supplementary Figure 25. Lag models over 5th-95th percentile lag range: additional cognitive, clinical and functional measures
Supplementary Figure 26. Lag models over 5th-95th percentile lag range: clinical, functional and biological measures
Supplementary Figure 27. Lag models over 5th-95th percentile lag range: amyloid PET and white matter hyperintensity burden
Supplementary Figure 28. Direct PLSC saliences and bootstrap ratios at |BSR| ≥ 2.56
Supplementary Figure 29. Unique structural patterns and molecular correspondence
Supplementary Figure 30. Exploratory PET correspondence of intercept patterns
Supplementary Figure 31. Clinical progression with LC and SN-VTA included in the same model
Supplementary Figure 32. Mediation effects for five-year clinical progression
Supplementary Figure 33. Mediation pathways for five-year clinical progression
Supplementary Figure 34. LC and SN-VTA CR interactions with amyloid-related memory change
Supplementary Table 1. Sample characteristics
Supplementary Table 2. ICC
Supplementary Table 3. Associations of LC and SN-VTA CR with neuropathological hypopigmentation
Supplementary Table 4. Pairwise associations among neuropathological measures
Supplementary Table 5. PLSC associations of LC and SN-VTA CR with neuropathology
Supplementary Table 6. PLSC loadings for neuropathological measures
Supplementary Table 7. Demographic associations of LC and SN-VTA CR
Supplementary Table 8. Hemispheric differences by scanner manufacturer and field strength
Supplementary Table 9. Concurrent associations of LC and SN-VTA CR with cognitive, clinical and AD-related measures
Supplementary Table 10. Agreement of concurrent and prospective CR coefficients across model specifications
Supplementary Table 11. Prospective associations of LC and SN-VTA CR with cognitive, clinical and AD-related measures
Supplementary Table 12. Lag-dependent association intervals for LC and SN-VTA CR
Supplementary Table 13. Summary of lag-dependent association intervals
Supplementary Table 14. Complete regional LV1 saliences and bootstrap ratios
Supplementary Table 15. PLSC analyses
Supplementary Table 16. Unique structural PLSC and primary molecular-correspondence results
Supplementary Table 17. Molecular PET correspondence
Supplementary Table 18. Clinical progression
Supplementary Table 19. Clinical progression, mediation and continuous-amyloid memory sensitivities
Supplementary Table 20. Primary mediation of five-year clinical progression through longitudinal brain volume patterns
Supplementary Table 21. Mediation using PLSC patterns estimated in CN and MCI participants

Supplementary Table 22. Temporal-order mediation sensitivity: T1 after LC/SN-VTA MRI
Supplementary Table 23. Amyloid-memory interaction

### Supplementary Methods

#### Neuropathology associations

The five neuropathological measures have been described in the Methods (Neuropathologic assessment section). Pairwise associations were assessed using phi coefficients and Fisher's exact tests for binary pairs and point-biserial Pearson correlations for binary-continuous pairs. In the PLSC analyses, one adjusted CR measure was related to the five neuropathological measures. Global PLSC P values were obtained by permutation testing. P values were presented in Supplementary Tables 4, 5. Loadings for the neuropathological measures were reported in Supplementary Table 6 and Supplementary Fig. 5.

#### LC-SN-VTA CR association

To assess covariance between the two CR measurements, using the same data from the Demographic, longitudinal, diagnostic, and hemispheric associations analysis, the following model was fitted:

$$R_{ij}^{LC} = \beta_0 + \beta_R Z_{ij}^{SN-VTA} + \gamma^T C_{ij}^{dem} + b_{0i} + b_{1i} t_{ij} + u_{s(ij)} + \varepsilon_{ij}.$$

Participant and scan were indexed by $i$ and $j$, respectively. $R^{LC}$ denoted raw bilateral LC CR, and $Z^{SN-VTA}$ denoted bilateral SN-VTA CR. $\beta_0$ was the intercept and $\beta_R$ the adjusted LC CR difference per SD of SN-VTA CR. $C^{dem}$ contained the demographic model's fixed covariates except hemisphere: baseline age, imaging time, sex, baseline diagnosis, field strength, education, APOE ε4 count, EFC, and echo-time category. Uncorrelated participant intercepts and imaging-time slopes were represented by $b_{0i}$ and $b_{1i}$; $u_{s(ij)}$ was the site intercept and $\varepsilon_{ij}$ the residual error (Supplementary Fig. 7).

#### Correlation between 1.5 T and 3 T CRs

To test correspondence between 1.5 T and 3 T CRs, MRI scans were paired within participants at the same site and within 30 days. Separate models were fitted for LC and SN-VTA using 176 paired scans from 86 participants:

$$R_{ij}^{q,\,3\,T} = \beta_0 + \beta_R Z_{ij}^{q,\,1.5\,T} + \gamma^T C_{ij}^{pair} + b_{0i} + b_{1i} t_{ij} + u_{s(ij)} + \varepsilon_{ij}.$$

Participant and paired scans were indexed by $i$ and $j$, respectively, and $q$ denoted LC or SN-VTA. $R^{q,\,3\,T}$ denoted raw bilateral CR at 3 T, and $Z^{q,\,1.5\,T}$ denoted bilateral CR at 1.5 T standardized within the complete paired sample. $\beta_0$ was the intercept and $\beta_R$ the correspondence effect of interest. $C^{pair}$ contained the demographic model's fixed covariates except hemisphere and field strength: baseline age, imaging time, sex, baseline diagnosis, education, APOE ε4 count, and EFC and echo-time category from each paired scan. Imaging time $t_{ij}$ corresponded to the 3-T scan. Uncorrelated participant intercepts and imaging-time slopes were represented by $b_{0i}$ and $b_{1i}$; $u_{s(ij)}$ was the site intercept and $\varepsilon_{ij}$ the residual error. Results for LC and SN-VTA are shown in Supplementary Fig. 3a,b, respectively.

#### Hemispheric differences by scanner manufacturer and field strength

To examine hemispheric differences within acquisition groups, we fitted separate mixed-effects models for GE, Siemens and Philips scanners. Analyses were conducted across available field strengths and separately at 1.5 T and 3 T. We retained the covariates and random-effects structure of the primary demographic models, except that field strength and echo-time terms were omitted when constant within a group. (Supplementary Table 8).

#### Concurrent and prospective sensitivity analyses

Alternative follow-up windows were evaluated using the same settings as the primary prospective model (see Methods, Prospective associations section). Only the eligibility window was changed: future assessments were required to occur at least 180 days or at least 90 days after the LC/SN-VTA MRI, without an upper limit (Supplementary Fig. 9; Supplementary Table 11).

The effect of reversing the concurrent model's response was assessed by treating the contemporaneous outcome as the response and standardized bilateral CR as the predictor. This response direction matched the prospective models (see Methods, Concurrent associations and Prospective associations sections). The following model was fitted:

$$X_{ij} = \beta_0 + \beta_R R^q_{ij} + \beta^T_D D_{ij} + \gamma^T C_{ij} + V_{ij} + \varepsilon_{ij}.$$

$X_{ij}$ denoted the outcome and $R^q_{ij}$ standardized bilateral LC or SN-VTA CR. The diagnosis groups and fixed covariates were represented by $D_{ij}$ and $C_{ij}$, as in the primary concurrent model, except that hemisphere was omitted. The CR-by-diagnosis interaction $\beta^T_{RD}\left(R^q_{ij} D_{ij}\right)$ was additionally included for cognitive outcomes. $V_{ij}$ contained participant and site intercepts and the participant imaging-time slope, as in the prospective model. A scan-set intercept was not included (Supplementary Fig. 8, first pair).

To assess associations conditional on the other CR measure, the single-CR term in the outcome-response concurrent and primary prospective models was replaced by:

$$\beta_{LC} R^{LC}_{ij} + \beta_{SN-VTA} R^{SN-VTA}_{ij}.$$

Both CR measures were entered as fixed effects. All remaining model terms were unchanged. Separate LC and SN-VTA models were also fitted, with the other CR term omitted. LC unique and SN-VTA unique referred to the conditional coefficients (Supplementary Fig. 8, pairs 2-3 and 5-6; Supplementary Tables 9, 11).

The estimated agreement between models was summarized in Supplementary Table 10.

#### Analyses restricted to CN and MCI at scan

Dependence on the diagnostic grouping was assessed by restricting the prospective and lag analyses to participants classified as CN or MCI at the LC/SN-VTA MRI, including those who subsequently progressed. CN versus MCI at that MRI was included as a covariate. The primary 180-1,095-day prospective window was retained. For cognitive outcomes, a common CR slope adjusted for CN/MCI was estimated instead of separate diagnosis-group slopes (see Methods, Prospective associations section):

$$Y_{ij}^{F} = \alpha_0 + \alpha_B Y_{ij}^{B} + \alpha_R R_{ij}^{q} + \alpha_D D_{ij}^{CN/MCI} + \alpha_L L_{ij} + \delta^T C_{ij} + V_{ij} + \eta_{ij}.$$

$Y^B$ and $Y^F$ denoted baseline and future outcome values; $R^q$ was standardized bilateral CR; $L$ was the MRI-to-assessment interval; and $C$ and $V$ were the primary fixed covariates and random effects. $D^{CN/MCI}$ distinguished CN from MCI at MRI, and $\eta$ was residual error. The same diagnosis restriction and covariate replacement were applied to the lag models (Supplementary Fig. 8, fourth pair; Supplementary Figs. 16-19).

#### Lag-dependent sensitivity analyses

The primary GAM specification and simultaneous confidence-band procedure were retained (see Methods, Lag-dependent generalized additive model section). Sensitivity models were fitted using assessments at least 180 days after MRI, the CN/MCI restriction above, or models with both CR measurements. In the model containing both measurements, the CR main-effect and lag-smooth terms were replaced by:

$$\theta_{LC} R^{LC} + R^{LC} s_{LC}(l) + \theta_{SN-VTA} R^{SN-VTA} + R^{SN-VTA} s_{SN-VTA}(l).$$

$l$ denoted elapsed time after the LC/SN-VTA MRI, and $s_{LC}$ and $s_{SN-VTA}$ were the separate lag-dependent smooths. (Supplementary Figs. 12-23). The number and direction of intervals excluding zero were summarized from Supplementary Table 12 (Supplementary Table 13).

The primary and sensitivity curves in Fig. 3c and Supplementary Figs. 10-23 were evaluated over the minimum-to-maximum lag separately by outcome and nucleus. Simultaneous confidence bands were calibrated over the corresponding evaluation grid. Model curves over the 5th-95th-percentile lag ranges, with their range-specific simultaneous bands, are provided for comparison in Supplementary Figs. 24-27. For graphical highlighting, each contiguous zero-excluding interval had to last at least 20% of its corresponding curve span (maximum minus minimum lag). Curves without a zero-excluding interval use reduced line opacity (0.35).

**Unique structural patterns and molecular correspondence**

The unique PLSC analyses were performed using the residualization and otherwise identical procedures specified in the main manuscript (see Methods, Partial least squares correlation analysis of whole-brain structural images section). Molecular correspondence was evaluated for the two unique intercept patterns (see Methods, Spatial correspondence with neurotransmitter receptor and transporter maps section; Supplementary Fig. 29; Supplementary Tables 15-17). Complete salience and BSR values by region are supplied in Supplementary Table 14; direct salience/BSR displays are in Supplementary Fig. 28.

**Post hoc PET-map adjustment**

To examine whether the exploratory molecular correspondences persisted after accounting for spatial overlap with the primary catecholaminergic PET maps, we conducted additional partial correlation analyses. For LC, we adjusted for the Hesse MRB norepinephrine-transporter map; for SN-VTA, we adjusted for the SCH23390, FLB457, and fallypride dopamine-receptor maps, both separately and jointly. Partial Spearman correlations were calculated between the ranked cortical saliences and exploratory PET distributions after removing their associations with the ranked control maps. To account for spatial autocorrelation, we evaluated these correlations against 10,000 Moran-spectral surrogate maps generated from the residualized saliences, with randomization constrained to remain orthogonal to the control maps. P values were calculated from the proportion of surrogate correlations at least as extreme as the observed correlation, with a +1 correction, followed by FDR correction across the 16 exploratory maps within each nucleus and adjustment model.

**Clinical progression with both LC and SN-VTA CR (mutually adjusted models)**

Both adjusted bilateral CR measurements were included to assess their conditional progression associations (see Methods, Five-year clinical progression section):

$$logitPr\left(Y_{is} = 1\right) = \alpha + \beta_{LC} Z_i^{LC} + \beta_{SN-VTA} Z_i^{SN-VTA} + \gamma^T C_i + u_s.$$

$Y$ denoted progression, $Z$ the standardized CR measurements, $C$ the primary covariate vector, and $u_s$ the site intercept. (Supplementary Fig. 31; Supplementary Tables 18, 19).

Odds ratios are reported per one SD lower CR, holding the other nucleus constant in the joint model. Their reported P values are unadjusted; the primary separate-model FDR family comprises LC and SN-VTA within each endpoint (Supplementary Table 18).

**Standardization for progression and mediation analyses**

For the logistic progression analyses, the SDs of bilateral technically adjusted CR were 0.04374 for LC and 0.02625 for SN-VTA, calculated from 3,412 and 3,404 eligible MRI examinations, respectively.

These calculations included repeated examinations at both field strengths and preceded selection of the progression samples. The same means and SDs were used for both progression outcomes.

The primary mediation analyses standardized CR, brain scores, and TPV separately for each nucleus and outcome. For MCI-to-AD dementia, means and SDs were calculated in the 233 eligible participants (one non-AD dementia case was excluded afterward); the fitted models included 232 participants. For any progression, they were calculated in the 335 included participants. Models with and without TPV adjustment used the same standardized values.

**Mediation using CN/MCI-trained PLSC patterns**

To assess dependence on the training population, LC and SN-VTA intercept PLSCs were re-estimated separately in participants classified as CN or MCI. The original regional-volume and CR intercepts were retained, but all 145 regional weights were re-estimated within the restricted population using the primary PLSC procedure, with 5,000 permutations and 5,000 bootstraps (see Methods, Partial least squares correlation analysis of whole-brain structural images section). The resulting participant-level PLSC intercept LV1 values replaced the original values in the same primary mediation samples, with and without TPV adjustment (Supplementary Table 21). CR and TPV used the same means and SDs as in the corresponding primary mediation analysis. The new brain scores were standardized in the same 233-participant or 335-participant samples used to standardize the primary mediation variables. (One non-AD dementia case was excluded afterward)

**Temporal-order mediation sensitivity: T1 after LC/SN-VTA MRI**

Two further sensitivity analyses required a T1 acquisition after the LC/SN-VTA MRI; events occurring before or at the selected T1 scan were excluded, while eligible participants without progression were retained. Each model retained the original CR, diagnosis, covariates, and progression outcome from the corresponding primary mediation model (see Methods, Associational indirect effects through distributed structural brain patterns section).

The first T1 acquisition within 365 days after the LC/SN-VTA MRI was selected. Participants whose first progression event preceded or coincided with that T1 scan were excluded. In the first analysis, the original participant-level PLSC intercept LV1 values from repeated T1 scans were used to establish the mediation model (Supplementary Table 22, Original PLSC).

In the second analysis, the LC and SN-VTA PLSCs were re-estimated in the pooled CN/MCI population meeting the later-T1 structural-availability requirements (Supplementary Table 22, Re-estimated PLSC).

Both temporal analyses retained the primary TPV estimate and the primary SEM specification, effect definitions, and indirect-effect correction procedure. Indirect effects are summarized in Supplementary Table 19. Brain scores were standardized within the corresponding later-T1 sample, separately for the original and re-estimated PLSC patterns. CR and TPV instead used the means and SDs from the corresponding primary mediation analysis.

**Amyloid-memory sensitivity analysis and additional displays**

To assess whether the findings depended on classifying amyloid as positive or negative, the analyses were repeated using continuous Centiloid burden instead of binary amyloid status. All other model terms and covariates were retained (see Methods, Amyloid-related longitudinal memory resilience section). Predicted memory trajectories and annual memory change across LC or SN-VTA CR values were presented in Supplementary Fig. 34. Interaction estimates were reported in Supplementary Tables 19, 23.

### Supplementary Note

**Cohort characteristics and MRI availability**

A total of 3,782 matched MRI scan sets from 739 participants were included in this study. Each scan set comprised one dual-echo PD-T2 acquisition sensitive to catecholaminergic nuclei and the same-field-strength T1-weighted scan used to derive atrophy estimates. The sample comprised a 1.5-T dataset including 3,273 scan sets from 710 participants, as well as a 3-T dataset including 509 scan sets from 145 participants; 116 participants contributed at both field strengths. Following manual quality control, 3,457 matched scan sets from 731 participants were retained (exclusion rates were 8.7% at 1.5 T and 7.9% at 3 T). After extreme-value screening, 3,428 scan sets from 726 participants contributed at least one LC or SN-VTA CR measure from either hemisphere, and 3,390 scan sets from 724 participants contributed CR measurements in both hemispheres of both nuclei.

Across the 3,763 scan sets with age information, age at imaging was 77.1 ± 7.0 (Mean ± SD) years. 321 of 739 participants were female (43.4%), and 367 of 739 carried at least one apolipoprotein E (APOE) ε4 allele (49.7%). Baseline diagnosis was available for 720 of 739 participants (97.4%), including 204 who were CN, 350 with MCI, and 166 with AD. Diagnosis at imaging was available for 3,694 scan sets, including 1,136 CN, 1,507 MCI, 1,048 AD, and 3 non-AD dementia. Participant demographic characteristics are reported in Supplementary Table 1.

Participants contributed a mean of 5.12 ± 2.90 matched MRI scan sets. Among the 663 participants with repeated imaging, the mean interval between consecutive scan sets was 0.72 ± 0.29 years, and the mean first-to-last MRI interval was 3.39 ± 2.42 years (range, 0.04-10.29 years). Field strength-specific summaries are provided in Supplementary Table 1.

**Hemispheric differences by scanner manufacturer and field strength**

SN-VTA CR was higher on the left in every manufacturer and field-strength group (all $P \leq 0.004$; Supplementary Table 8). For LC, GE scans showed no clear hemispheric difference, whereas Siemens scans showed higher CR on the right at 1.5 T and on the left at 3 T. Philips estimates indicated higher right LC CR and higher left SN-VTA CR, but this group included only 11 participants and 20 scans, all at 3 T.

**Alternative concurrent models and mutual adjustment**

We fitted alternative concurrent models with each outcome as the dependent variable and bilateral LC or SN-VTA CR as the predictor. Coefficient directions agreed between these specifications for 31 of 38 LC and 29 of 38 SN-VTA estimates. We then compared separate outcome-response models with models containing both CR measurements. Including both measurements retained 36 of 38 LC and 37 of 38 SN-VTA coefficient directions; correlations between the separate-model and mutually adjusted coefficients were $r = 0.992$ and $r = 0.996$, respectively. All three associations that were FDR significant in the separate models remained significant after mutual adjustment. Higher LC CR was associated with fewer difficulties keeping track of current events, whereas higher SN-VTA CR was associated with less impairment in home and hobbies and fewer difficulties with appointments or medications, after accounting for the other CR measurement (Supplementary Fig. 8; Supplementary Tables 9 and 10).

**Prospective assessment windows and mutual adjustment**

We evaluated alternative prospective assessment windows requiring assessments at least 180 or at least 90 days after MRI, without an upper time limit. Across these alternatives, 36-37 of 38 LC and all 38 SN-VTA coefficients retained their direction, with correlations across coefficients of $r = 0.994$-$0.997$. Including both CR measurements retained 36 of 38 coefficient directions for each nucleus relative to separate models fitted to the same observations (LC, $r = 0.991$; SN-VTA, $r = 0.989$). Seven of nine LC and two of nine SN-VTA associations that were FDR significant in these separate models remained significant after mutual adjustment. The retained LC associations involved memory, language and verbal fluency; the retained SN-VTA associations involved community functioning and travel or transportation, assessed using CDR Community Affairs and FAQ Travel or Transportation (Supplementary Figs. 8 and 9; Supplementary Tables 10 and 11).

In the alternative diagnosis specification restricted to CN or MCI at MRI, coefficient directions agreed with the primary prospective models for 31 of 37 LC and 33 of 37 SN-VTA estimates ($r = 0.840$ and $r = 0.813$, respectively). (Supplementary Fig. 8; Supplementary Table 10).

**Lag-dependent models**

For lag models presented in Supplementary Figs. 10 to 11, higher LC CR was associated with larger left and right hippocampal volumes at 7.49-18.84 and 9.18-18.84 years after MRI, respectively, and larger left and right entorhinal volumes at 7.08-18.84 and 8.75-17.69 years. Higher LC CR was also associated with lower SPARE-AD scores at 8.27-18.84 years. Higher SN-VTA CR was associated with larger left hippocampal volume at 6.84-18.84 years. Both measures were associated with lower WMH burden at later follow-up: 5.65-19.98 years for LC and 6.46-16.14 years for SN-VTA. These intervals were defined by within-curve simultaneous 95% confidence bands excluding zero (Fig. 3c; Supplementary Fig. 11; Supplementary Table 12).

We assessed sensitivity to assessment selection, diagnostic specification and inclusion of both CR measurements. Across the evaluated specifications, 25-26 LC and 21-25 SN-VTA curves contained at least one interval in which the within-curve simultaneous 95% confidence band excluded zero. Positive memory-composite intervals at later follow-up were retained across all specifications. After mutual adjustment, 25 LC and 24 SN-VTA curves contained an interval excluding zero. These included positive associations with the memory composite and RAVLT immediate learning, delayed recall and recognition for both CR measurements (Supplementary Figs. 12-23; Supplementary Tables 12 and 13).

The late LC associations with bilateral hippocampal and entorhinal volumes and SPARE-AD, the SN-VTA association with left hippocampal volume, and the WMH associations for both nuclei retained intervals excluding zero after accounting for the other CR measurement. The mutually adjusted SN-VTA model additionally showed a negative association with left entorhinal volume at 8.52-11.88 years (Supplementary Figs. 22 and 23; Supplementary Table 12).

**Structural covariance after accounting for the other nucleus**

After residualizing each CR parameter against the corresponding parameter of the other nucleus, the LC PLSC associations remained significant for intercepts ($r = 0.219$; $P_{perm} < 0.001$) and rates of change ($r = 0.173$; $P_{perm} = 0.019$). The corresponding SN-VTA associations were $r = 0.250$ ($P_{perm} < 0.001$) and $r = 0.173$ ($P_{perm} = 0.013$), respectively (Supplementary Fig. 29; Supplementary Tables 15 and 16).

**Molecular-map correspondence after mutual adjustment**

After accounting for LC CR, all three SN-VTA dopamine-receptor correspondences remained FDR significant ($\rho = 0.472$-$0.584$; $P_{FDR} = 0.002$-$0.006$). After accounting for SN-VTA CR, the LC pattern retained a positive correspondence with the Hesse et al. norepinephrine-transporter map, but this correspondence did not meet the FDR threshold ($\rho = 0.361$; nominal $P = 0.008$; Supplementary Fig. 29; Supplementary Tables 16 and 17).

**Molecular-map correspondence after NET or DA maps adjustment**

Among the 16 exploratory PET maps, eight showed significant spatial correspondence with the SN-VTA intercept pattern before PET-map adjustment ($P_{FDR} < 0.05$). After adjusting separately for the SCH23390, FLB457, and fallypride dopamine-receptor maps, three, one, and four correspondences, respectively, met the exploratory adjusted threshold ($P_{Partial,FDR} < 0.05$). None met this threshold when all three dopamine-receptor maps were included together. For LC, none of the 16 exploratory correspondences met the corresponding threshold before or after adjustment for the Hesse MRB NET map (Supplementary Table 17). The attenuation of the SN-VTA associations suggests that these exploratory correspondences may partly reflect spatial variation shared with dopamine-receptor distributions.

**Clinical progression after mutual adjustment**

When both CR measurements were included, one SD lower LC CR was associated with higher odds of MCI-to-AD progression (OR = 1.828; 95% CI, 1.217-2.747; P = 0.004) and any diagnostic progression (OR = 1.457; 95% CI, 1.068-1.987; P = 0.017). The corresponding SN-VTA associations were not significant (P = 0.373 and P = 0.059, respectively). (Supplementary Fig. 31; Supplementary Tables 18 and 19).

**Indirect associations: Models without TPV adjustment**

Without TPV adjustment, the LC and SN-VTA indirect effects were −0.194 and −0.158, respectively, for MCI-to-AD progression, compared with −0.148 and −0.085 in the TPV-adjusted models. For any progression, the corresponding estimates were −0.135 and −0.145 without TPV adjustment, compared with −0.091 and −0.062 with adjustment. All four indirect associations remained significant after FDR correction in the models without TPV adjustment (Fig. 5b, right column; Supplementary Figs. 32 and 33; Supplementary Table 20).

**Indirect associations: PLSC training population and later-T1 eligibility**

Re-estimating the PLSC patterns in a CN/MCI training population retained all eight FDR-significant indirect associations across the two nuclei, two progression outcomes and models with or without TPV adjustment (Supplementary Table 21). Further analyses required a T1 scan within 365 days after the LC/SN-VTA MRI and excluded progression events occurring at or before that scan. Using the original PLSC patterns, the TPV-adjusted LC indirect effects were −0.184 for MCI-to-AD dementia progression ($P_{FDR}$ = 0.038) and −0.106 for any progression ($P_{FDR}$ = 0.049). Re-estimating the PLSC patterns in the eligible CN/MCI population yielded corresponding effects of −0.213 and −0.137 (both $P_{FDR}$ = 0.025). All four corresponding SN-VTA indirect-effect confidence intervals included zero. (Supplementary Tables 19 and 22).

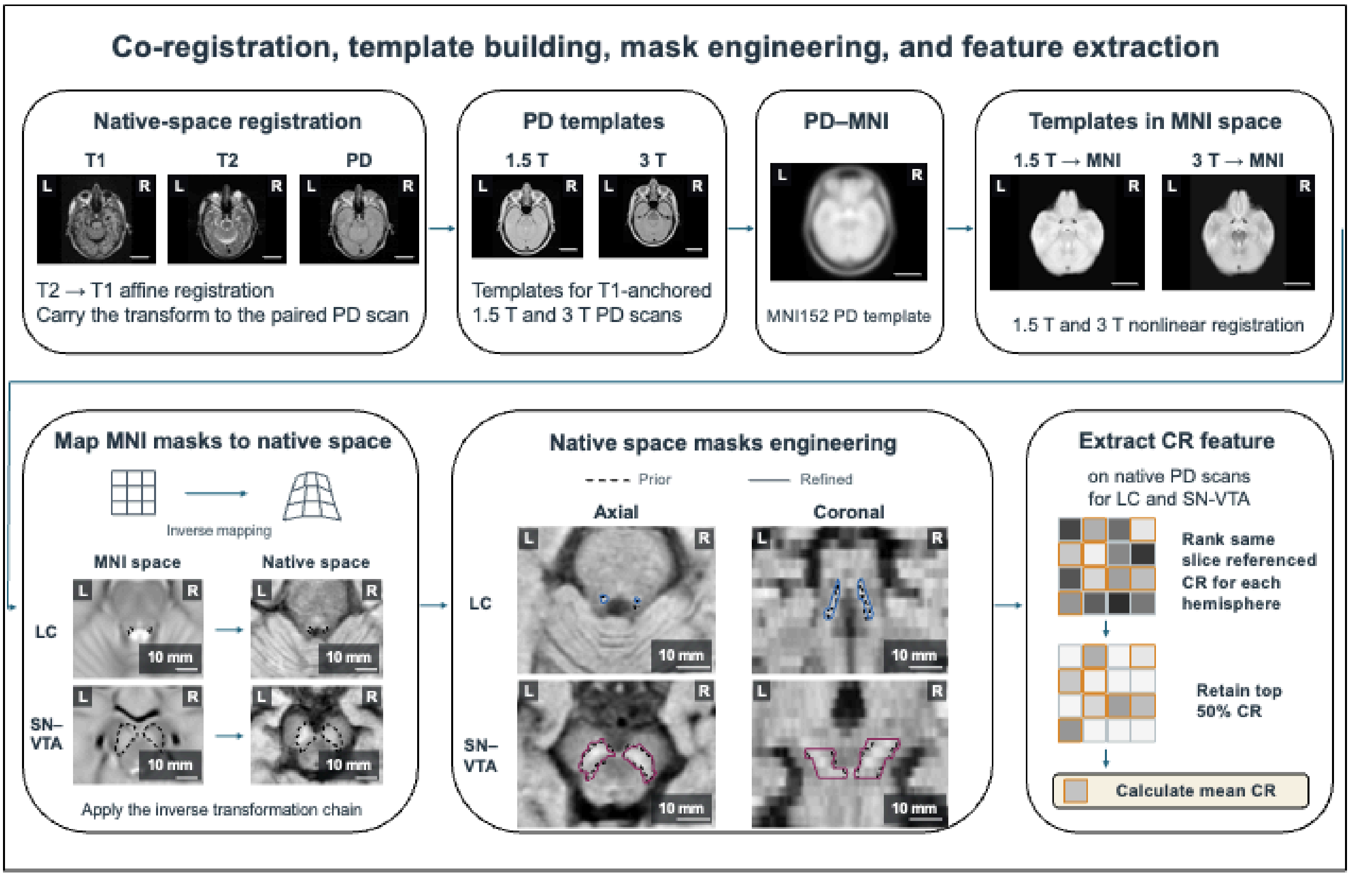


**Supplementary Figure 1**. **Scan-processing flowchart.** T2-weighted images were affinely registered to T1-weighted images, and the resulting transformations were applied to the paired PD scans. Separate 1.5-T and 3-T PD templates were constructed and nonlinearly registered to the MNI152 PD template. Meta masks in the MNI space of the LC and SN-VTA were then mapped from MNI space back to native PD space using the inverse transformations and refined in axial and coronal views. Dashed outlines indicate the initial masks, and solid outlines indicate the refined masks. For each hemisphere, slice-referenced CRs were calculated within the masks and ranked. The highest 50% of CR values were retained, and their mean was used as the hemispheric CR. L and R indicate left and right, respectively.

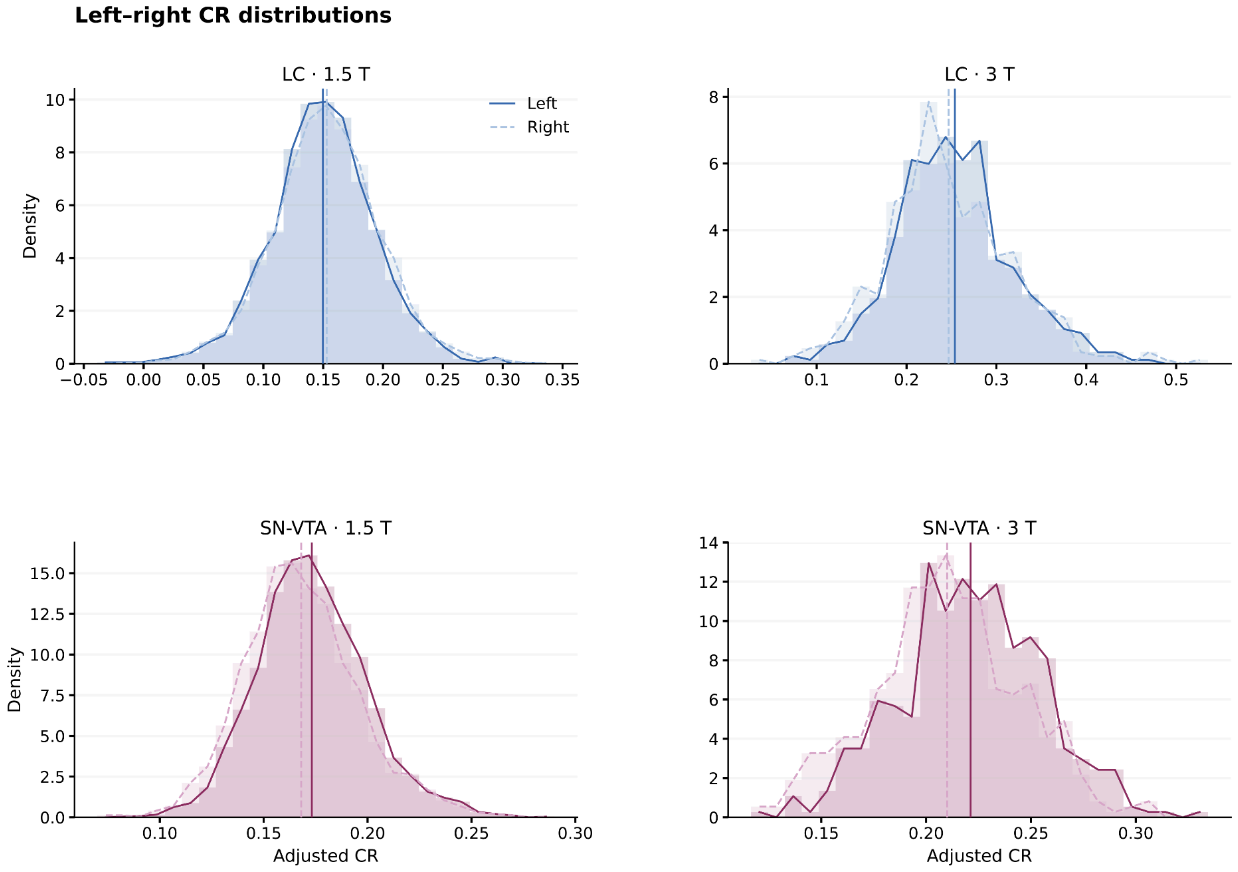


**Supplementary Figure 2. Left-right LC and SN-VTA CR distributions by field strength.** Hemisphere-specific LC and SN-VTA CR distributions are shown separately at 1.5 T and 3 T. Dark solid curves indicate left-hemisphere observations and lighter dashed curves indicate right-hemisphere observations.

CR, contrast ratio; LC, locus coeruleus; SN-VTA, substantia nigra-ventral tegmental area.

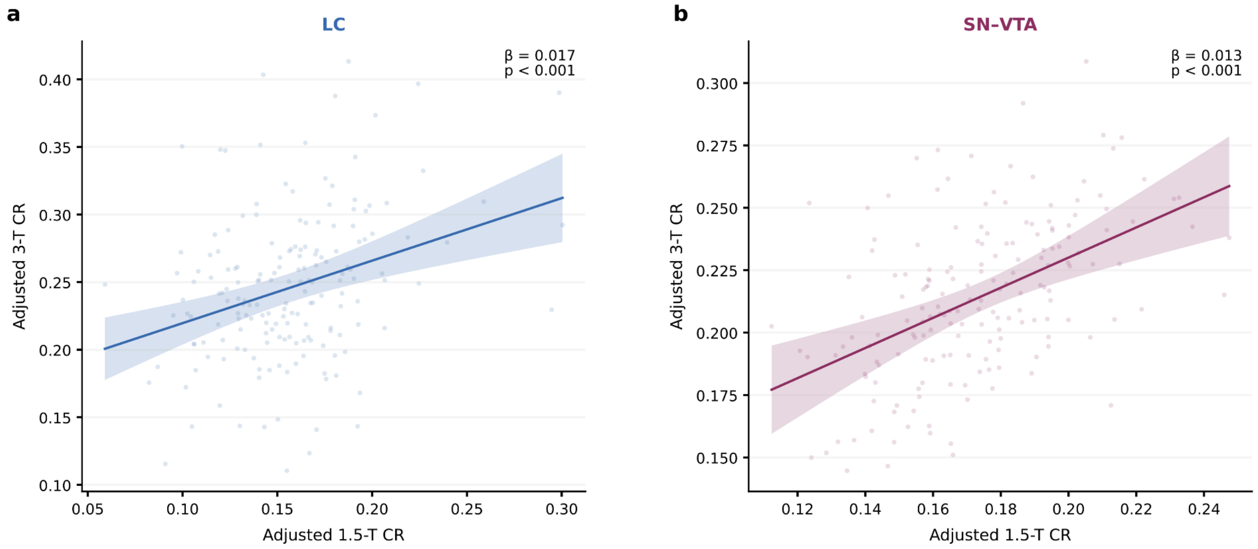


**Supplementary Figure 3. LC and SN-VTA CR correspondence between 1.5 T and 3 T.** a, LC. b, SN-VTA**.** Points show adjusted bilateral LC (a) and SN-VTA (b) CR at the two field strengths across 176 paired scans from 86 participants. The solid line is a descriptive least-squares fit; shading indicates its pointwise 95% confidence interval. The β and P value are from the covariate-adjusted mixed model described in Supplementary Methods; β is expressed as the 3-T CR difference per SD of 1.5-T CR in the paired sample.

CR, contrast ratio; LC, locus coeruleus; SD, standard deviation; SN-VTA, substantia nigra-ventral tegmental area.

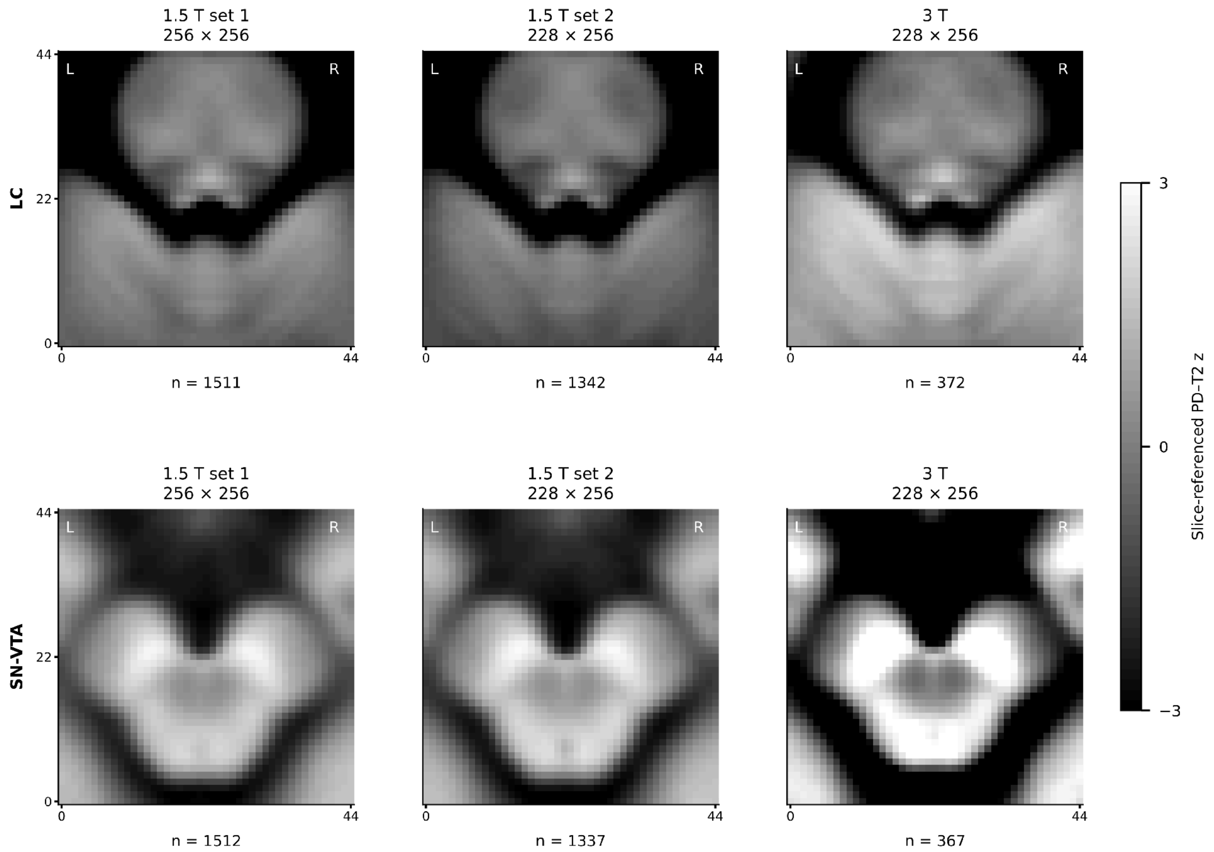


**Supplementary Figure 4**. Original acquisition-geometry-specific mean LC and SN-VTA images: 1.5-T set 1 (256 × 256 source matrix), 1.5-T set 2, and 3 T (228 × 256 source matrix). All panels retain 45 × 45-voxel local patches with 0.9375 × 0.9375-mm in-plane spacing and 3-mm slice spacing. The two 1.5-T acquisition-geometry-specific averages shown here were combined for the pooled 1.5-T display in main Fig. 1a.

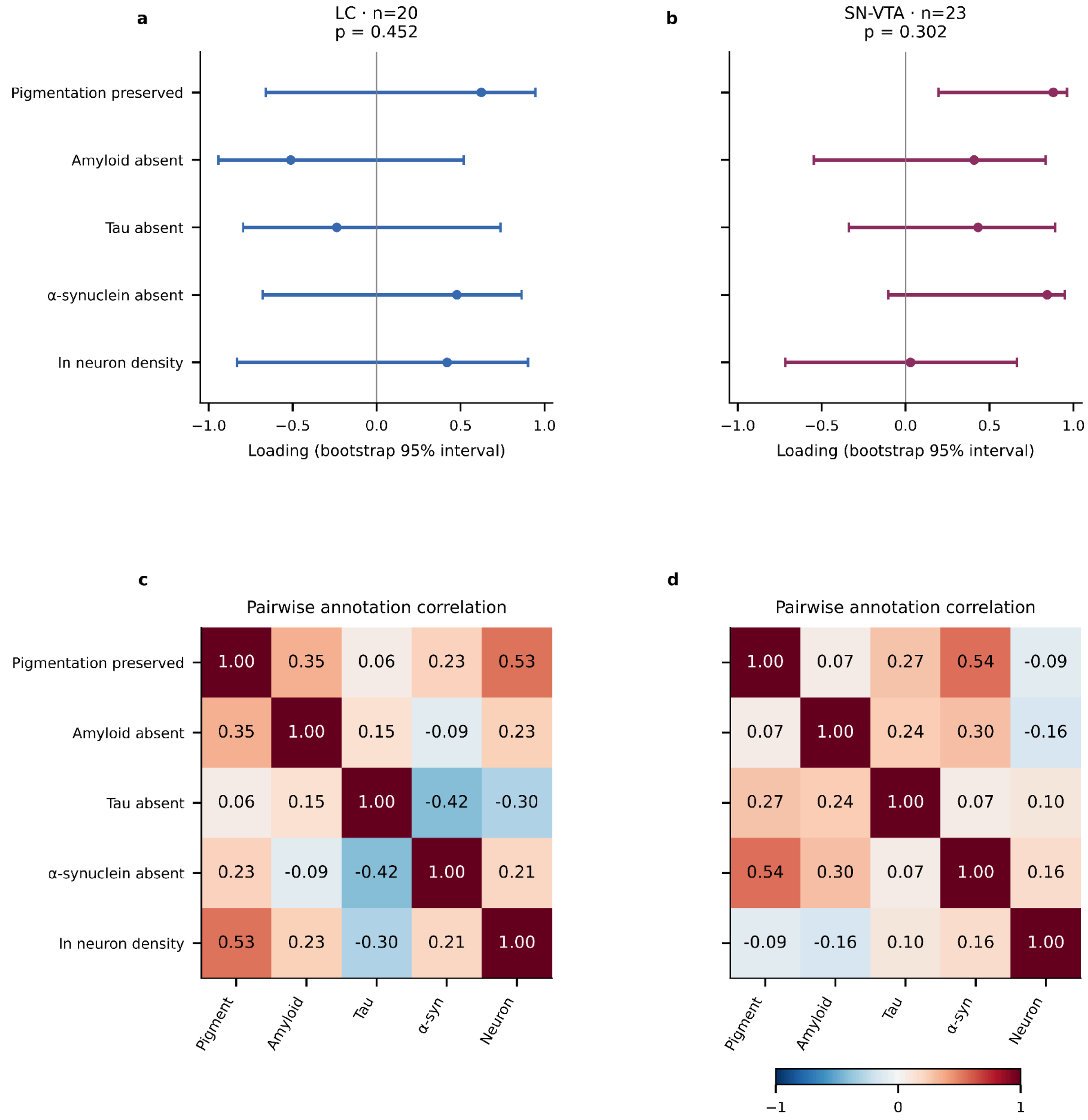


**Supplementary Figure 5. Neuropathological associations with LC and SN-VTA CR.** a,b, Loadings of the neuropathological measures for the LC and SN-VTA PLSCs, respectively. Points and horizontal bars show loadings and 95% bootstrap confidence intervals; the annotated P values refer to the global permutation tests. Complete-case samples contain 20 LC and 23 SN-VTA participants. c,d, Pairwise correlations among the neuropathological measures for LC and SN sections. Positive values indicate concordance between the two measures; the sample size varies by pair (LC, 22-30; SN, 25-32).

CR, contrast ratio; LC, locus coeruleus; PLSC, partial least squares correlation; SN, substantia nigra; SN-VTA, substantia nigra-ventral tegmental area.

**Supplementary Figure 6. In-vivo MRI and postmortem brainstem histology from the same participants.** Illustrative images from four participants are shown. Participants 1 and 2 show MRI and tissue sections of the LC. Participants 3 and 4 show MRI of the combined SN-VTA region and tissue sections of the SN. Within these pairs, Participants 1 and 3 had higher bilateral MRI CRs, after adjustment for confounding factors, and higher neuronal densities from histological annotations than Participants 2 and 4, respectively. Hypopigmentation was rated as absent (grade 0) in Participants 1 and 3 and mild (grade 1) in Participants 2 and 4.

In the overview, rows correspond to Participants 1-4, and columns show axial MRI, coronal MRI, hematoxylin and eosin (H&E) staining, and immunohistochemical staining for amyloid-β, tau and α-synuclein, from left to right. Red arrows on the in-vivo MRI mark the suggested LC or SN-VTA locations. Yellow outlines on tissue sections mark the annotated tissue regions. The overview is followed by participant-specific layouts and enlarged histological images, ordered by participant and then by stain: H&E, amyloid-β, tau and α-synuclein. MRI scale bars, 10 mm. Tissue scale bars are indicated in the panels; enlarged LC and SN images have scale bars of 200 μm and 500 μm, respectively. MRI, magnetic resonance imaging; L, left; R, right.

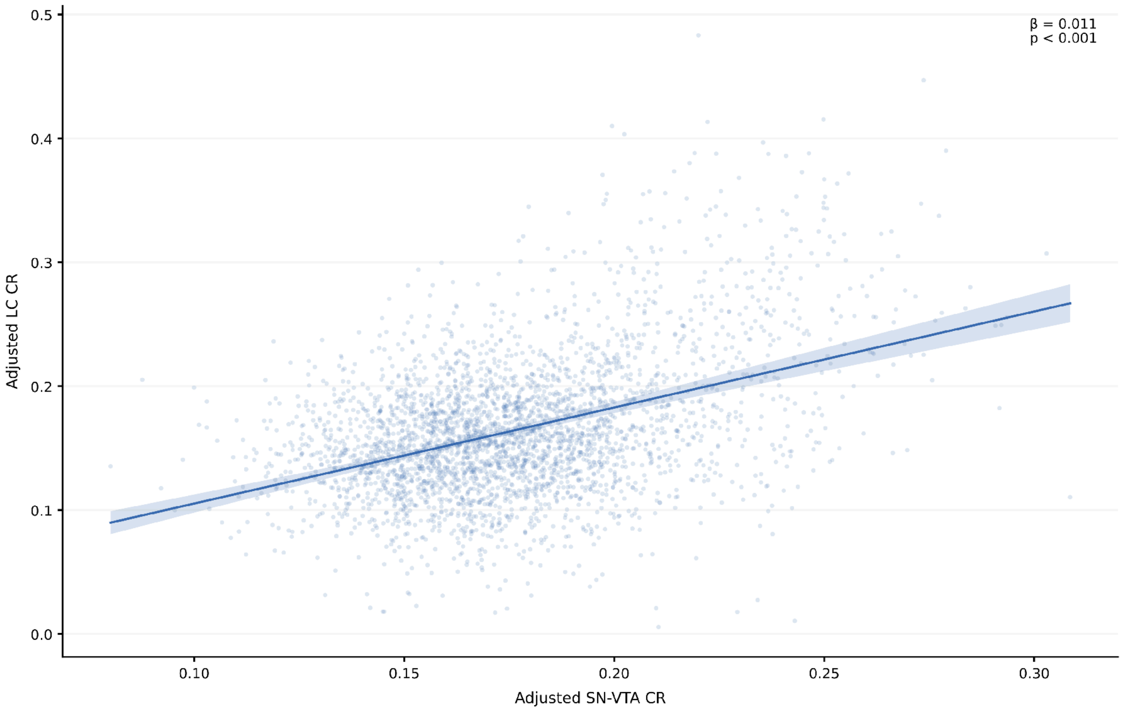


**Supplementary Figure 7. LC-SN-VTA CR association.** Points show adjusted SN-VTA CR and LC CR. The solid line is a descriptive least-squares fit; shading indicates its pointwise 95% confidence interval; β is expressed per SD of SN-VTA CR.

CR, contrast ratio; LC, locus coeruleus; MRI, magnetic resonance imaging; SD, standard deviation; SN-VTA, substantia nigra-ventral tegmental area.

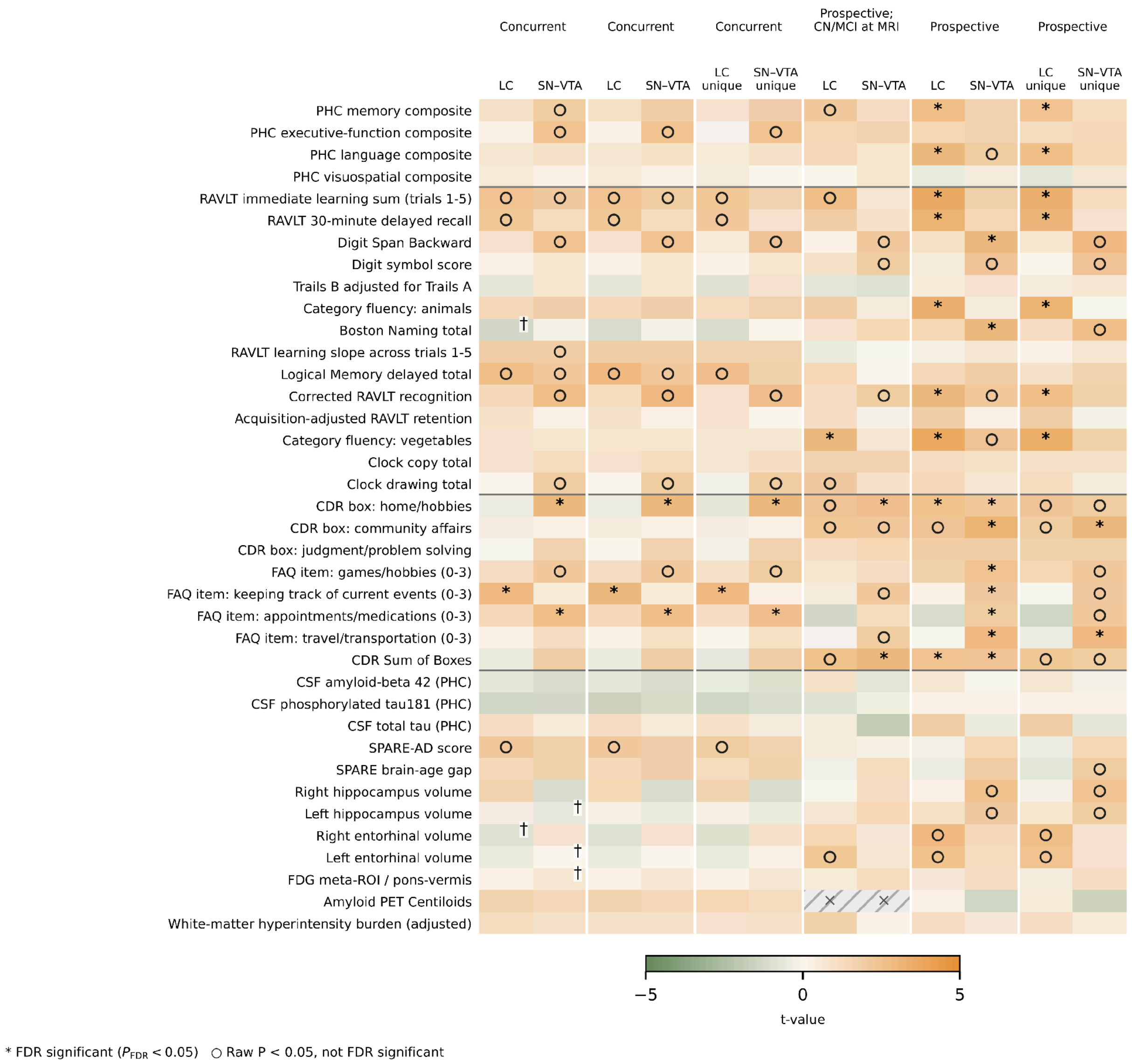


**Supplementary Figure 8. Concurrent and prospective LC and SN-VTA model comparisons.** The six LC/SN-VTA pairs, from left to right, are: Concurrent (bilateral CR predictor); Concurrent (separate models); Concurrent (LC and SN-VTA included); Prospective (CN/MCI at LC/SN-VTA MRI); Prospective (separate models); and Prospective (LC and SN-VTA included). "Unique" denotes adjustment for the other CR measure. Signs were reversed for color mapping for measures where lower values indicate better outcomes. Therefore, for all measures, orange indicates an association of higher CR with better outcomes and green the opposite association. Asterisks denote $P_{FDR} < 0.05$ and open circles denote $P < 0.05$ without meeting that FDR criterion. Daggers identify qualified fits; hatched/crossed cells identify unavailable results.

CN, cognitively normal; CR, contrast ratio; FDR, false discovery rate; LC, locus coeruleus; MCI, mild cognitive impairment; SN-VTA, substantia nigra-ventral tegmental area. Other outcome abbreviations follow main Fig. 3.

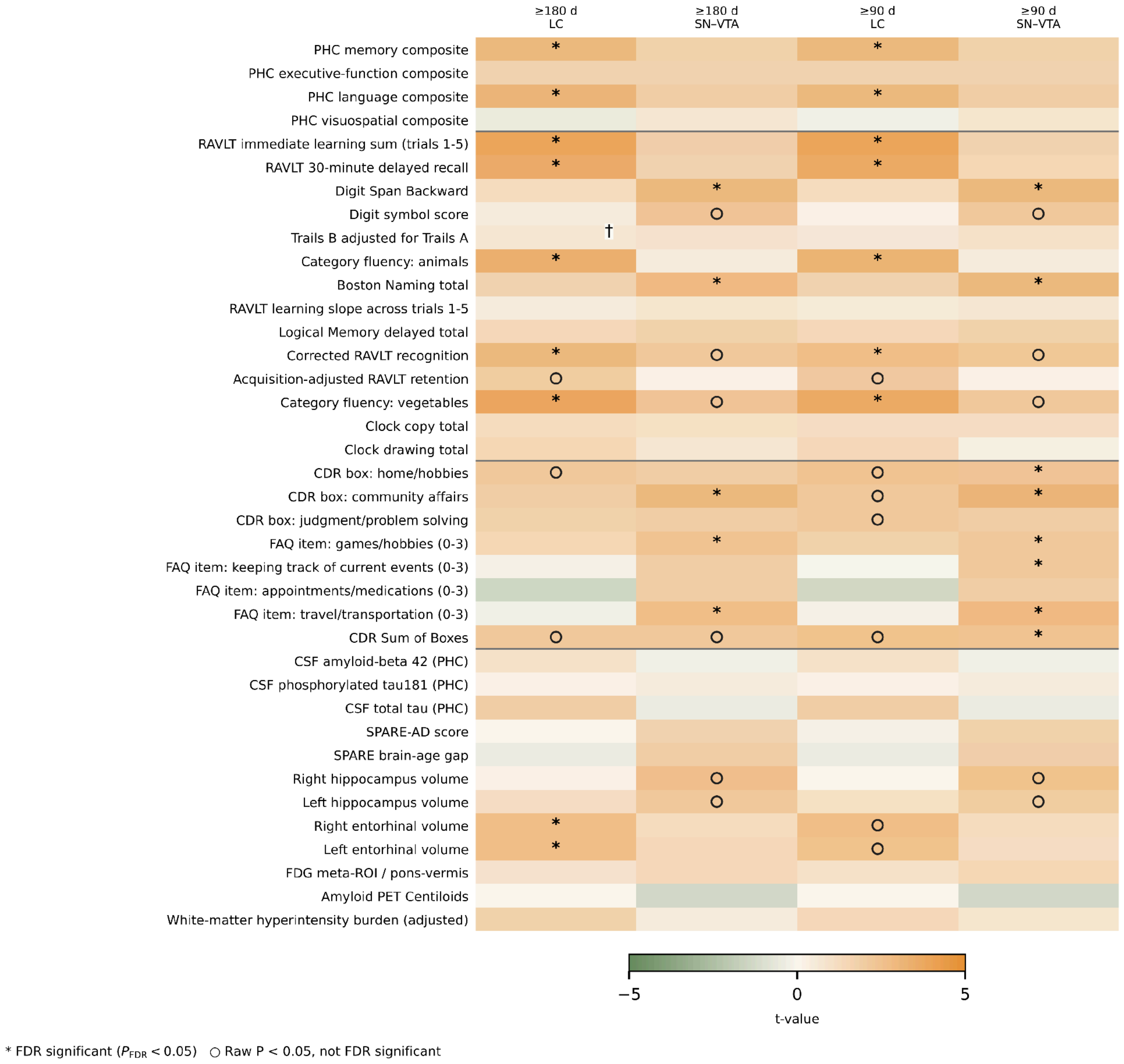


**Supplementary Figure 9. Prospective associations across alternative follow-up windows.** The left LC/SN-VTA pair shows "Prospective (≥180 days)," and the right pair shows "Prospective (≥90 days)." Both analyses remove the upper follow-up limit while retaining baseline-outcome adjustment and selection of the future assessment nearest day 365. Signs were reversed for color mapping for measures where lower values indicate better outcomes. Therefore, for all measures, orange indicates an association of higher CR with better outcomes and green the opposite association. Asterisks denote $P_{FDR} < 0.05$; open circles denote $P < 0.05$ without meeting that FDR criterion. Daggers identify qualified fits; hatched/crossed cells identify unavailable estimates.

CR, contrast ratio; FDR, false discovery rate; LC, locus coeruleus; SN-VTA, substantia nigra-ventral tegmental area. Outcome abbreviations follow main Fig. 3.

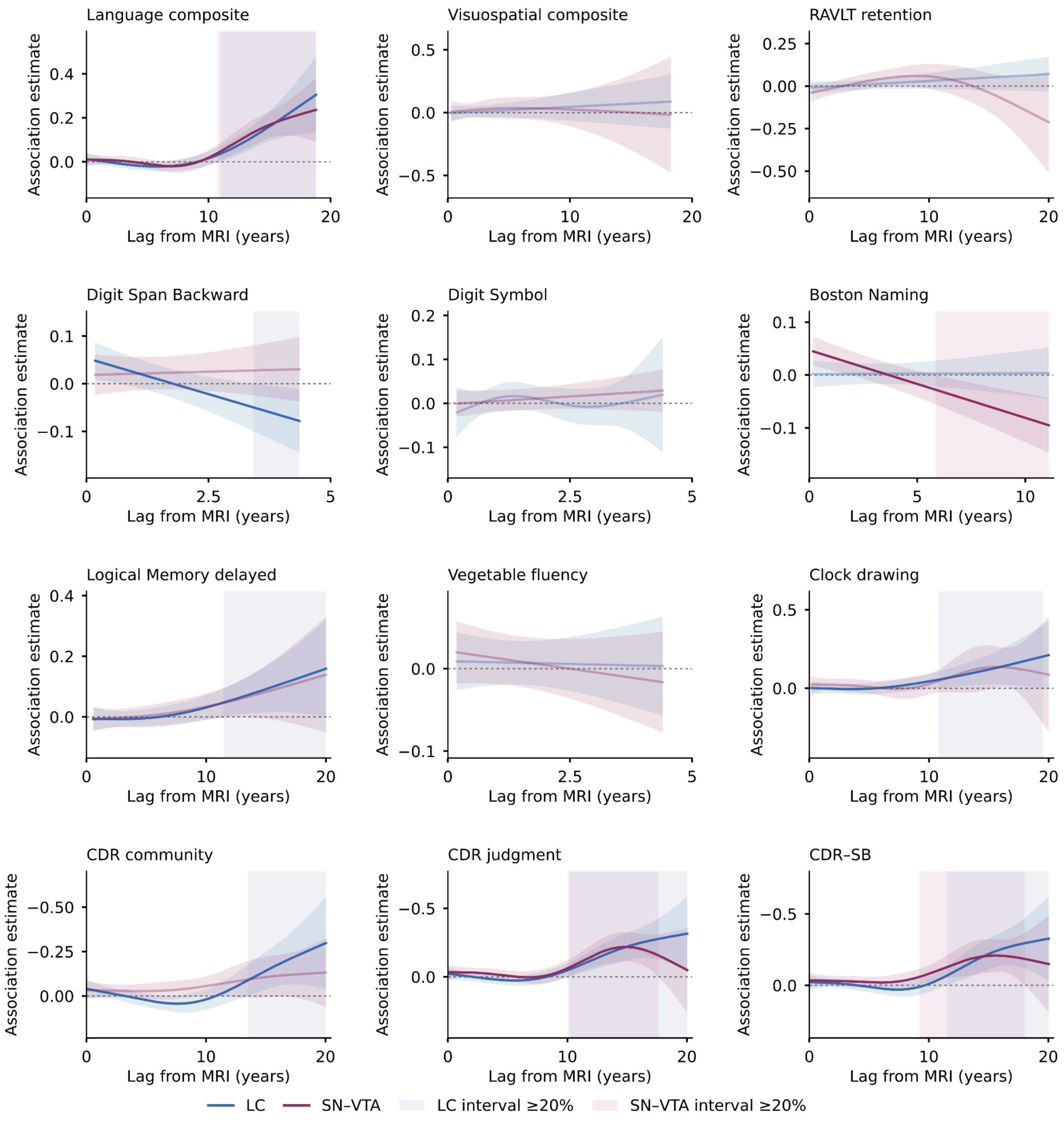


**Supplementary Figure 10. Lag-dependent associations with additional cognitive and clinical measures.** The remaining 20 outcomes from the primary all-assessment models are shown across Supplementary Figs. 10, 11; the other 18 outcomes appear in main Fig. 3c. Lines show the CR main effect plus its lag-dependent smooth per SD of CR as standardized for that model (LC, blue; SN-VTA, plum). Shaded ribbons are within-curve simultaneous 95% confidence bands; pale vertical shading denotes individual zero-excluding intervals lasting at least 20% of the corresponding curve's observed lag span. Shorter intervals remain reported in Supplementary Table 12. Curves without a zero-excluding interval use reduced line opacity (0.35). For measures where lower values indicate better outcomes, y-axes are reversed so that negative coefficients appear above zero; the original coefficient values and signs are retained. A higher association estimate is associated with better outcomes for all variables.

CR, contrast ratio; FDR, false discovery rate; LC, locus coeruleus; SD, standard deviation; SN-VTA, substantia nigra-ventral tegmental area. Outcome abbreviations follow main Fig. 3.

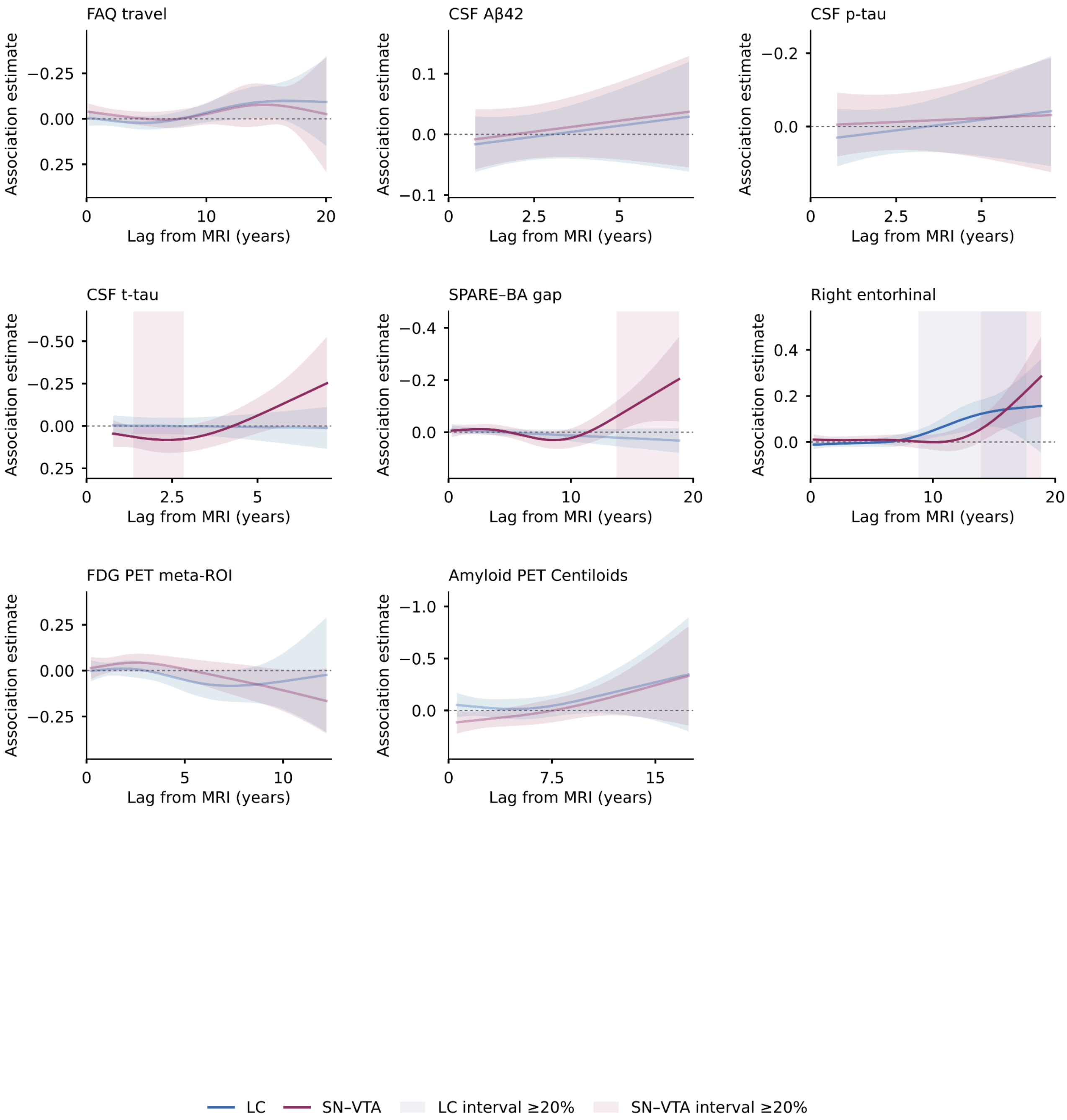


**Supplementary Figure 11. Lag-dependent associations with additional functional, molecular and brain-imaging measures.** The remaining 20 outcomes from the primary all-assessment models are shown across Supplementary Figs. 10, 11; the other 18 outcomes appear in main Fig. 3c. Lines show the CR main effect plus its lag-dependent smooth per SD of CR as standardized for that model (LC, blue; SN-VTA, plum). Shaded ribbons are within-curve simultaneous 95% confidence bands; pale vertical shading denotes individual zero-excluding intervals lasting at least 20% of the corresponding curve's observed lag span. Shorter intervals remain reported in Supplementary Table 12. Curves without a zero-excluding interval use reduced line opacity (0.35). For measures where lower values indicate better outcomes, y-axes are reversed so that negative coefficients appear above zero; the original coefficient values and signs are retained. A higher association estimate is associated with better outcomes for all variables.

CR, contrast ratio; FDR, false discovery rate; LC, locus coeruleus; SD, standard deviation; SN-VTA, substantia nigra-ventral tegmental area. Outcome abbreviations follow main Fig. 3.

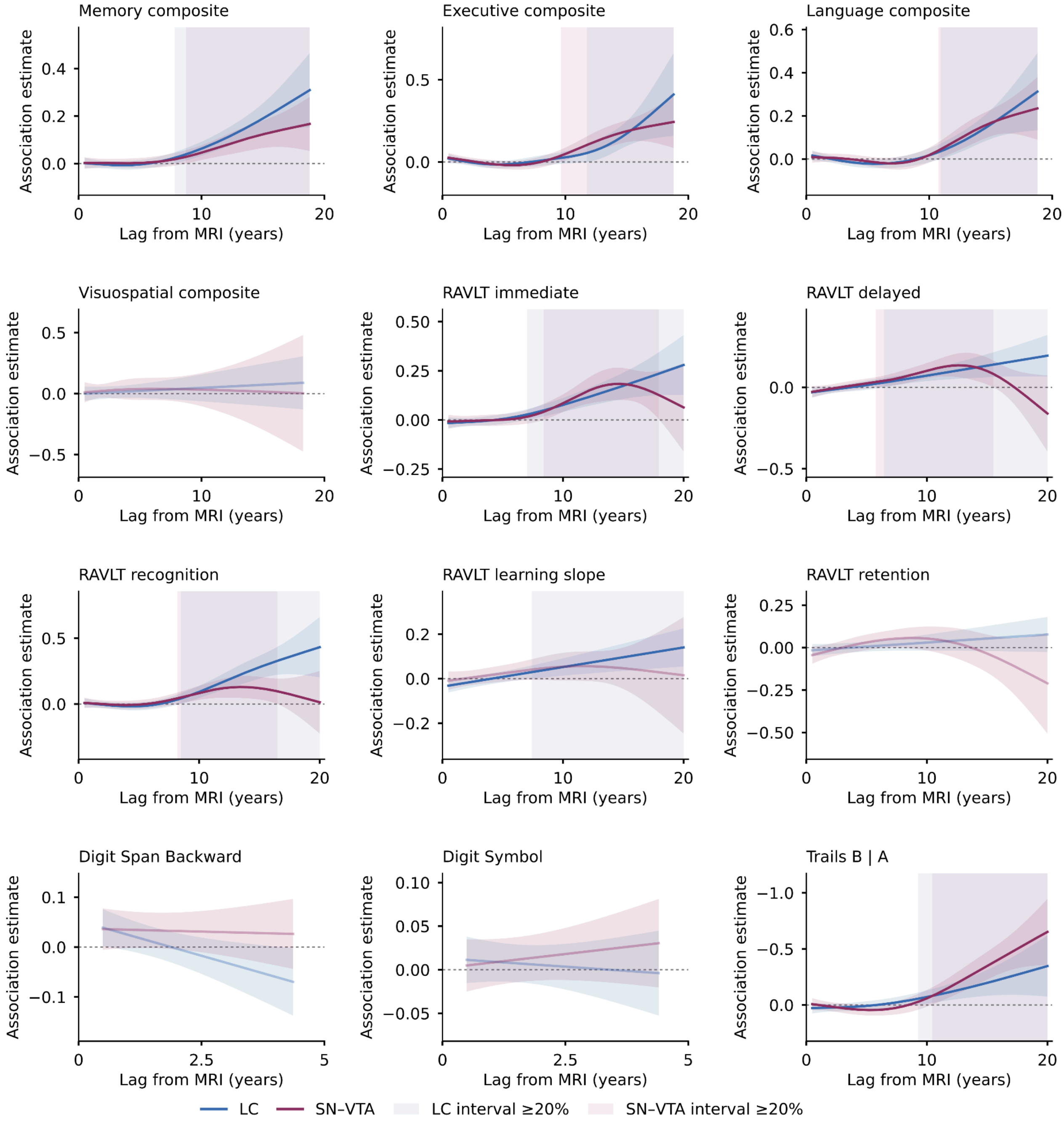


**Supplementary Figure 12. Lag-dependent associations from 180 days after MRI with cognitive composites and cognitive tests.** All 38 outcomes are shown across Supplementary Figs. 12-15. Models use assessments at least 180 days after the LC/SN-VTA MRI and are refitted, not cropped primary curves. Lines show the CR main effect plus its lag-dependent smooth per SD of CR as standardized for that model (LC, blue; SN-VTA, plum). Shaded ribbons are within-curve simultaneous 95% confidence bands; pale vertical shading denotes individual zero-excluding intervals lasting at least 20% of the corresponding curve's observed lag span. Shorter intervals remain reported in Supplementary Table 12. Curves without a zero-excluding interval use reduced line opacity (0.35). For measures where lower values indicate better outcomes, y-axes are reversed so that negative coefficients appear above zero; the original coefficient values and signs are retained. A higher association estimate is associated with better outcomes for all variables.

CR, contrast ratio; FDR, false discovery rate; LC, locus coeruleus; SD, standard deviation; SN-VTA, substantia nigra-ventral tegmental area. Outcome abbreviations follow main Fig. 3.

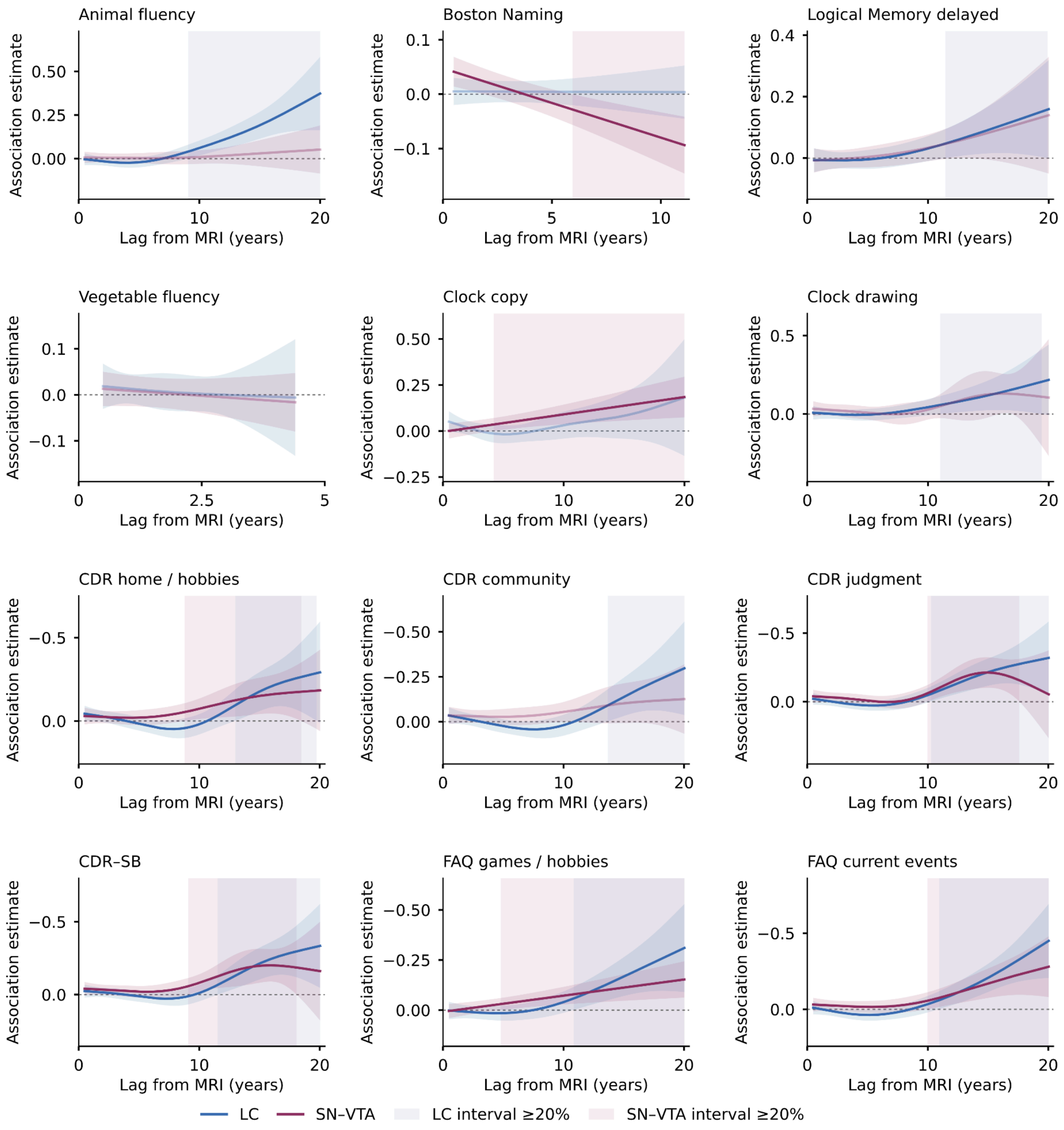


**Supplementary Figure 13. Lag-dependent associations from 180 days after MRI with additional cognitive, clinical and functional measures.** All 38 outcomes are shown across Supplementary Figs. 12-15. Models use assessments at least 180 days after the LC/SN-VTA MRI and are refitted, not cropped primary curves. Lines show the CR main effect plus its lag-dependent smooth per SD of CR as standardized for that model (LC, blue; SN-VTA, plum). Shaded ribbons are within-curve simultaneous 95% confidence bands; pale vertical shading denotes individual zero-excluding intervals lasting at least 20% of the corresponding curve's observed lag span. Shorter intervals remain reported in

Supplementary Table 12. Curves without a zero-excluding interval use reduced line opacity (0.35). For measures where lower values indicate better outcomes, y-axes are reversed so that negative coefficients appear above zero; the original coefficient values and signs are retained. A higher association estimate is associated with better outcomes for all variables.

CR, contrast ratio; FDR, false discovery rate; LC, locus coeruleus; SD, standard deviation; SN-VTA, substantia nigra-ventral tegmental area. Outcome abbreviations follow main Fig. 3.

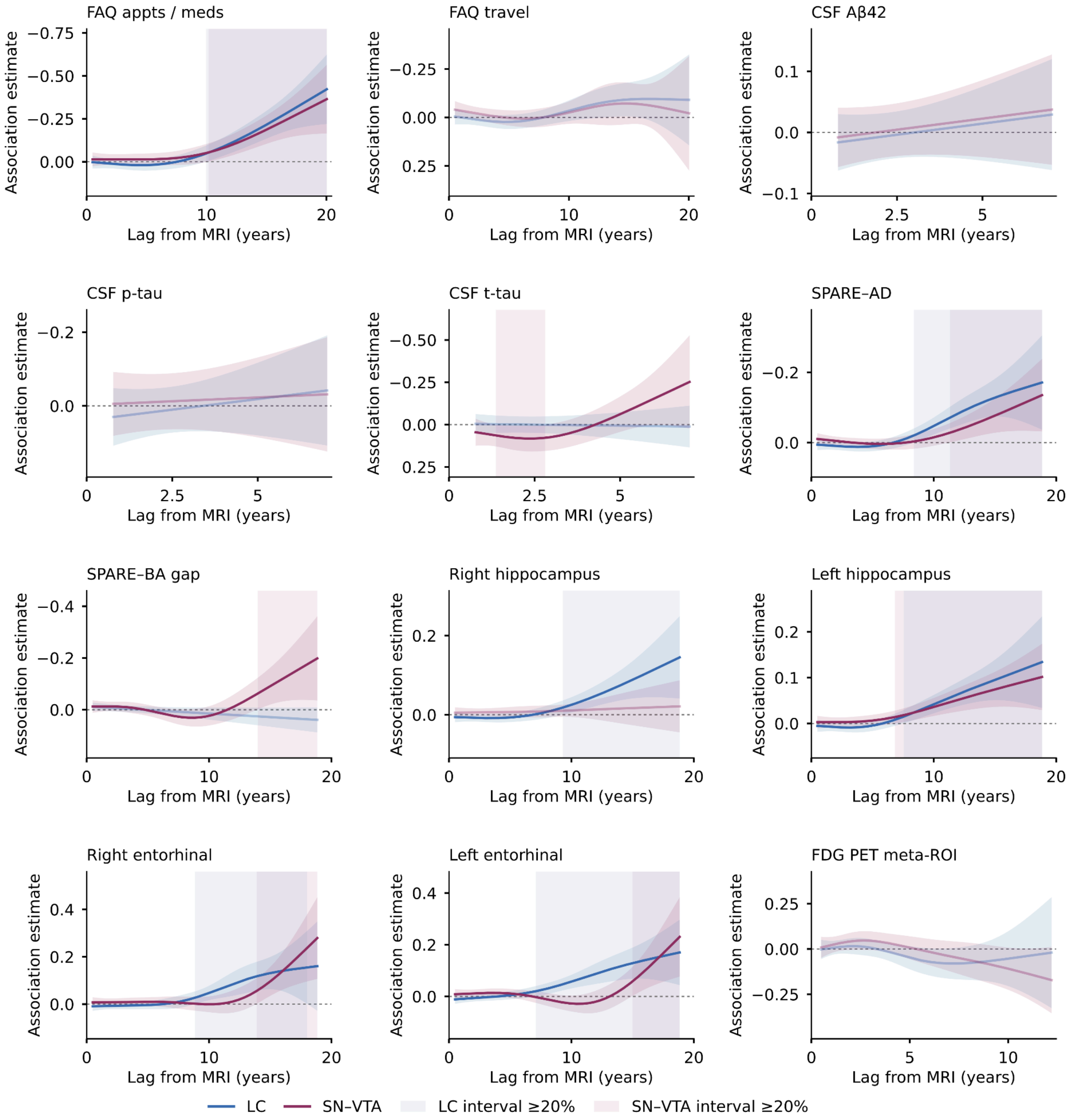


**Supplementary Figure 14. Lag-dependent associations from 180 days after MRI with clinical, functional and biological measures.** All 38 outcomes are shown across Supplementary Figs. 12-15. Models use assessments at least 180 days after the LC/SN-VTA MRI and are refitted, not cropped primary curves. Lines show the CR main effect plus its lag-dependent smooth **per SD of CR as**

standardized for that model (LC, blue; SN-VTA, plum). Shaded ribbons are within-curve simultaneous 95% confidence bands; pale vertical shading denotes individual zero-excluding intervals lasting at least 20% of the corresponding curve's observed lag span. Shorter intervals remain reported in Supplementary Table 12. Curves without a zero-excluding interval use reduced line opacity (0.35). For measures where lower values indicate better outcomes, y-axes are reversed so that negative coefficients appear above zero; the original coefficient values and signs are retained. A higher association estimate is associated with better outcomes for all variables.

CR, contrast ratio; FDR, false discovery rate; LC, locus coeruleus; SD, standard deviation; SN-VTA, substantia nigra-ventral tegmental area. Outcome abbreviations follow main Fig. 3.

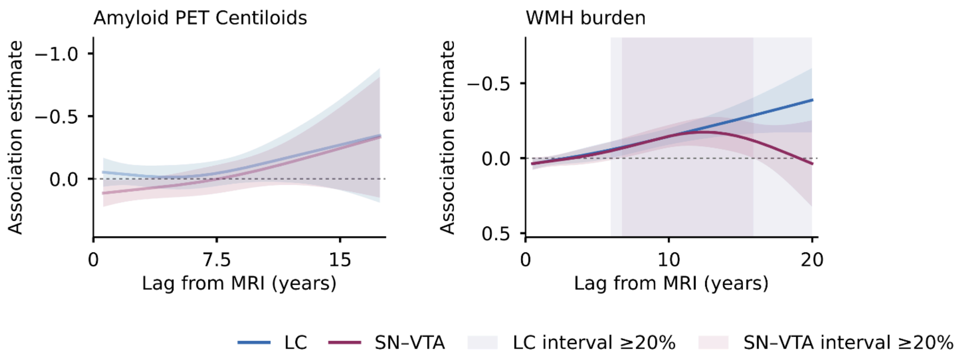


**Supplementary Figure 15. Lag-dependent associations from 180 days after MRI with amyloid PET and white matter hyperintensity burden.** All 38 outcomes are shown across Supplementary Figs. 12-15. Models use assessments at least 180 days after the LC/SN-VTA MRI and are refitted, not cropped primary curves. Lines show the CR main effect plus its lag-dependent smooth per SD of CR as standardized for that model (LC, blue; SN-VTA, plum). Shaded ribbons are within-curve simultaneous 95% confidence bands; pale vertical shading denotes individual zero-excluding intervals lasting at least 20% of the corresponding curve's observed lag span. Shorter intervals remain reported in Supplementary Table 12. Curves without a zero-excluding interval use reduced line opacity (0.35). For measures where lower values indicate better outcomes, y-axes are reversed so that negative coefficients appear above zero; the original coefficient values and signs are retained. A higher association estimate is associated with better outcomes for all variables.

CR, contrast ratio; FDR, false discovery rate; LC, locus coeruleus; SD, standard deviation; SN-VTA, substantia nigra-ventral tegmental area. Outcome abbreviations follow main Fig. 3.

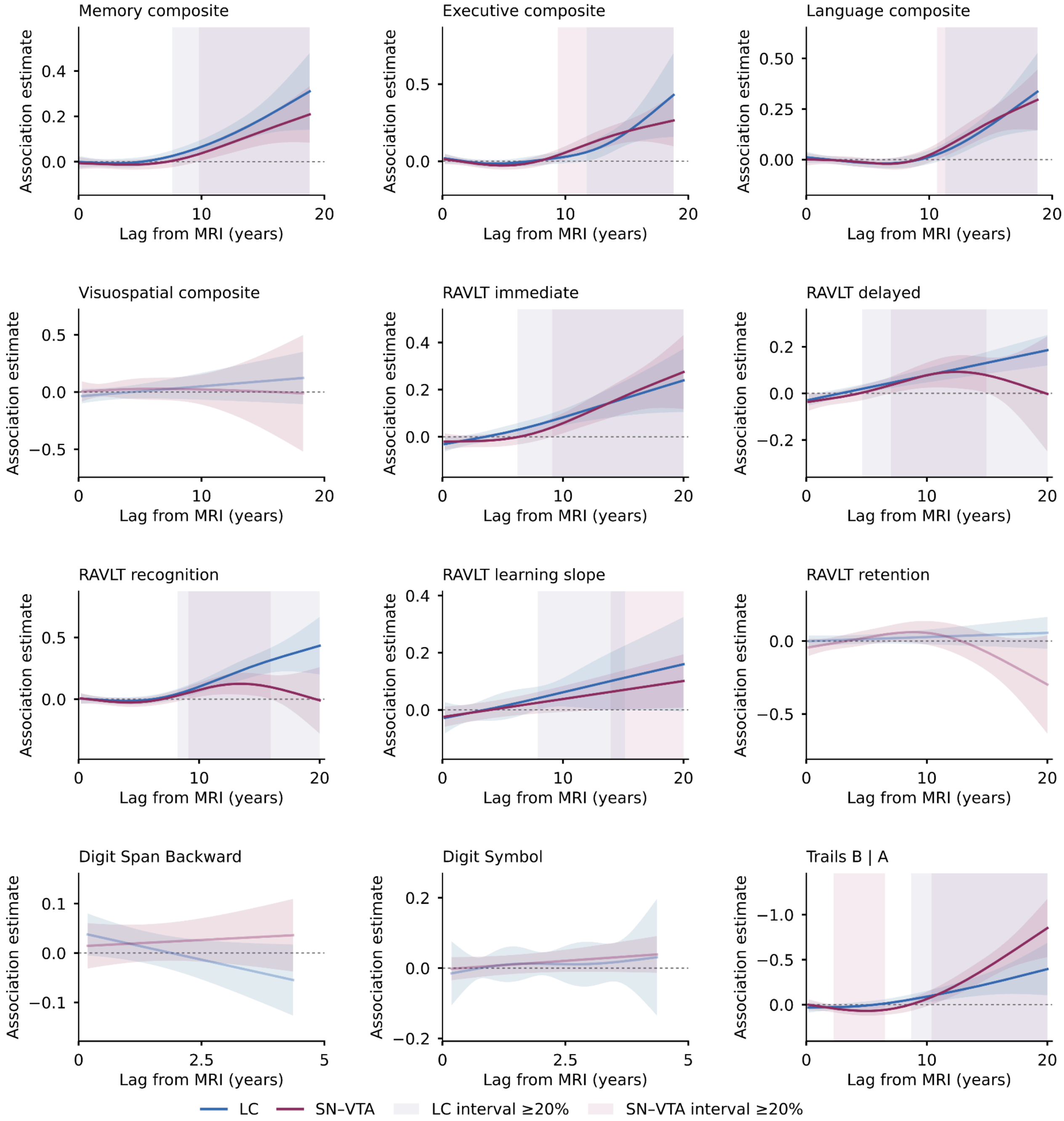


**Supplementary Figure 16. Lag-dependent associations with cognitive composites and cognitive tests in participants with CN or MCI at MRI.** All 38 outcomes are shown across Supplementary Figs. 16-19. Models include participants classified as CN or MCI at the LC/SN-VTA MRI, adjust for that diagnosis and retain later progressors. Lines show the CR main effect plus its lag-dependent smooth per SD of CR as standardized for that model (LC, blue; SN-VTA, plum). Shaded ribbons are within-curve simultaneous 95% confidence bands; pale vertical shading denotes individual zero-excluding intervals lasting at least 20% of the corresponding curve's observed lag span. Shorter intervals remain reported in Supplementary Table 12. Curves without a zero-excluding interval use reduced line opacity (0.35). For measures where lower values indicate better outcomes, y-axes are reversed so that negative coefficients appear above zero; the original coefficient values and signs are retained. A higher association estimate is associated with better outcomes for all variables.

CN, cognitively normal; MCI, mild cognitive impairment. CR, contrast ratio; FDR, false discovery rate; LC, locus coeruleus; SD, standard deviation; SN-VTA, substantia nigra-ventral tegmental area. Outcome abbreviations follow main Fig. 3.

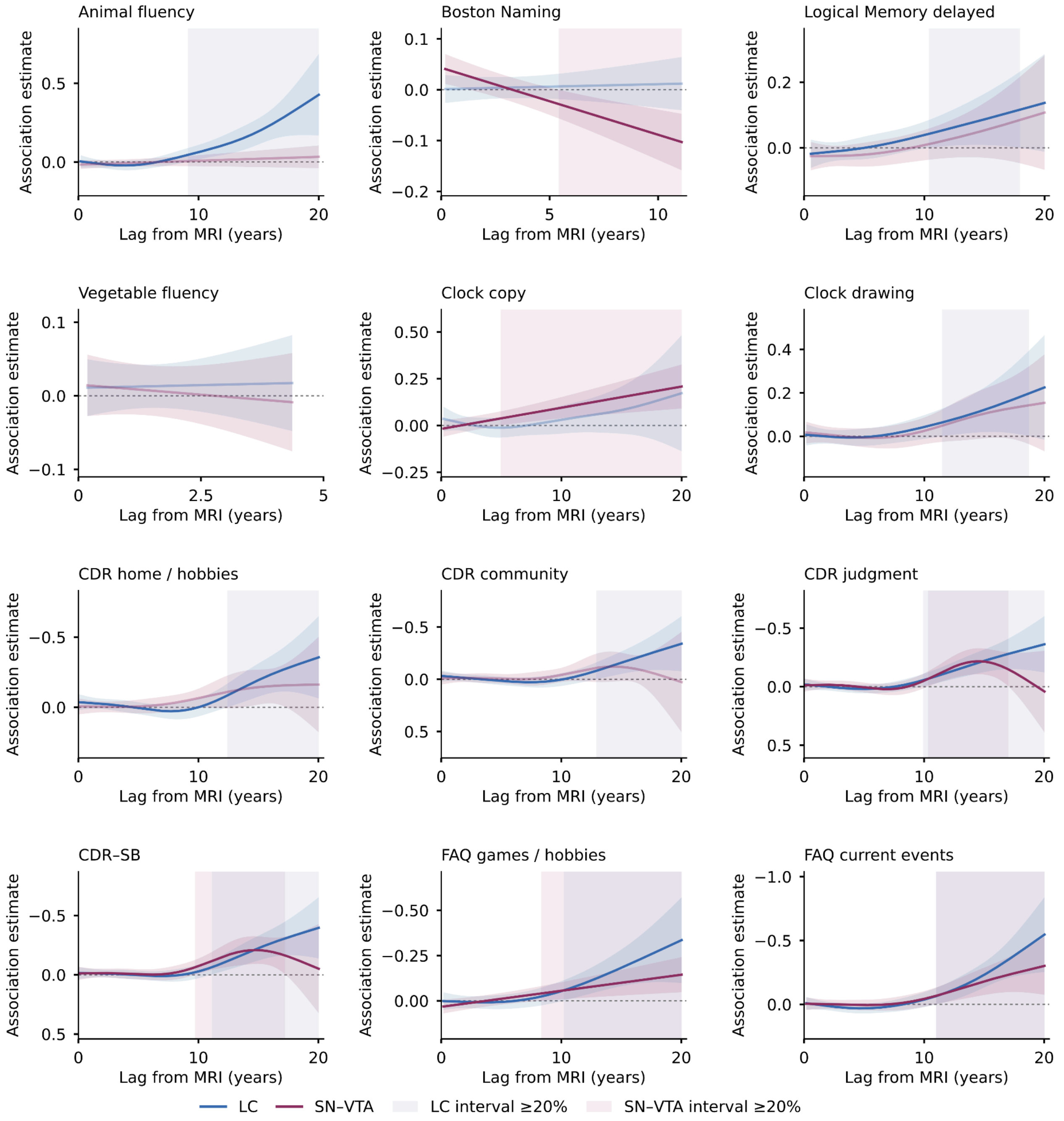


**Supplementary Figure 17. Lag-dependent associations with additional cognitive, clinical and functional measures in participants with CN or MCI at MRI.** All 38 outcomes are shown across Supplementary Figs. 16-19. Models include participants classified as CN or MCI at the LC/SN-VTA MRI, adjust for that diagnosis and retain later progressors. Lines show the CR main effect plus its lag-dependent smooth per SD of CR as standardized for that model (LC, blue; SN-VTA, plum). Shaded ribbons are within-curve simultaneous 95% confidence bands; pale vertical shading denotes individual zero-excluding intervals lasting at least 20% of the corresponding curve's observed lag span. Shorter intervals remain reported in Supplementary Table 12. Curves without a zero-excluding interval use reduced line opacity (0.35). For measures where lower values indicate better outcomes, y-axes are reversed so that negative coefficients appear above zero; the original coefficient values and signs are retained. A higher association estimate is associated with better outcomes for all variables.

CN, cognitively normal; MCI, mild cognitive impairment. CR, contrast ratio; FDR, false discovery rate; LC, locus coeruleus; SD, standard deviation; SN-VTA, substantia nigra-ventral tegmental area. Outcome abbreviations follow main Fig. 3.

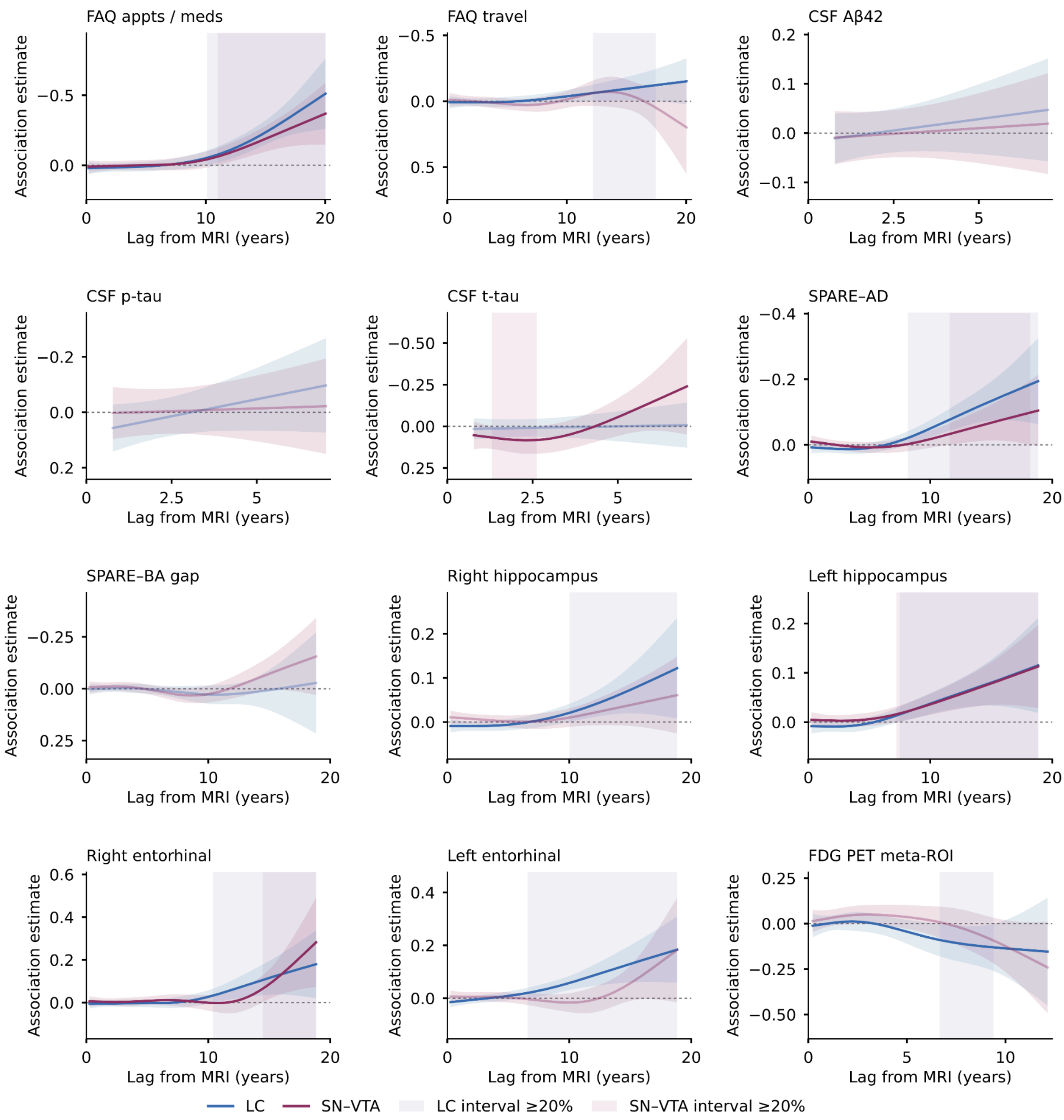

**Supplementary Figure 18. Lag-dependent associations with clinical, functional and biological measures in participants with CN or MCI at MRI.** All 38 outcomes are shown across Supplementary Figs. 16-19. Models include participants classified as CN or MCI at the LC/SN-VTA MRI, adjust for that diagnosis and retain later progressors. Lines show the CR main effect plus its lag-dependent smooth per SD of CR as standardized for that model (LC, blue; SN-VTA, plum). Shaded ribbons are within-curve simultaneous 95% confidence bands; pale vertical shading denotes individual zero-excluding intervals lasting at least 20% of the corresponding curve's observed lag span. Shorter intervals remain reported in Supplementary Table 12. Curves without a zero-excluding interval use reduced line opacity (0.35). For measures where lower values indicate better outcomes, y-axes are reversed so that negative coefficients appear above zero; the original coefficient values and signs are retained. A higher association estimate is associated with better outcomes for all variables.

CN, cognitively normal; MCI, mild cognitive impairment. CR, contrast ratio; FDR, false discovery rate; LC, locus coeruleus; SD, standard deviation; SN-VTA, substantia nigra-ventral tegmental area. Outcome abbreviations follow main Fig. 3.

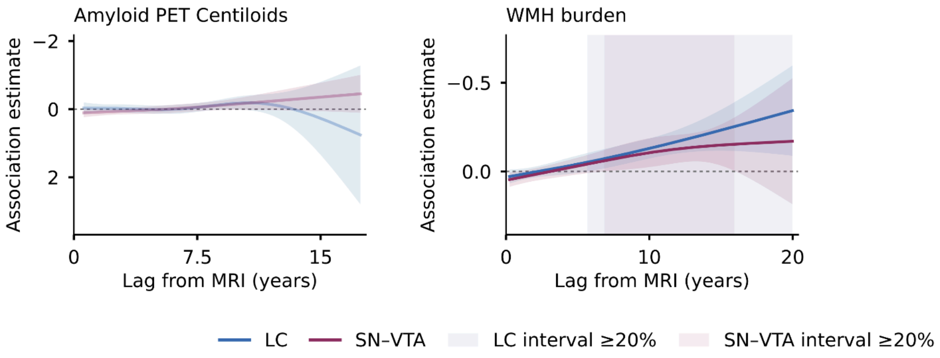


**Supplementary Figure 19. Lag-dependent associations with amyloid PET and white matter hyperintensity burden in participants with CN or MCI at MRI.** All 38 outcomes are shown across Supplementary Figs. 16-19. Models include participants classified as CN or MCI at the LC/SN-VTA MRI, adjust for that diagnosis and retain later progressors. Lines show the CR main effect plus its lag-dependent smooth per SD of CR as standardized for that model (LC, blue; SN-VTA, plum). Shaded ribbons are within-curve simultaneous 95% confidence bands; pale vertical shading denotes individual zero-excluding intervals lasting at least 20% of the corresponding curve's observed lag span. Shorter intervals remain reported in Supplementary Table 12. Curves without a zero-excluding interval use reduced line opacity (0.35). For measures where lower values indicate better outcomes, y-axes are reversed so that negative coefficients appear above zero; the original coefficient values and signs are retained. A higher association estimate is associated with better outcomes for all variables.

CN, cognitively normal; MCI, mild cognitive impairment. CR, contrast ratio; FDR, false discovery rate; LC, locus coeruleus; SD, standard deviation; SN-VTA, substantia nigra-ventral tegmental area. Outcome abbreviations follow main Fig. 3.

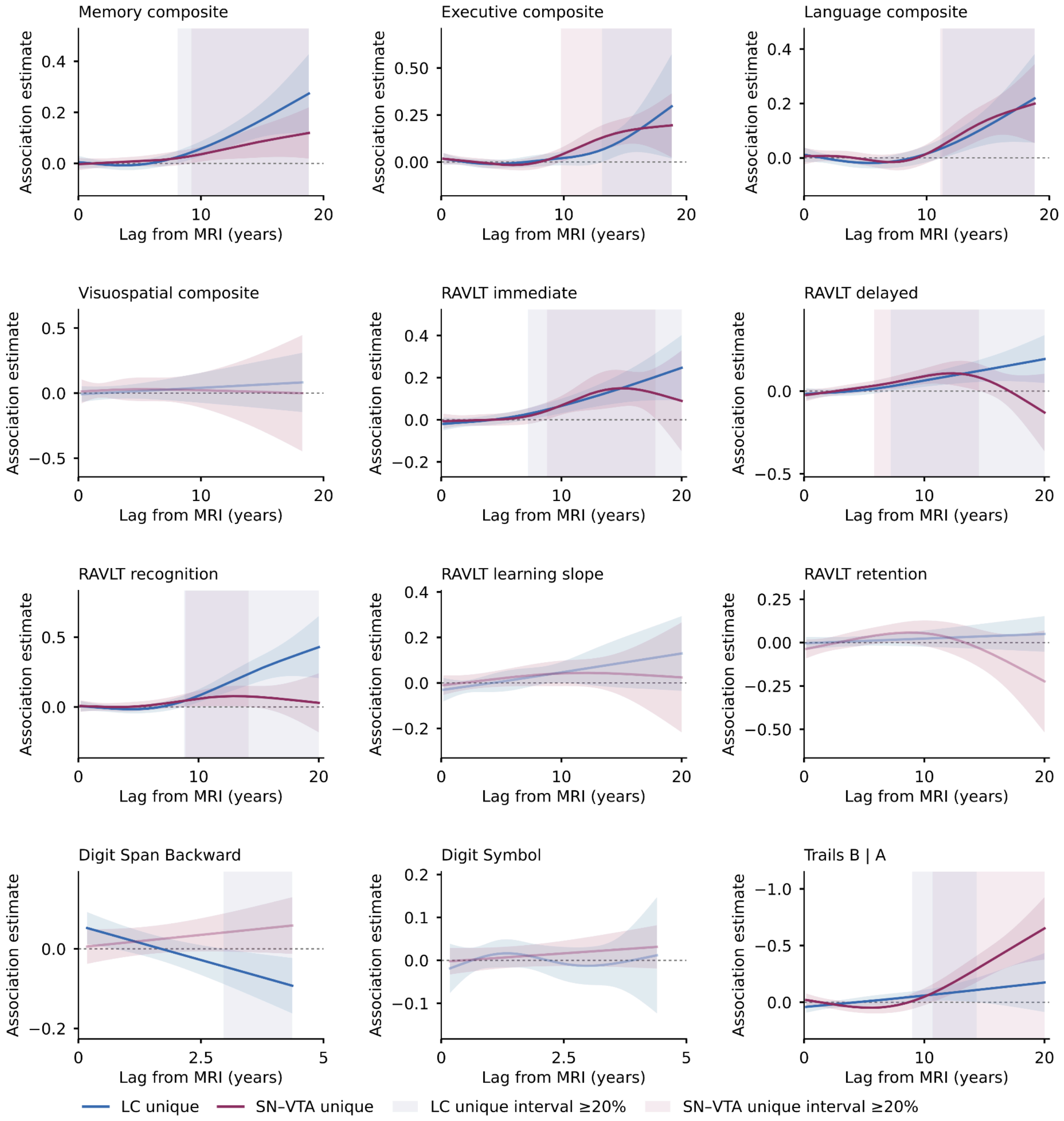


**Supplementary Figure 20. Mutually adjusted LC and SN-VTA lag-dependent associations with cognitive composites and cognitive tests.** All 38 outcomes are shown across Supplementary Figs. 20 -23. LC unique and SN-VTA unique associations are the conditional lag-dependent associations when both CR measures are included. Lines show the CR main effect plus its lag-dependent smooth per SD of CR as standardized for that model (LC, blue; SN-VTA, plum). Shaded ribbons are within-curve simultaneous 95% confidence bands; pale vertical shading denotes individual zero-excluding intervals lasting at least 20% of the corresponding curve's observed lag span. Shorter intervals remain reported in Supplementary Table 12. Curves without a zero-excluding interval use reduced line opacity (0.35). For measures where lower values indicate better outcomes, y-axes are reversed so that negative coefficients appear above zero; the original coefficient values and signs are retained. A higher association estimate is associated with better outcomes for all variables.

CR, contrast ratio; FDR, false discovery rate; LC, locus coeruleus; SD, standard deviation; SN-VTA, substantia nigra-ventral tegmental area. Outcome abbreviations follow main Fig. 3.

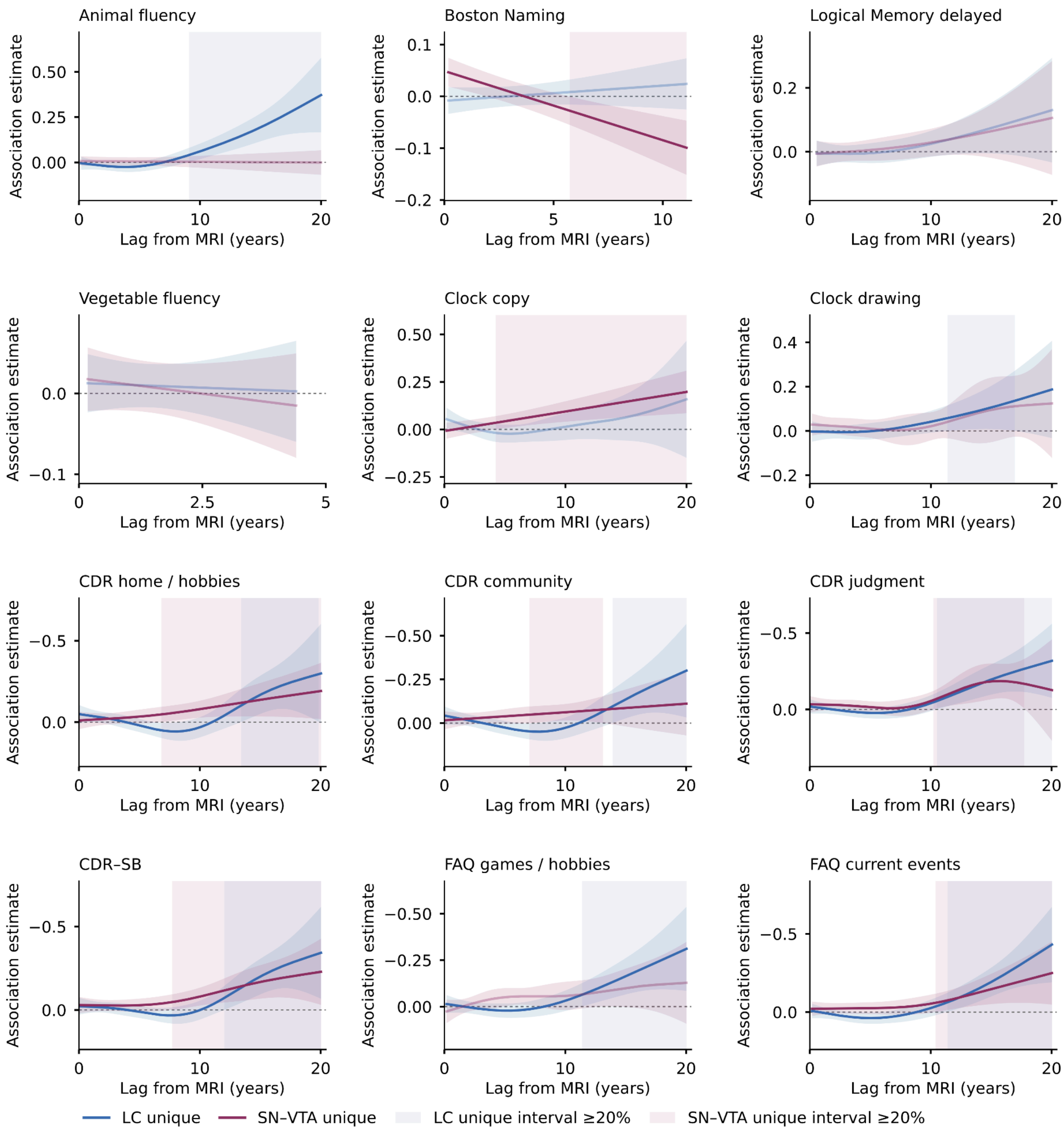

**Supplementary Figure 21. Mutually adjusted LC and SN-VTA lag-dependent associations with additional cognitive, clinical and functional measures.** All 38 outcomes are shown across Supplementary Figs. 20-23. LC unique and SN-VTA unique are the conditional lag-dependent associations when both CR measures are included. Lines show the CR main effect plus its lag-dependent smooth per SD of CR as standardized for that model (LC, blue; SN-VTA, plum). Shaded ribbons are within-curve simultaneous 95% confidence bands; pale vertical shading denotes individual zero-excluding intervals lasting at least 20% of the corresponding curve's observed lag span. Shorter intervals remain reported in Supplementary Table 12. Curves without a zero-excluding interval use reduced line opacity (0.35). For measures where lower values indicate better outcomes, y-axes are reversed so that negative coefficients appear above zero; the original coefficient values and signs are retained. A higher association estimate is associated with better outcomes for all variables.

CR, contrast ratio; FDR, false discovery rate; LC, locus coeruleus; SD, standard deviation; SN-VTA, substantia nigra-ventral tegmental area. Outcome abbreviations follow main Fig. 3.

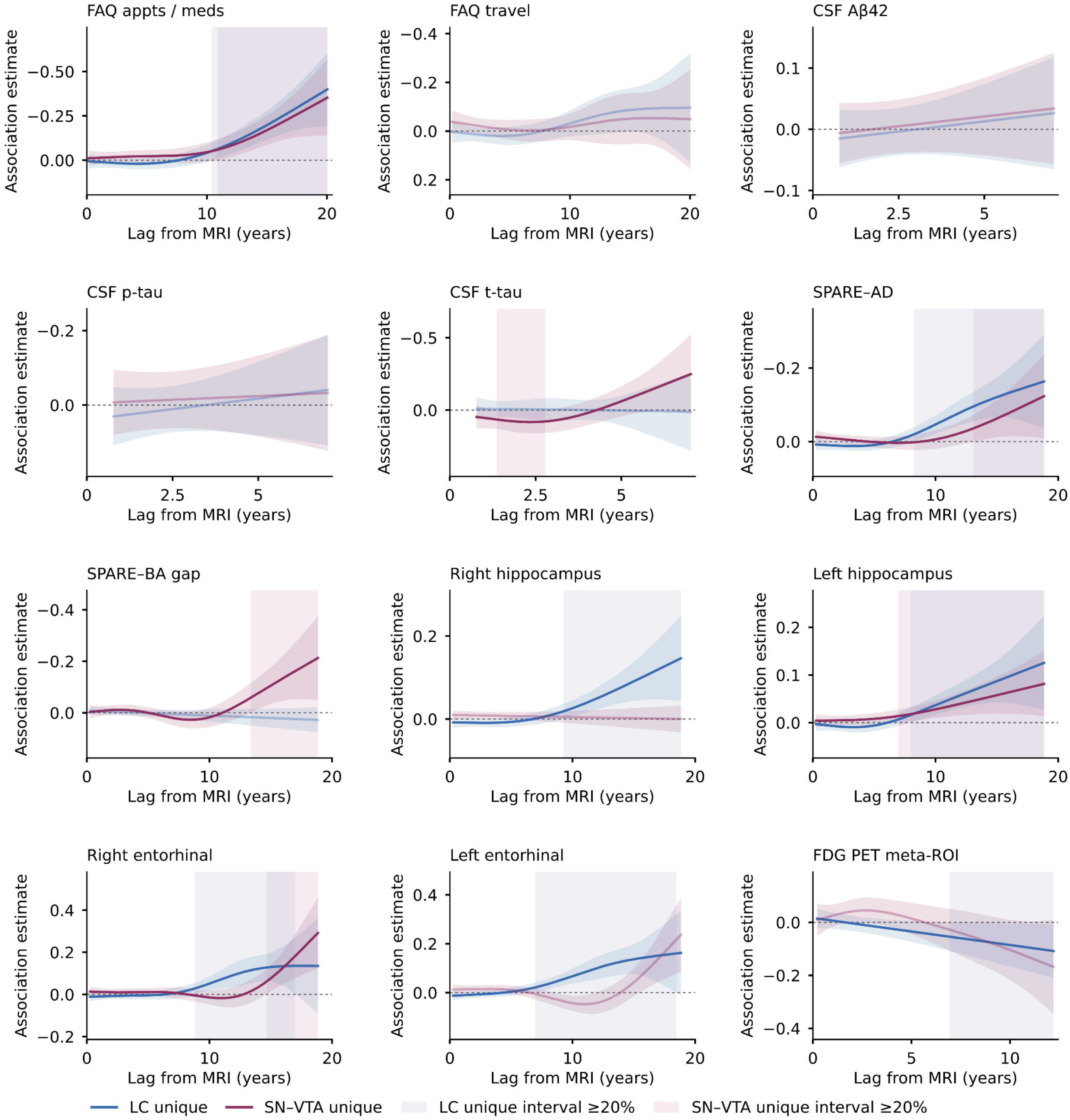


**Supplementary Figure 22. Mutually adjusted LC and SN-VTA lag-dependent associations with clinical, functional and biological measures.** All 38 outcomes are shown across Supplementary Figs. 20 -23. LC unique and SN-VTA unique are the conditional lag-dependent associations when both CR measures are included. Lines show the CR main effect plus its lag-dependent smooth per SD of CR as standardized for that model (LC, blue; SN-VTA, plum). Shaded ribbons are within-curve simultaneous 95% confidence bands; pale vertical shading denotes individual zero-excluding intervals lasting at least 20% of the corresponding curve's observed lag span. Shorter intervals remain reported in Supplementary Table 12. Curves without a zero-excluding interval use reduced line opacity (0.35). For measures where lower values indicate better outcomes, y-axes are reversed so that negative coefficients appear above zero; the original coefficient values and signs are retained. A higher association estimate is associated with better outcomes for all variables.

CR, contrast ratio; FDR, false discovery rate; LC, locus coeruleus; SD, standard deviation; SN-VTA, substantia nigra-ventral tegmental area. Outcome abbreviations follow main Fig. 3.

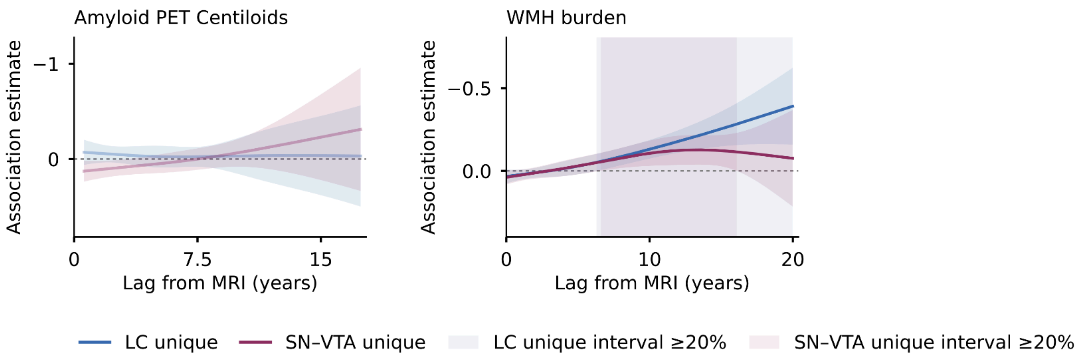


**Supplementary Figure 23. Mutually adjusted LC and SN-VTA lag-dependent associations with amyloid PET and white matter hyperintensity burden.** All 38 outcomes are shown across Supplementary Figs. 20 -23. LC unique and SN-VTA unique are the conditional lag-dependent associations when both CR measures are included. Lines show the CR main effect plus its lag-dependent smooth per SD of CR as standardized for that model (LC, blue; SN-VTA, plum). Shaded ribbons are within-curve simultaneous 95% confidence bands; pale vertical shading denotes individual zero-excluding intervals lasting at least 20% of the corresponding curve's observed lag span. Shorter intervals remain reported in Supplementary Table 12. Curves without a zero-excluding interval use reduced line opacity (0.35). For measures where lower values indicate better outcomes, y-axes are reversed so that negative coefficients appear above zero; the original coefficient values and signs are retained. A higher association estimate is associated with better outcomes for all variables.

CR, contrast ratio; FDR, false discovery rate; LC, locus coeruleus; SD, standard deviation; SN-VTA, substantia nigra-ventral tegmental area. Outcome abbreviations follow main Fig. 3.

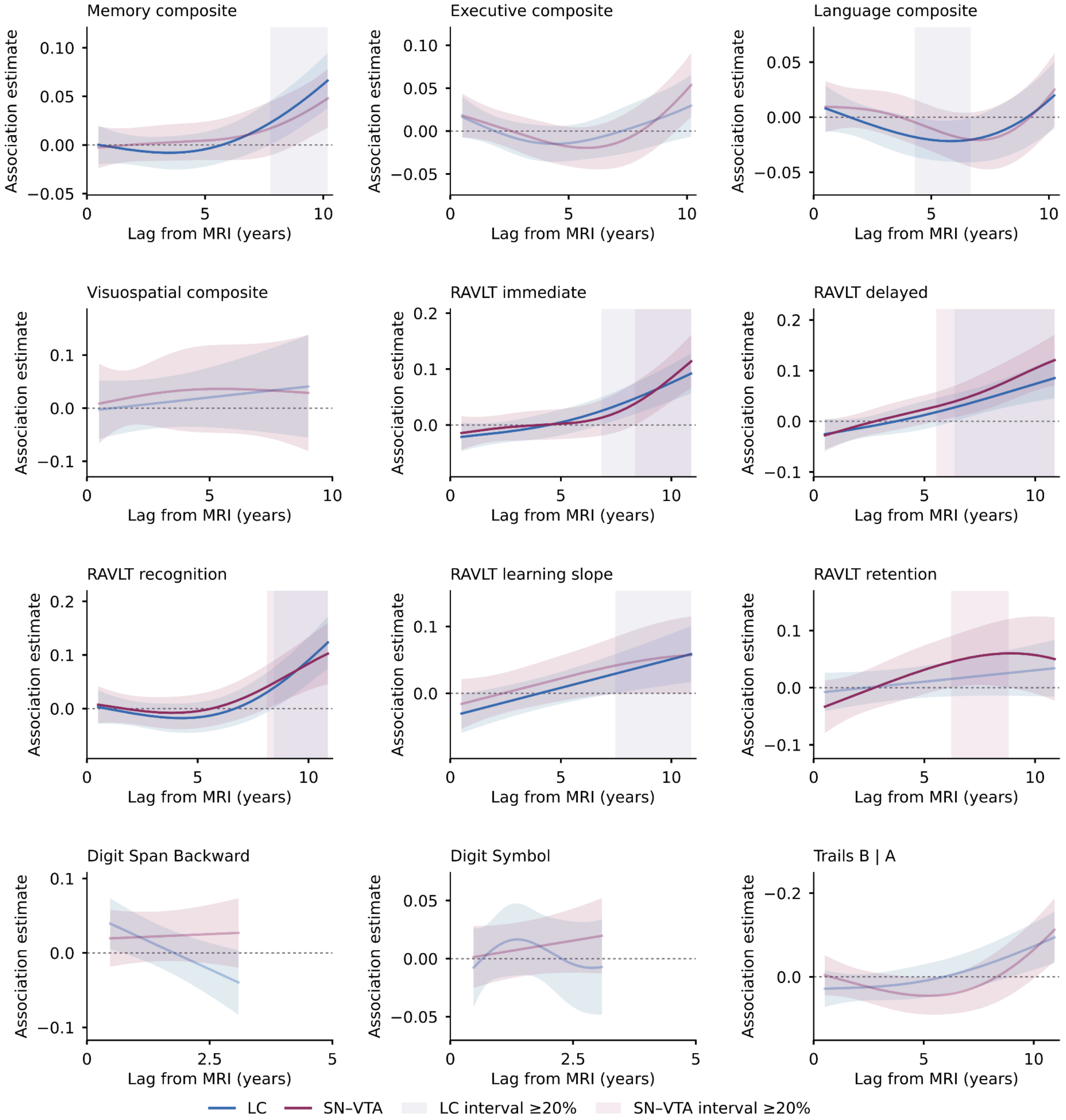


**Supplementary Figure 24. Lag models over 5th-95th percentile lag range: cognitive composites and cognitive tests.** All 38 outcomes are shown across Supplementary Figs. 24-27. LC and SN-VTA curves retain 5th-95th-percentile evaluation ranges and corresponding simultaneous confidence bands. Lines show the CR main effect plus its lag-dependent smooth per SD of CR as standardized for that model (LC, blue; SN-VTA, plum). Shaded ribbons are within-curve simultaneous 95% confidence bands; pale vertical shading denotes individual zero-excluding intervals lasting at least 20% of the corresponding curve's observed lag span. Shorter intervals remain reported in Supplementary Table 12. Curves without a zero-excluding interval use reduced line opacity (0.35). For measures where lower values indicate better outcomes, y-axes are reversed so that negative coefficients appear above zero; the original coefficient values and signs are retained. A higher association estimate is associated with better outcomes for all variables.

CR, contrast ratio; FDR, false discovery rate; LC, locus coeruleus; SD, standard deviation; SN-VTA, substantia nigra-ventral tegmental area. Outcome abbreviations follow main Fig. 3.

.

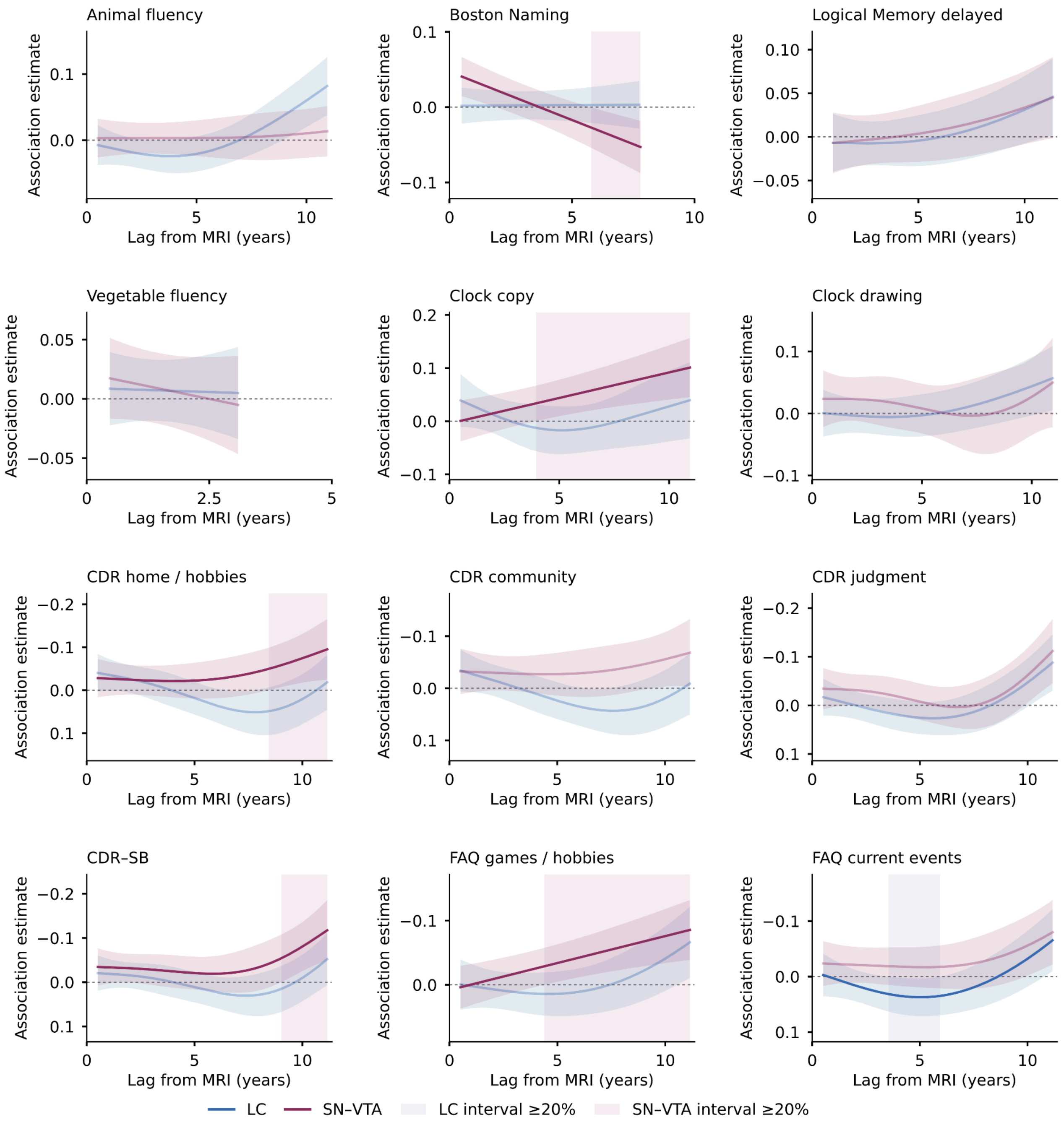


**Supplementary Figure 25. Lag models over 5th-95th percentile lag range: additional cognitive, clinical and functional measures.** All 38 outcomes are shown across Supplementary Figs. 24-27. LC and SN-VTA curves retain 5th-95th-percentile evaluation ranges and corresponding simultaneous confidence bands. Lines show the CR main effect plus its lag-dependent smooth per SD of CR as standardized for that model (LC, blue; SN-VTA, plum). Shaded ribbons are within-curve simultaneous 95% confidence bands; pale vertical shading denotes individual zero-excluding intervals lasting at least 20% of the corresponding curve's observed lag span. Shorter intervals remain reported in Supplementary Table 12. Curves without a zero-excluding interval use reduced line opacity (0.35). For measures where lower values indicate better outcomes, y-axes are reversed so that negative coefficients appear above zero; the original coefficient values and signs are retained. A higher association estimate is associated with better outcomes for all variables.

CR, contrast ratio; FDR, false discovery rate; LC, locus coeruleus; SD, standard deviation; SN-VTA, substantia nigra-ventral tegmental area. Outcome abbreviations follow main Fig. 3.

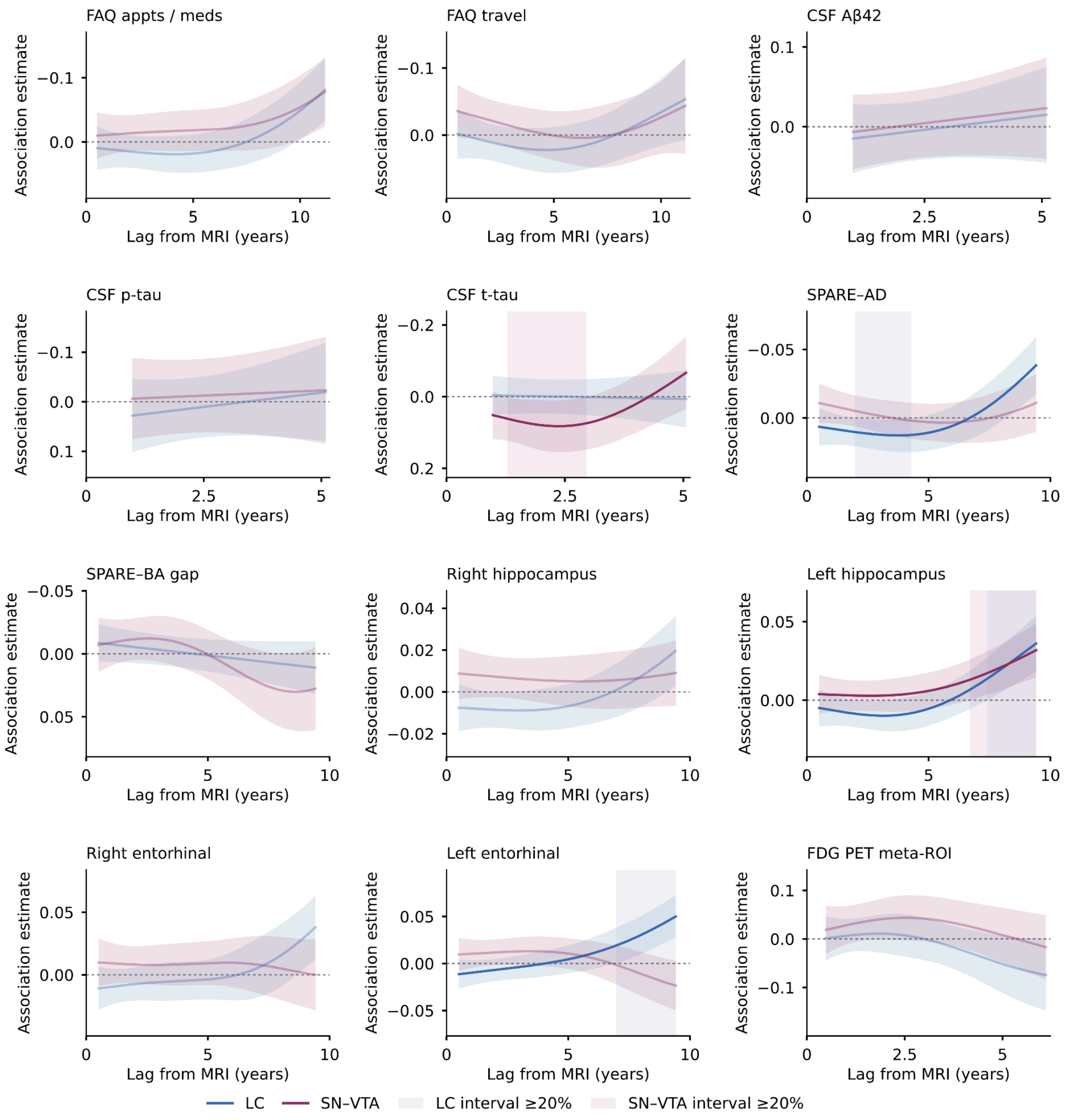


**Supplementary Figure 26. Lag models over 5th-95th percentile lag range: clinical, functional and biological measures.** All 38 outcomes are shown across Supplementary Figs. 24-27. LC and SN-VTA curves retain 5th-95th-percentile evaluation ranges and corresponding simultaneous confidence bands. Lines show the CR main effect plus its lag-dependent smooth per SD of CR as standardized for that model (LC, blue; SN-VTA, plum). Shaded ribbons are within-curve simultaneous 95% confidence bands; pale vertical shading denotes individual zero-excluding intervals lasting at least 20% of the corresponding curve's observed lag span. Shorter intervals remain reported in Supplementary Table 12. Curves without a zero-excluding interval use reduced line opacity (0.35). For measures where lower values indicate better outcomes, y-axes are reversed so that negative coefficients appear above zero; the original coefficient values and signs are retained. A higher association estimate is associated with better outcomes for all variables.

CR, contrast ratio; FDR, false discovery rate; LC, locus coeruleus; SD, standard deviation; SN-VTA, substantia nigra-ventral tegmental area. Outcome abbreviations follow main Fig. 3.

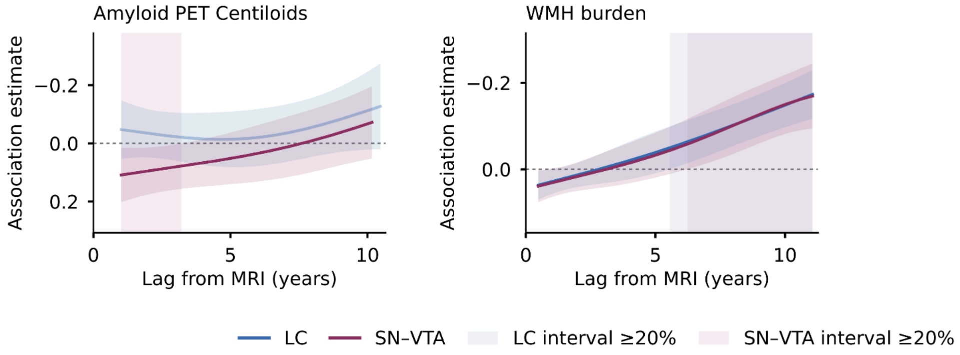


**Supplementary Figure 27. Lag models over 5th-95th percentile lag range: amyloid PET and white matter hyperintensity burden.** All 38 outcomes are shown across Supplementary Figs. 24-27. LC and SN-VTA curves retain 5th-95th-percentile evaluation ranges and corresponding simultaneous confidence bands. Lines show the CR main effect plus its lag-dependent smooth per SD of CR as standardized for that model (LC, blue; SN-VTA, plum). Shaded ribbons are within-curve simultaneous 95% confidence bands; pale vertical shading denotes individual zero-excluding intervals lasting at least 20% of the corresponding curve's observed lag span. Shorter intervals remain reported in Supplementary Table 12. Curves without a zero-excluding interval use reduced line opacity (0.35). For measures where lower values indicate better outcomes, y-axes are reversed so that negative coefficients appear above zero; the original coefficient values and signs are retained. A higher association estimate is associated with better outcomes for all variables.

CR, contrast ratio; FDR, false discovery rate; LC, locus coeruleus; SD, standard deviation; SN-VTA, substantia nigra-ventral tegmental area. Outcome abbreviations follow main Fig. 3.

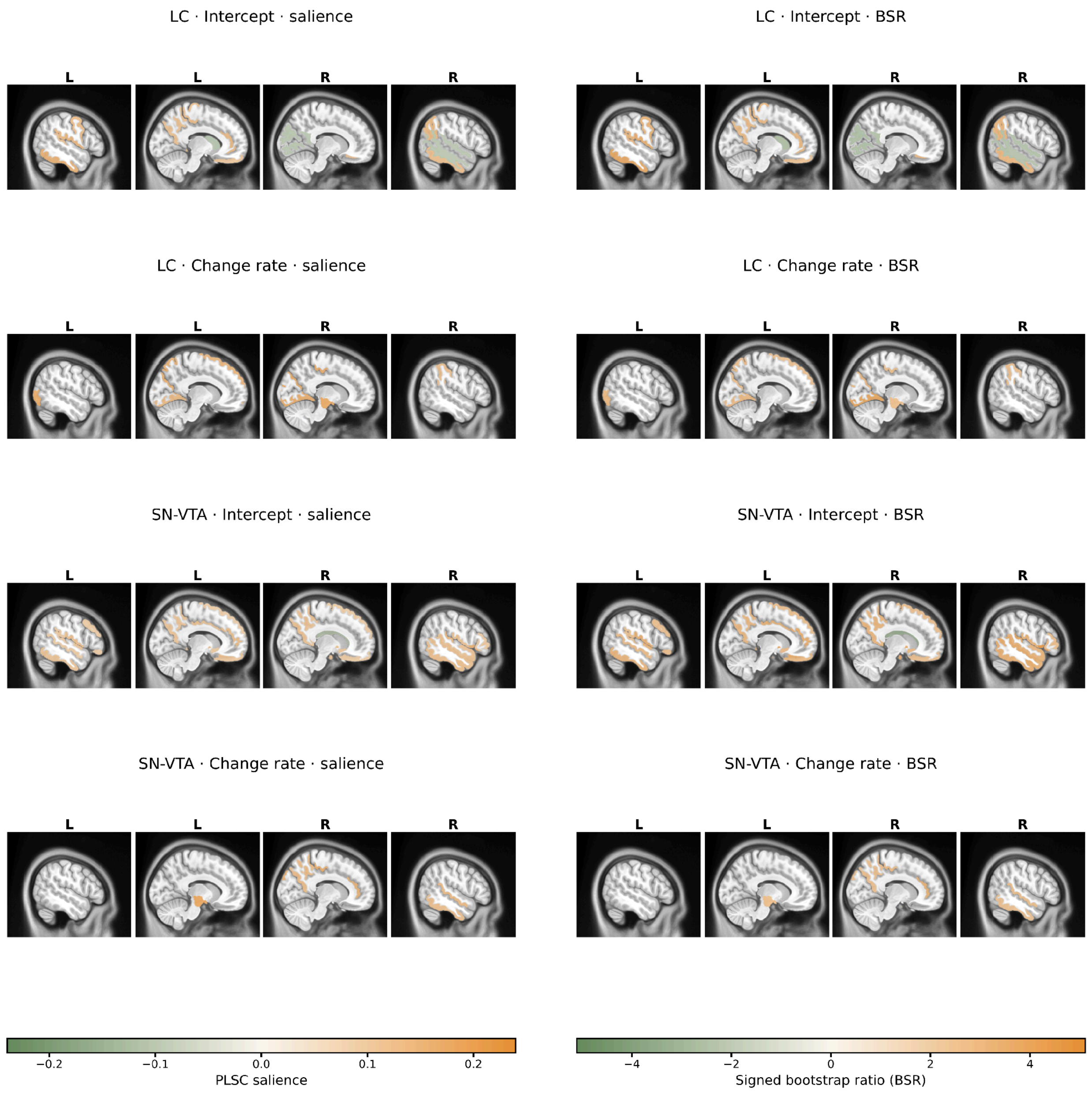


**Supplementary Figure 28**. Direct PLSC saliences and signed bootstrap ratios at |BSR| ≥ 2.56. Rows show the LC intercept, LC rate-of-change, SN-VTA intercept and SN-VTA rate-of-change patterns, respectively. The left column shows saliences and the right column BSRs on matched anatomical views. Both display regions satisfying |BSR| ≥ 2.56. Green indicates negative values and orange positive values; separate color bars indicate the salience and BSR scales.

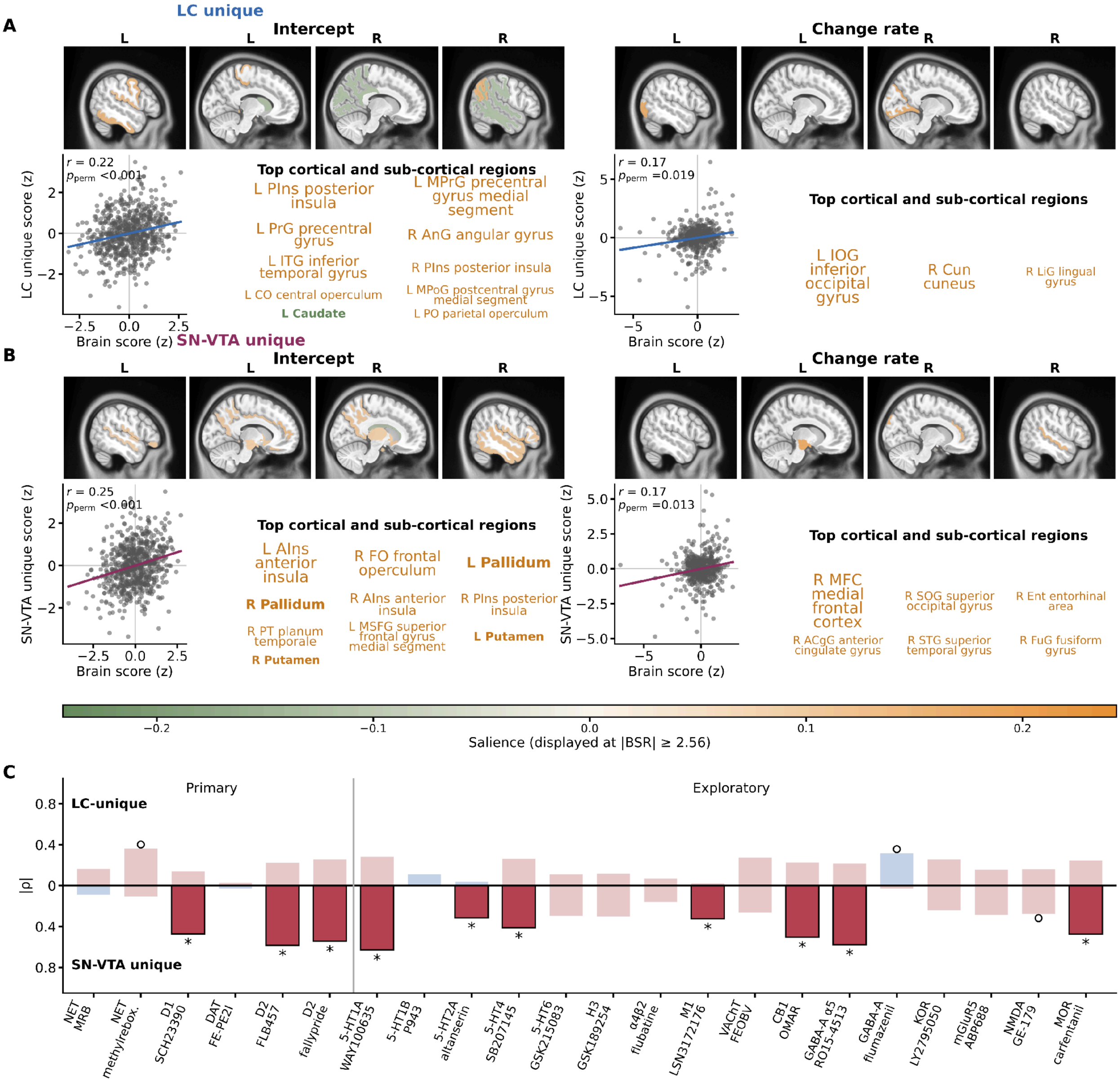


**Supplementary Figure 29. Unique structural patterns and molecular correspondence.** a,b, LC and SN-VTA intercept and rate-of-change patterns after residualizing the corresponding CR trajectory parameter against the other region's parameter. Brain maps show saliences for regions satisfying |BSR| ≥ 2.56; green indicates negative and orange positive salience. Scatterplots show participant brain and residual-CR scores with descriptive fitted lines; r is their correlation and P refers to the global permutation test. c, Spearman correspondence between unique intercept patterns and 22 PET maps. Upper/lower bars represent LC/SN-VTA rather; bar length represents |ρ|, and red/blue denotes positive/negative correspondence. Asterisks identify $P_{FDR}$ < 0.05; open circles identify P < 0.05 without meeting the FDR criterion.

BSR, bootstrap ratio; CB1, cannabinoid receptor 1; CR, contrast ratio; D1, dopamine D1-like receptor; D2, dopamine D2-family receptor; DAT, dopamine transporter; FDR, false discovery rate; 5-HT, 5-hydroxytryptamine/serotonin; GABA$_A$, γ-aminobutyric acid type A receptor; H3, histamine H3 receptor;

KOR, κ-opioid receptor; LC, locus coeruleus; LV, latent variable; mGluR5, metabotropic glutamate receptor 5; M1, muscarinic M1 receptor; MOR, μ-opioid receptor; VAChT, vesicular acetylcholine transporter. MUSE, multi-atlas region segmentation utilizing ensembles; NET, norepinephrine transporter; NMDA, N-methyl-D-aspartate receptor; PET, positron emission tomography; PLSC, partial least squares correlation; SN-VTA, substantia nigra-ventral tegmental area.

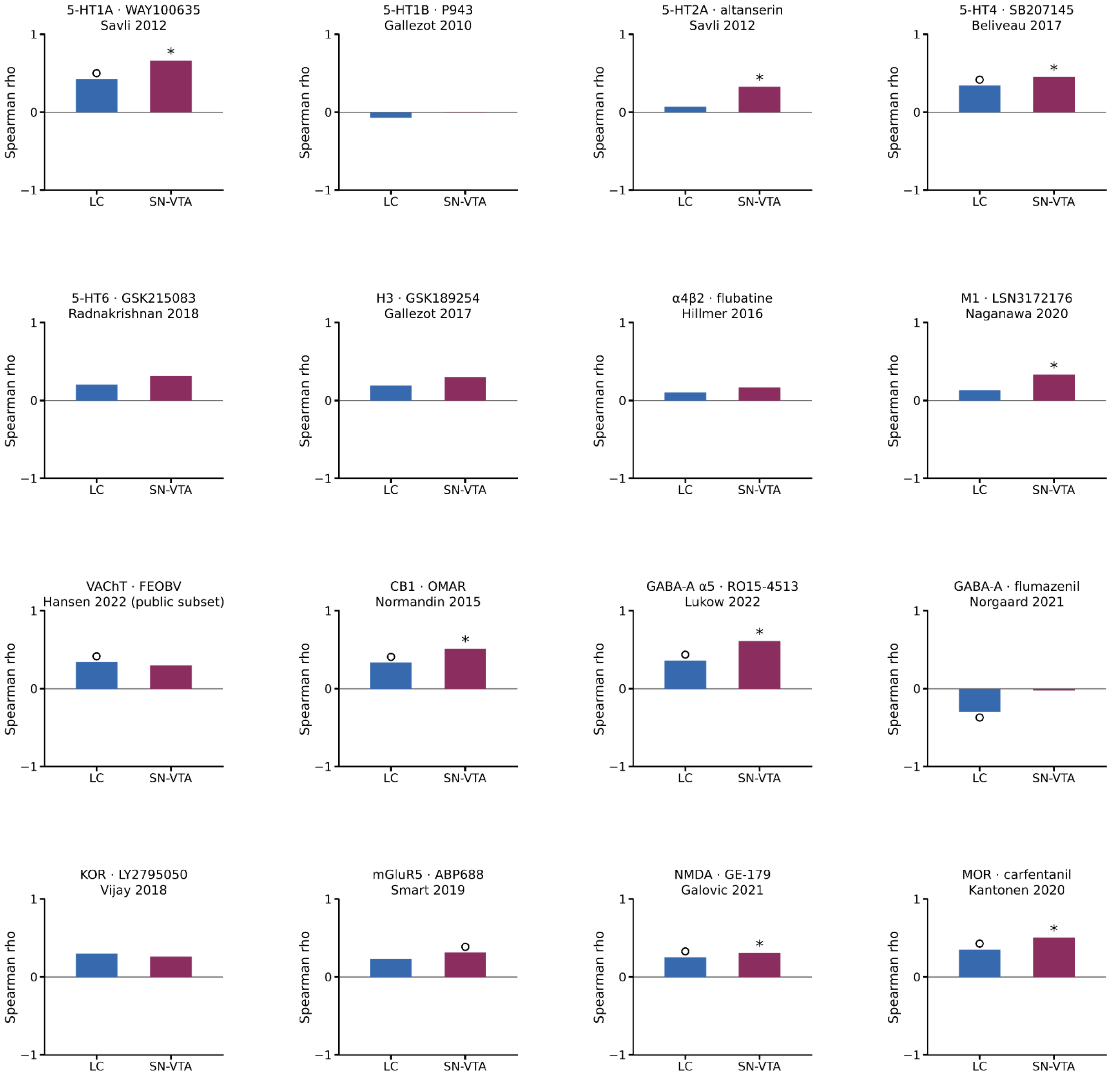


**Supplementary Figure 30.** Exploratory PET correspondence of intercept patterns. Each panel shows one of the 16 exploratory PET maps. Bars show Spearman correlations with the cortical LC and SN-VTA intercept saliences; blue identifies LC and plum SN-VTA. Asterisks indicate $P_{FDR} < 0.05$; hollow circles indicate nominal P < 0.05 without meeting the FDR correction. Two-sided spatial probabilities were estimated from 10,000 Moran-spectral surrogate maps. Complete signed correlations and probabilities are reported in Supplementary Table 17. PET-target abbreviations follow Supplementary Fig. 29.

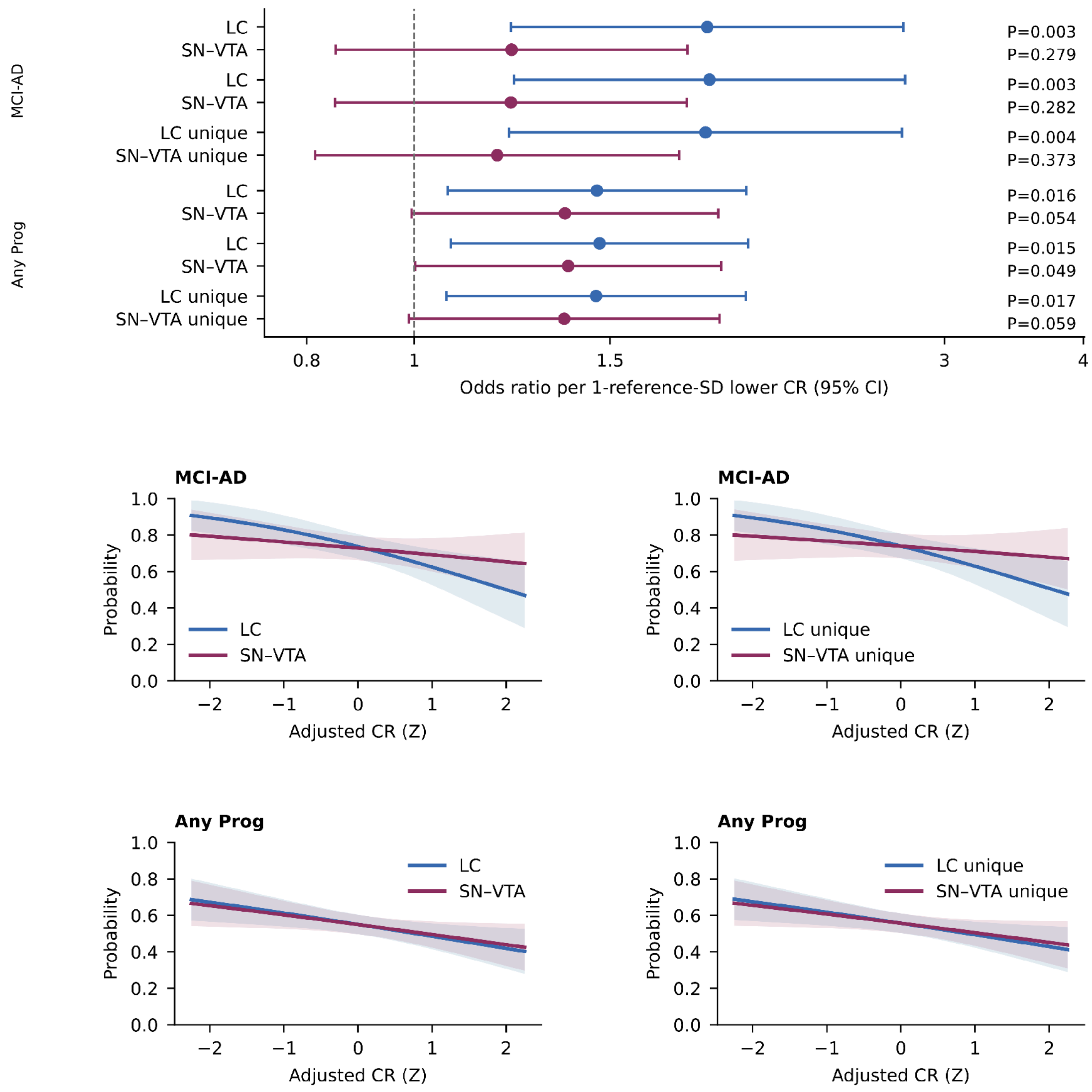


**Supplementary Figure 31. Clinical progression with LC and SN-VTA included in the same model.** The upper panel shows odds ratios and 95% confidence intervals per one SD lower CR for MCI-to-AD and any progression. LC unique and SN-VTA unique are the conditional coefficients in the model containing both CR measures. Blue identifies LC and plum SN-VTA. Displayed P values are unadjusted. Lower panels show five-year probabilities and 95% confidence intervals for the separate model (left) and models containing both CR measures (right) (Supplementary Table 18).

CR, contrast ratio; LC, locus coeruleus; MCI-AD, mild cognitive impairment to AD dementia; Any Prog, any clinical progression; SN-VTA, substantia nigra-ventral tegmental area.

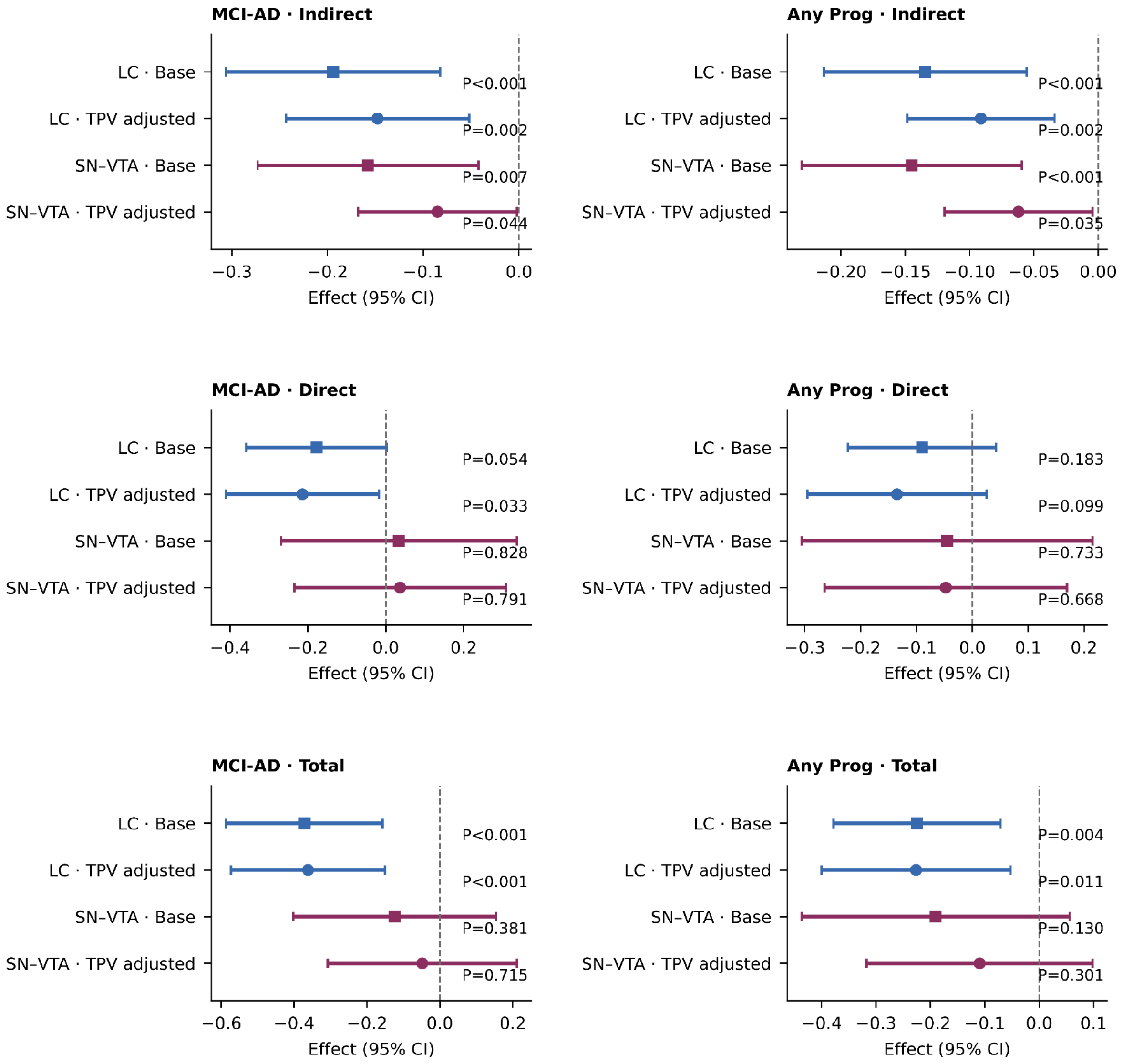


**Supplementary Figure 32. Mediation effects for five-year clinical progression.**

Top, middle, and bottom rows show indirect, direct, and total effects, respectively; columns show MCI-to-AD (left) and any progression (right). Blue and plum identify LC and SN-VTA. Squares indicate models without TPV and circles indicate models with TPV. Bars are 95% confidence intervals. Annotated P values and full effects are reported in Supplementary Table 20. The model was specified in the main manuscript (see Methods, Associational indirect effects through distributed structural brain patterns section).

LC, locus coeruleus; SEM, structural equation model; SN-VTA, substantia nigra-ventral tegmental area; TPV, total parenchymal volume.

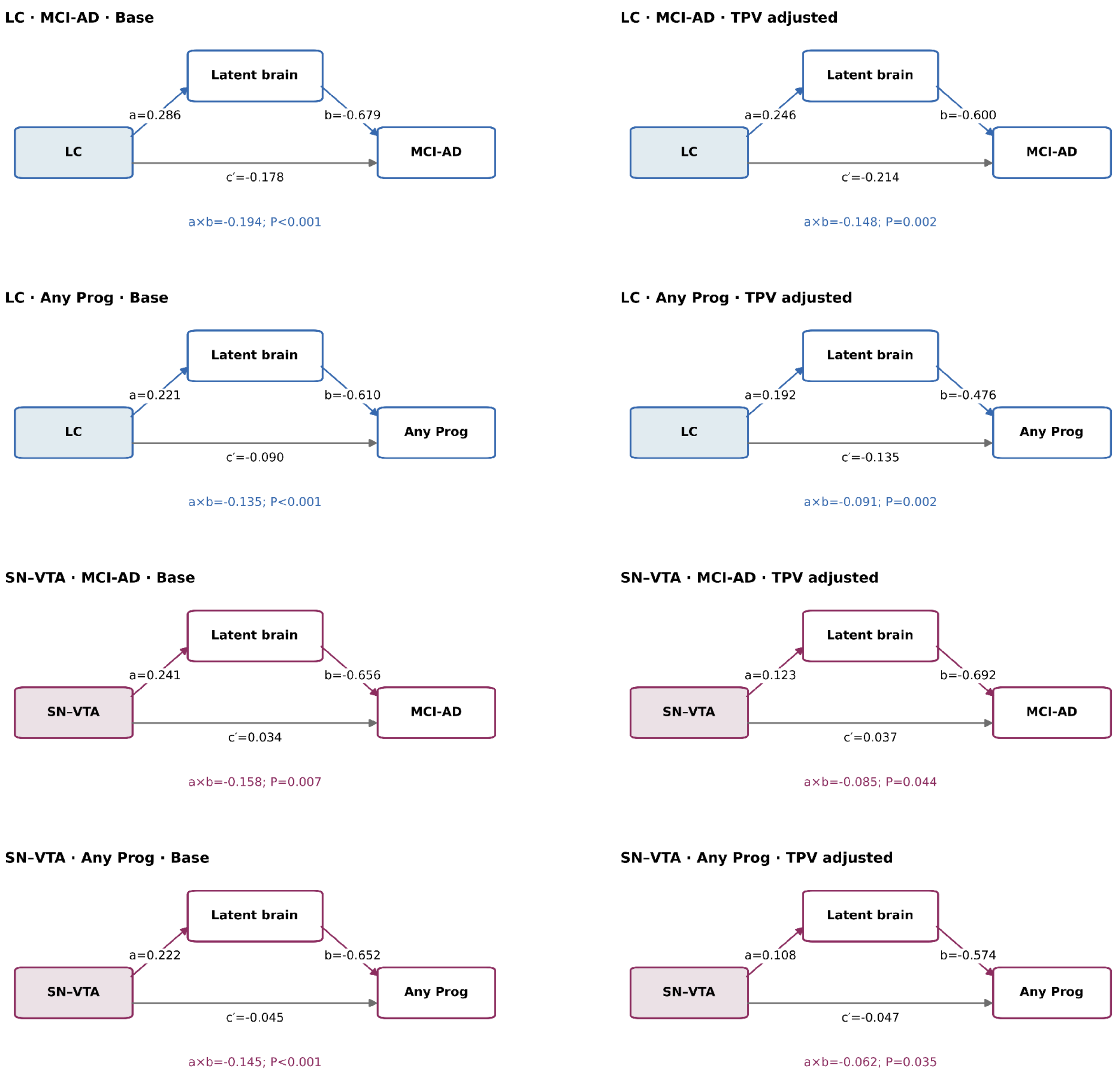


**Supplementary Figure 33. Mediation pathways for five-year clinical progression.**

Rows show LC MCI-to-AD, LC any progression, SN-VTA MCI-to-AD and SN-VTA any progression; columns compare models without TPV (left) and with TPV (right).

Path labels show fitted coefficients; the indirect, direct and total effects and their probabilities are reported in Supplementary Table 20.

CR, contrast ratio; LC, locus coeruleus; MCI-AD, mild cognitive impairment to AD; Any Prog, any clinical progression; PLSC, partial least squares correlation; SN-VTA, substantia nigra-ventral tegmental area; TPV, total parenchymal volume.

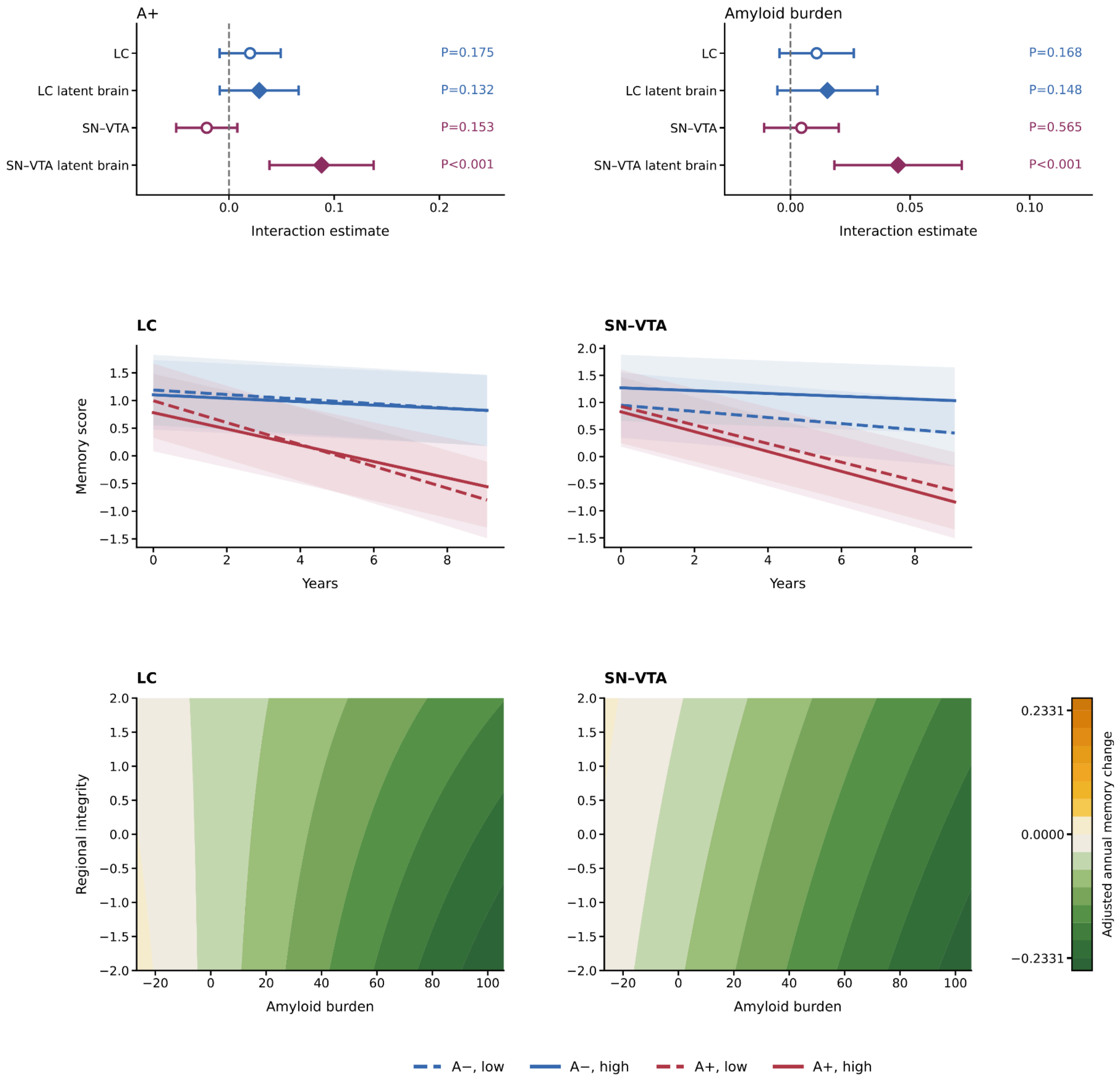


**Supplementary Figure 34. LC and SN-VTA CR interactions with amyloid-related memory change.** Upper panels show CR (open circles) and brain-pattern (filled diamonds) interactions with amyloid and time, with 95% confidence intervals and P values. Middle panels show predicted memory at low/high LC or SN-VTA CR (−1/+1 model SD; dashed/solid lines); green/orange identifies A−/A+. Bands are 95% confidence intervals. Lower panels show adjusted annual memory change across Centiloids and CR. “Regional integrity” is standardized bilateral CR.

A−/A+, amyloid negative/positive; CR, contrast ratio; LC, locus coeruleus; SD, standard deviation; SN-VTA, substantia nigra-ventral tegmental area.

### Supplementary Tables

**Supplementary Table 1. Sample characteristics**

| Characteristic | Overall | Any 1.5-T data | Any 3-T data | Both field strengths |
|---|---|---|---|---|
| Participants, n | 739 | 710 | 145 | 116 |
| Reconciled MRI scan sets, n | 3,782 | 3,273 | 509 | 848 |
| Baseline age, years | 75.42 (6.89) n=720; missing 19 | 75.44 (6.90) n=691; missing 19 | 75.01 (7.11) n=145; missing 0 | 75.00 (7.24) n=116; missing 0 |
| Education, years | 15.54 (3.05) n=737; missing 2 | 15.51 (3.07) n=708; missing 2 | 15.70 (2.95) n=144; missing 1 | 15.57 (3.08) n=115; missing 1 |
| Female sex, n/available N (%) | 321/739 (43.4%) 0 missing | 310/710 (43.7%) 0 missing | 74/145 (51.0%) 0 missing | 63/116 (54.3%) 0 missing |
| APOE ε4 carrier, n/available N (%) | 367/739 (49.7%) 0 missing | 349/710 (49.2%) 0 missing | 71/145 (49.0%) 0 missing | 53/116 (45.7%) 0 missing |
| Baseline diagnosis: CN | 204/720 (28.3%) 19 missing | 196/691 (28.4%) 19 missing | 46/145 (31.7%) 0 missing | 38/116 (32.8%) 0 missing |
| Baseline diagnosis: MCI | 350/720 (48.6%) 19 missing | 335/691 (48.5%) 19 missing | 69/145 (47.6%) 0 missing | 54/116 (46.6%) 0 missing |
| Baseline diagnosis: AD | 166/720 (23.1%) 19 missing | 160/691 (23.2%) 19 missing | 30/145 (20.7%) 0 missing | 24/116 (20.7%) 0 missing |
| Baseline diagnosis: other | 0/720 (0.0%) 19 missing | 0/691 (0.0%) 19 missing | 0/145 (0.0%) 0 missing | 0/116 (0.0%) 0 missing |
| MRI scan sets per participant | 5.12 (2.90) n=739; missing 0 | 4.61 (2.63) n=710; missing 0 | 3.51 (1.41) n=145; missing 0 | 7.31 (3.33) n=116; missing 0 |
| Participants with repeated MRI | 663/739 (89.7%) 0 missing | 611/710 (86.1%) 0 missing | 131/145 (90.3%) 0 missing | 116/116 (100.0%) 0 missing |

| Characteristic | Overall | Any 1.5-T data | Any 3-T data | Both field strengths |
|---|---|---|---|---|
| Mean consecutive MRI interval, years | 0.72 (0.29) n=663; missing 0 | 0.82 (0.37) n=611; missing 0 | 0.77 (0.38) n=131; missing 0 | 0.51 (0.30) n=116; missing 0 |
| First-to-last MRI span, years | 3.39 (2.42) n=663; missing 0 | 3.50 (2.45) n=611; missing 0 | 1.99 (0.86) n=131; missing 0 | 3.52 (2.84) n=116; missing 0 |
| First-to-last MRI span, range (years) | 0.04-10.29 | 0.36-10.29 | 0.29-3.27 | 0.04-10.29 |

**Supplementary Table 2. ICC**

| Region | Field strength | Hemisphere | N | Observations | ICC | CI lower | CI upper |
|---|---|---|---|---|---|---|---|
| LC | 1.5T | L | 267 | 912 | 0.478 | 0.409 | 0.547 |
| LC | 1.5T | R | 267 | 912 | 0.533 | 0.466 | 0.596 |
| SN-VTA | 1.5T | L | 268 | 913 | 0.653 | 0.595 | 0.703 |
| SN-VTA | 1.5T | R | 267 | 911 | 0.595 | 0.532 | 0.652 |
| LC | 3T | L | 63 | 188 | 0.729 | 0.615 | 0.812 |
| LC | 3T | R | 63 | 188 | 0.618 | 0.477 | 0.728 |
| SN-VTA | 3T | L | 62 | 186 | 0.797 | 0.704 | 0.860 |
| SN-VTA | 3T | R | 62 | 186 | 0.810 | 0.721 | 0.869 |

ICC, intraclass correlation coefficient. CI, confidence interval.

**Supplementary Table 3. Associations of LC and SN-VTA CR with neuropathological hypopigmentation**

| Region | Rating | N | Rating absent N | Rating present N | β | SE | CI lower | CI upper | P |
|---|---|---|---|---|---|---|---|---|---|
| LC | LC hypopigmentation | 50 | 27 | 23 | −0.036 | 0.015 | −0.066 | −0.006 | 0.019 |
| LC | SN hypopigmentation | 54 | 37 | 17 | −0.021 | 0.012 | −0.045 | 0.003 | 0.085 |
| SN-VTA | LC hypopigmentation | 50 | 27 | 23 | −0.010 | 0.007 | −0.024 | 0.003 | 0.127 |
| SN-VTA | SN hypopigmentation | 54 | 37 | 17 | −0.008 | 0.007 | −0.022 | 0.005 | 0.219 |

SE, standard error. P, P value. CI, confidence interval.

CR, contrast ratio.

**Supplementary Table 4. Pairwise associations among neuropathological measures**

| Region | Measure 1 | Measure 2 | Pairwise N | r | P |
|---|---|---|---|---|---|
| LC | Pigmentation preserved | Amyloid absent | 25 | 0.351 | 0.116 |
| LC | Pigmentation preserved | Tau absent | 24 | 0.060 | 1.000 |
| LC | Pigmentation preserved | Alpha-synuclein / Lewy bodies absent | 22 | 0.226 | 0.376 |
| LC | Pigmentation preserved | In neuron density | 25 | 0.531 | 0.006 |
| LC | Amyloid absent | Tau absent | 29 | 0.153 | 0.651 |
| LC | Amyloid absent | Alpha-synuclein / Lewy bodies absent | 26 | −0.087 | 1.000 |
| LC | Amyloid absent | In neuron density | 30 | 0.227 | 0.227 |
| LC | Tau absent | Alpha-synuclein / Lewy bodies absent | 25 | −0.421 | 0.070 |
| LC | Tau absent | In neuron density | 29 | −0.302 | 0.111 |
| LC | Alpha-synuclein / Lewy bodies absent | In neuron density | 26 | 0.208 | 0.307 |
| SN | Pigmentation preserved | Amyloid absent | 28 | 0.067 | 1.000 |
| SN | Pigmentation preserved | Tau absent | 27 | 0.271 | 0.274 |

| Region | Measure 1 | Measure 2 | Pairwise N | r | P |
|---|---|---|---|---|---|
| SN | Pigmentation preserved | Alpha-synuclein / Lewy bodies absent | 25 | 0.542 | 0.011 |
| SN | Pigmentation preserved | In neuron density | 28 | −0.094 | 0.635 |
| SN | Amyloid absent | Tau absent | 31 | 0.243 | 0.535 |
| SN | Amyloid absent | Alpha-synuclein / Lewy bodies absent | 29 | 0.296 | 0.143 |
| SN | Amyloid absent | In neuron density | 32 | −0.155 | 0.396 |
| SN | Tau absent | Alpha-synuclein / Lewy bodies absent | 28 | 0.067 | 1.000 |
| SN | Tau absent | In neuron density | 31 | 0.098 | 0.600 |
| SN | Alpha-synuclein / Lewy bodies absent | In neuron density | 29 | 0.159 | 0.410 |

P, P value.

**Supplementary Table 5. PLSC associations of LC and SN-VTA CR with neuropathology**

| Region | N | P |
|---|---|---|
| LC | 20 | 0.452 |
| SN-VTA | 23 | 0.302 |

PLSC, partial least squares correlation. CR, contrast ratio. P, P value.

**Supplementary Table 6. PLSC loadings for neuropathological measures**

| Region | Neuropathological measure | Loading |
|---|---|---|
| LC | Pigmentation preserved | 0.624 |
| LC | Amyloid absent | −0.512 |
| LC | Tau absent | −0.238 |
| LC | Alpha-synuclein / Lewy bodies absent | 0.478 |
| LC | ln neuron density | 0.419 |
| SN-VTA | Pigmentation preserved | 0.880 |
| SN-VTA | Amyloid absent | 0.408 |
| SN-VTA | Tau absent | 0.432 |
| SN-VTA | Alpha-synuclein / Lewy bodies absent | 0.844 |
| SN-VTA | ln neuron density | 0.030 |

PLSC, partial least squares correlation.

**Supplementary Table 7. Demographic associations of LC and SN-VTA CR**

| Region | Variables | β | SE | t | P |
|---|---|---|---|---|---|
| LC | Intercept | 0.134 | 0.007 | 18.541 | $6.222 \times 10^{-44}$ |
| LC | Baseline age (years) | −0.001 | $2.035 \times 10^{-4}$ | −4.948 | $9.501 \times 10^{-7}$ |
| LC | Imaging time (years) | $-8.451 \times 10^{-4}$ | $4.243 \times 10^{-4}$ | −1.992 | 0.047 |
| LC | Male sex (reference level: female) | −0.003 | 0.003 | −0.994 | 0.320 |
| LC | Baseline MCI (reference level: CN) | −0.003 | 0.003 | −0.793 | 0.428 |
| LC | Baseline AD (reference level: CN) | −0.002 | 0.004 | −0.536 | 0.592 |
| LC | Right versus left hemisphere | 0.002 | $6.740 \times 10^{-4}$ | 3.280 | 0.001 |
| LC | 3 T (reference level: 1.5 T) | 0.078 | 0.003 | 22.527 | $1.340 \times 10^{-102}$ |
| LC | Education (years) | $1.204 \times 10^{-4}$ | $4.764 \times 10^{-4}$ | 0.253 | 0.801 |
| LC | APOE ε4 allele count | 0.001 | 0.002 | 0.569 | 0.569 |
| LC | Entropy focus criterion | −0.170 | 0.047 | −3.619 | $3.011 \times 10^{-4}$ |
| LC | Echo time 0.011 s versus reference level | 0.012 | 0.007 | 1.806 | 0.073 |

| Region | Variables | β | SE | t | P |
|---|---|---|---|---|---|
| LC | Echo time 0.012 s versus reference level | 0.039 | 0.007 | 5.682 | $6.032 \times 10^{-8}$ |
| LC | Echo time 0.013 s versus reference level | 0.025 | 0.008 | 3.248 | 0.001 |
| SN-VTA | Intercept | 0.162 | 0.004 | 37.423 | $5.033 \times 10^{-97}$ |
| SN-VTA | Baseline age (years) | $-6.556 \times 10^{-4}$ | $1.321 \times 10^{-4}$ | −4.962 | $8.831 \times 10^{-7}$ |
| SN-VTA | Imaging time (years) | $-5.373 \times 10^{-4}$ | $2.363 \times 10^{-4}$ | −2.274 | 0.024 |
| SN-VTA | Male sex (reference: female) | $-1.758 \times 10^{-4}$ | 0.002 | −0.088 | 0.930 |
| SN-VTA | Baseline MCI (reference level: CN) | −0.003 | 0.002 | −1.426 | 0.154 |
| SN-VTA | Baseline AD (reference level: CN) | −0.005 | 0.003 | −2.006 | 0.045 |
| SN-VTA | Right versus left hemisphere | −0.006 | $3.350 \times 10^{-4}$ | −18.236 | $6.374 \times 10^{-71}$ |
| SN-VTA | 3 T (reference level: 1.5 T) | 0.032 | 0.002 | 16.871 | $7.464 \times 10^{-61}$ |
| SN-VTA | Education (years) | $6.470 \times 10^{-5}$ | $3.084 \times 10^{-4}$ | 0.210 | 0.834 |
| SN-VTA | APOE ε4 allele count | 0.001 | 0.001 | 0.965 | 0.335 |

| Region | Variables | β | SE | t | P |
|---|---|---|---|---|---|
| SN-VTA | Entropy focus criterion | −0.107 | 0.026 | −4.075 | $4.723 \times 10^{-5}$ |
| SN-VTA | Echo time 0.011 s versus reference level | 0.006 | 0.004 | 1.514 | 0.132 |
| SN-VTA | Echo time 0.012 s versus reference level | 0.022 | 0.004 | 5.394 | $2.031 \times 10^{-7}$ |
| SN-VTA | Echo time 0.013 s versus reference level | 0.018 | 0.004 | 3.935 | $1.173 \times 10^{-4}$ |

SE, standard error. P, P value. CR, contrast ratio. EFC, entropy focus criterion. APOE, apolipoprotein E. CN, cognitively normal. MCI, mild cognitive impairment. AD, Alzheimer’s disease.

**Supplementary Table 8. Hemispheric differences in LC and SN-VTA CRs by manufacturer.**

| Nucleus | Manufacturer | Field strength | N / scans | β (95% CI) | SE | P |
|---|---|---|---|---|---|---|
| LC | GE | Pooled | 320 / 1,610 | $-5.28 \times 10^{-5}$ (−0.002, 0.001) | $7.83 \times 10^{-4}$ | 0.946 |
| LC | Siemens | Pooled | 413 / 1,766 | 0.004 (0.002, 0.006) | 0.001 | $1.40 \times 10^{-4}$ |
| LC | Philips | Pooled | 11 / 20 | 0.016 (0.003, 0.029) | 0.006 | 0.020 |
| SN-VTA | GE | Pooled | 320 / 1,608 | −0.004 (−0.005, −0.003) | $4.20 \times 10^{-4}$ | $3.75 \times 10^{-18}$ |
| SN-VTA | Siemens | Pooled | 413 / 1,760 | −0.008 (−0.009, −0.007) | $5.11 \times 10^{-4}$ | $8.09 \times 10^{-55}$ |
| SN-VTA | Philips | Pooled | 11 / 20 | −0.011 (−0.019, −0.004) | 0.004 | 0.003 |
| LC | GE | 1.5 T | 320 / 1,547 | $-1.67 \times 10^{-4}$ (−0.002, 0.001) | $7.86 \times 10^{-4}$ | 0.832 |
| LC | GE | 3 T | 23 / 63 | 0.003 (−0.008, 0.013) | 0.005 | 0.604 |
| LC | Siemens | 1.5 T | 353 / 1,394 | 0.008 (0.005, 0.010) | 0.001 | $2.53 \times 10^{-12}$ |
| LC | Siemens | 3 T | 106 / 372 | −0.009 (−0.015, −0.003) | 0.003 | 0.004 |
| LC | Philips | 1.5 T | 0 / 0 | — | — | — |
| LC | Philips | 3 T | 11 / 20 | 0.016 (0.003, 0.029) | 0.006 | 0.020 |
| SN-VTA | GE | 1.5 T | 320 / 1,545 | −0.004 (−0.004, −0.003) | $4.31 \times 10^{-4}$ | $1.01 \times 10^{-16}$ |
| SN-VTA | GE | 3 T | 23 / 63 | −0.006 (−0.009, −0.002) | 0.002 | 0.004 |
| SN-VTA | Siemens | 1.5 T | 353 / 1,390 | −0.008 (−0.009, −0.007) | $5.44 \times 10^{-4}$ | $7.99 \times 10^{-43}$ |
| SN-VTA | Siemens | 3 T | 106 / 370 | −0.010 (−0.013, −0.008) | 0.001 | $6.83 \times 10^{-14}$ |
| SN-VTA | Philips | 1.5 T | 0 / 0 | — | — | — |
| SN-VTA | Philips | 3 T | 11 / 20 | −0.011 (−0.019, −0.004) | 0.004 | 0.003 |

**Supplementary Table 9. Concurrent associations of LC and SN-VTA CR with cognitive, clinical and AD-related measures.**

Results for all 38 outcomes in the primary and concurrent sensitivity models, including β, SE, 95% confidence intervals and P values. Supplied as a separate CSV file.

File: electronic_tables/Supplementary_Table_9_Concurrent_Associations.csv

**Supplementary Table 10. Agreement of concurrent and prospective CR coefficients across model specifications**

| Model comparison | Region | Same direction (n/N) | r | Spearman ρ |
|---|---|---|---|---|
| Concurrent: CR response → outcome response | LC | 31/38 | — | — |
| Concurrent: CR response → outcome response | SN-VTA | 29/38 | — | — |
| Concurrent: separate LC and SN-VTA models → both included | LC | 36/38 | 0.992 | 0.989 |
| Concurrent: separate LC and SN-VTA models → both included | SN-VTA | 37/38 | 0.996 | 0.995 |
| Prospective: primary → ≥180 days | LC | 36/38 | 0.995 | 0.997 |
| Prospective: primary → ≥180 days | SN-VTA | 38/38 | 0.997 | 0.995 |
| Prospective: primary → ≥90 days | LC | 37/38 | 0.994 | 0.993 |
| Prospective: primary → ≥90 days | SN-VTA | 38/38 | 0.995 | 0.991 |
| Prospective: primary → LC and SN-VTA included | LC | 36/38 | 0.990 | 0.986 |
| Prospective: primary → LC and SN-VTA included | SN-VTA | 36/38 | 0.988 | 0.982 |
| Prospective: primary → CN/MCI at MRI | LC | 31/37 | 0.840 | 0.821 |
| Prospective: primary → CN/MCI at MRI | SN-VTA | 33/37 | 0.813 | 0.769 |
| Prospective: separate LC and SN-VTA models → both included | LC | 36/38 | 0.991 | 0.985 |
| Prospective: separate LC and SN-VTA models → both included | SN-VTA | 36/38 | 0.989 | 0.985 |
| Outcome-response concurrent → primary prospective | LC | 24/38 | 0.427 | 0.447 |
| Outcome-response concurrent → primary prospective | SN-VTA | 32/38 | 0.635 | 0.644 |

**Supplementary Table 11. Prospective associations of LC and SN-VTA CR with cognitive, clinical and AD-related measures.**

File: Supplementary_Table_11_Prospective_Associations.csv

**Supplementary Table 12. Lag-dependent association intervals for LC and SN-VTA CR.**

File: Supplementary_Table_12_Lag_Interval_Summary.csv

**Supplementary Table 13. Summary of lag-dependent association intervals**

| Model type | Region | Outcomes with an interval (n/38) | Outcomes with a positive interval (n) | Outcomes with a negative interval (n) | Memory interval (years) |
|---|---|---|---|---|---|
| Primary (full range) | LC | 26/38 | 18 | 15 | 7.846-18.801 |
| Primary (full range) | SN-VTA | 26/38 | 15 | 12 | 8.963-18.801 |
| ≥180 days | LC | 26/38 | 16 | 14 | 7.766-18.801 |
| ≥180 days | SN-VTA | 25/38 | 15 | 11 | 8.709-18.801 |
| CN/MCI at MRI | LC | 26/38 | 15 | 13 | 7.606-18.801 |
| CN/MCI at MRI | SN-VTA | 21/38 | 14 | 11 | 9.736-18.801 |
| LC and SN-VTA included | LC | 25/38 | 18 | 12 | 8.034-18.801 |
| LC and SN-VTA included | SN-VTA | 24/38 | 13 | 13 | 9.130-18.801 |
| Primary (5th-95th percentile range) | LC | 24/38 | 18 | 12 | 7.724-10.179 |
| Primary (5th-95th percentile range) | SN-VTA | 20/38 | 13 | 10 | 8.823-10.191 |

**Supplementary Table 14**. Complete regional LV1 saliences and bootstrap ratios.
File: Supplementary_Table_14_Regional_PLSC.csv

**Supplementary Table 15. PLSC analyses**

| Model type | Parameter | N | r | P |
|---|---|---|---|---|
| LC | Intercept | 697 | 0.228 | <0.001 |
| LC | Slope | 538 | 0.178 | 0.007 |
| SN-VTA | Intercept | 698 | 0.266 | <0.001 |
| SN-VTA | Slope | 539 | 0.178 | 0.006 |
| LC unique | Intercept | 697 | 0.219 | $7.998 \times 10^{-4}$ |
| LC unique | Slope | 538 | 0.173 | 0.019 |
| SN-VTA unique | Intercept | 697 | 0.250 | <0.001 |
| SN-VTA unique | Slope | 538 | 0.173 | 0.013 |

PLSC, partial least squares correlation. P, P value.

**Supplementary Table 16. Unique structural PLSC and primary molecular-correspondence results**

| Model type | Region | Parameter / PET tracer | r | Spearman ρ | P |
|---|---|---|---|---|---|
| Unique PLSC | LC | Intercept | 0.219 | — | $7.998 \times 10^{-4}$ |
| Unique PLSC | LC | Slope | 0.173 | — | 0.019 |
| Unique PLSC | SN-VTA | Intercept | 0.250 | — | 0 |
| Unique PLSC | SN-VTA | Slope | 0.173 | — | 0.013 |
| Unique-pattern PET | LC | MRB | — | 0.163 | 0.293 |
| Unique-pattern PET | LC | methylreboxetine | — | 0.361 | 0.008 |
| Unique-pattern PET | LC | SCH23390 | — | 0.138 | 0.196 |
| Unique-pattern PET | LC | FE-PE2I | — | 0.029 | 0.814 |
| Unique-pattern PET | LC | FLB457 | — | 0.225 | 0.264 |
| Unique-pattern PET | LC | fallypride | — | 0.256 | 0.219 |
| Unique-pattern PET | SN-VTA | MRB | — | −0.088 | 0.622 |
| Unique-pattern PET | SN-VTA | methylreboxetine | — | 0.106 | 0.622 |
| Unique-pattern PET | SN-VTA | SCH23390 | — | 0.472 | $4.000 \times 10^{-4}$ |
| Unique-pattern PET | SN-VTA | FE-PE2I | — | −0.030 | 0.807 |
| Unique-pattern PET | SN-VTA | FLB457 | — | 0.584 | 0.002 |
| Unique-pattern PET | SN-VTA | fallypride | — | 0.543 | 0.003 |

**Supplementary Table 17. Molecular PET correspondence**

| Model type | PET target | PET tracer | PET source | Family | Spearman ρ | P | $P_{FDR}$ | Adjustment | Partial ρ | $P_{partial}$ | $P_{partial,FDR}$ |
|---|---|---|---|---|---|---|---|---|---|---|---|
| LC unique | Serotonin 1A receptor | WAY100635 | savli2012 | exploratory | 0.282 | 0.119 | 0.250 | — | — | — | — |
| LC unique | Serotonin 1B receptor | P943 | gallezot2010 | exploratory | −0.111 | 0.322 | 0.469 | — | — | — | — |
| LC unique | Serotonin 2A receptor | altanserin | savli2012 | exploratory | −0.036 | 0.811 | 0.865 | — | — | — | — |
| LC unique | Serotonin 4 receptor | SB207145 | beliveau2017 | exploratory | 0.261 | 0.080 | 0.250 | — | — | — | — |
| LC unique | Serotonin 6 receptor | GSK215083 | radnakrishnan2018 | exploratory | 0.112 | 0.411 | 0.541 | — | — | — | — |
| LC unique | Histamine H3 receptor | GSK189254 | gallezot2017 | exploratory | 0.115 | 0.440 | 0.541 | — | — | — | — |
| LC unique | Nicotinic acetylcholine alpha-4 beta-2 receptor | flubatine | hillmer2016 | exploratory | 0.069 | 0.683 | 0.780 | — | — | — | — |
| LC unique | Muscarinic M1 receptor | LSN3172176 | naganawa2020 | exploratory | 0.022 | 0.865 | 0.865 | — | — | — | — |
| LC unique | Vesicular acetylcholine transporter | FEOBV | Hansen et al. (2022) public subset | exploratory | 0.275 | 0.095 | 0.250 | — | — | — | — |
| LC unique | Cannabinoid receptor 1 | OMAR | normandin2015 | exploratory | 0.226 | 0.105 | 0.250 | — | — | — | — |

| Model type | PET target | PET tracer | PET source | Family | Spearman ρ | P | $P_{FDR}$ | Adjustment | Partial ρ | $P_{partial}$ | $P_{partial,FDR}$ |
|---|---|---|---|---|---|---|---|---|---|---|---|
| LC unique | GABA-A alpha-5-enriched benzodiazepine-site availability | RO15-4513 | lukow2022 | exploratory | 0.214 | 0.124 | 0.250 | — | — | — | — |
| LC unique | Broad GABA-A benzodiazepine-site availability | flumazenil | norgaard2021 | exploratory | −0.316 | 0.021 | 0.250 | — | — | — | — |
| LC unique | Kappa-opioid receptor | LY2795050 | vijay2018 | exploratory | 0.255 | 0.217 | 0.386 | — | — | — | — |
| LC unique | Metabotropic glutamate receptor 5 | ABP688 | smart2019 | exploratory | 0.154 | 0.288 | 0.461 | — | — | — | — |
| LC unique | NMDA receptor open-channel-sensitive availability | GE-179 | galovic2021 | exploratory | 0.159 | 0.125 | 0.250 | — | — | — | — |
| LC unique | Mu-opioid receptor | carfentanil | kantonen2020 | exploratory | 0.244 | 0.062 | 0.250 | — | — | — | — |
| LC unique | Norepinephrine transporter | MRB | ding2010 | primary | 0.163 | 0.293 | 0.351 | — | — | — | — |
| LC unique | Norepinephrine transporter | MRB | hesse2017 | primary | 0.361 | 0.008 | 0.050 | — | — | — | — |
| LC unique | Dopamine D1-like receptor | SCH23390 | kaller2017 | primary | 0.138 | 0.196 | 0.351 | — | — | — | — |
| LC unique | Dopamine transporter | FE-PE2I | sasaki2012 | primary | 0.029 | 0.814 | 0.814 | — | — | — | — |
| LC unique | Dopamine D2-family receptor | FLB457 | smith2017 | primary | 0.225 | 0.264 | 0.351 | — | — | — | — |
| LC unique | Dopamine D2-family receptor | fallypride | jaworska2020 | primary | 0.256 | 0.219 | 0.351 | — | — | — | — |
| LC | Serotonin 1A receptor | WAY100635 | savli2012 | exploratory | 0.426 | 0.048 | 0.099 | MRB (Hesse) | 0.390 | 0.022 | 0.118 |

| Model type | PET target | PET tracer | PET source | Family | Spearman ρ | P | $P_{FDR}$ | Adjustment | Partial ρ | $P_{partial}$ | $P_{partial,FDR}$ |
|---|---|---|---|---|---|---|---|---|---|---|---|
| LC | Serotonin 1B receptor | P943 | gallezot2010 | exploratory | −0.070 | 0.591 | 0.631 | MRB (Hesse) | −0.179 | 0.175 | 0.701 |
| LC | Serotonin 2A receptor | altanserin | savli2012 | exploratory | 0.073 | 0.663 | 0.663 | MRB (Hesse) | −0.050 | 0.724 | 0.990 |
| LC | Serotonin 4 receptor | SB207145 | beliveau2017 | exploratory | 0.343 | 0.031 | 0.099 | MRB (Hesse) | 0.394 | 0.006 | 0.102 |
| LC | Serotonin 6 receptor | GSK215083 | radnakrishnan2018 | exploratory | 0.204 | 0.213 | 0.310 | MRB (Hesse) | −0.024 | 0.878 | 0.990 |
| LC | Histamine H3 receptor | GSK189254 | gallezot2017 | exploratory | 0.194 | 0.242 | 0.322 | MRB (Hesse) | 0.076 | 0.963 | 0.990 |
| LC | Nicotinic acetylcholine alpha-4 beta-2 receptor | flubatine | hillmer2016 | exploratory | 0.105 | 0.575 | 0.631 | MRB (Hesse) | −0.022 | 0.915 | 0.990 |
| LC | Muscarinic M1 receptor | LSN3172176 | naganawa2020 | exploratory | 0.130 | 0.353 | 0.435 | MRB (Hesse) | 0.042 | 0.752 | 0.990 |
| LC | Vesicular acetylcholine transporter | FEOBV | Hansen et al. (2022) public subset | exploratory | 0.342 | 0.049 | 0.099 | MRB (Hesse) | 0.164 | 0.784 | 0.990 |
| LC | Cannabinoid receptor 1 | OMAR | normandin2015 | exploratory | 0.334 | 0.041 | 0.099 | MRB (Hesse) | 0.361 | 0.239 | 0.762 |
| LC | GABA-A alpha-5-enriched benzodiazepine-site availability | RO15-4513 | lukow2022 | exploratory | 0.363 | 0.024 | 0.099 | MRB (Hesse) | 0.294 | 0.312 | 0.762 |
| LC | Broad GABA-A benzodiazepine-site availability | flumazenil | norgaard2021 | exploratory | −0.294 | 0.042 | 0.099 | MRB (Hesse) | −0.229 | 0.013 | 0.102 |
| LC | Kappa-opioid receptor | LY2795050 | vijay2018 | exploratory | 0.301 | 0.213 | 0.310 | MRB (Hesse) | 0.142 | 0.910 | 0.990 |
| LC | Metabotropic glutamate receptor 5 | ABP688 | smart2019 | exploratory | 0.237 | 0.149 | 0.265 | MRB (Hesse) | 0.024 | 0.953 | 0.990 |

| Model type | PET target | PET tracer | PET source | Family | Spearman ρ | P | $P_{FDR}$ | Adjustment | Partial ρ | $P_{partial}$ | $P_{partial,FDR}$ |
|---|---|---|---|---|---|---|---|---|---|---|---|
| LC | NMDA receptor open-channel-sensitive availability | GE-179 | galovic2021 | exploratory | 0.253 | 0.023 | 0.099 | MRB (Hesse) | −0.001 | 0.990 | 0.990 |
| LC | Mu-opioid receptor | carfentanil | kantonen2020 | exploratory | 0.353 | 0.013 | 0.099 | MRB (Hesse) | 0.334 | 0.333 | 0.762 |
| LC | Norepinephrine transporter | MRB | ding2010 | primary | 0.153 | 0.341 | 0.409 | — | — | — | — |
| LC | Norepinephrine transporter | MRB | hesse2017 | primary | 0.403 | 0.005 | 0.028 | — | — | — | — |
| LC | Dopamine D1-like receptor | SCH23390 | kaller2017 | primary | 0.271 | 0.024 | 0.071 | — | — | — | — |
| LC | Dopamine transporter | FE-PE2I | sasaki2012 | primary | 0.011 | 0.931 | 0.931 | — | — | — | — |
| LC | Dopamine D2-family receptor | FLB457 | smith2017 | primary | 0.351 | 0.113 | 0.170 | — | — | — | — |
| LC | Dopamine D2-family receptor | fallypride | jaworska2020 | primary | 0.379 | 0.108 | 0.170 | — | — | — | — |
| SN-VTA unique | Serotonin 1A receptor | WAY100635 | savli2012 | exploratory | 0.628 | 0.001 | 0.010 | — | — | — | — |
| SN-VTA unique | Serotonin 1B receptor | P943 | gallezot2010 | exploratory | −0.003 | 0.977 | 0.977 | — | — | — | — |
| SN-VTA unique | Serotonin 2A receptor | altanserin | savli2012 | exploratory | 0.316 | 0.017 | 0.040 | — | — | — | — |
| SN-VTA unique | Serotonin 4 receptor | SB207145 | beliveau2017 | exploratory | 0.413 | 0.009 | 0.024 | — | — | — | — |
| SN-VTA unique | Serotonin 6 receptor | GSK215083 | radnakrishnan2018 | exploratory | 0.295 | 0.061 | 0.099 | — | — | — | — |
| SN-VTA unique | Histamine H3 receptor | GSK189254 | gallezot2017 | exploratory | 0.303 | 0.082 | 0.120 | — | — | — | — |

| Model type | PET target | PET tracer | PET source | Family | Spearman ρ | P | $P_{FDR}$ | Adjustment | Partial ρ | $P_{partial}$ | $P_{partial,FDR}$ |
|---|---|---|---|---|---|---|---|---|---|---|---|
| SN-VTA unique | Nicotinic acetylcholine alpha-4 beta-2 receptor | flubatine | hillmer2016 | exploratory | 0.161 | 0.403 | 0.496 | — | — | — | — |
| SN-VTA unique | Muscarinic M1 receptor | LSN3172176 | naganawa2020 | exploratory | 0.325 | 0.006 | 0.023 | — | — | — | — |
| SN-VTA unique | Vesicular acetylcholine transporter | FEOBV | Hansen et al. (2022) public subset | exploratory | 0.263 | 0.204 | 0.272 | — | — | — | — |
| SN-VTA unique | Cannabinoid receptor 1 | OMAR | normandin2015 | exploratory | 0.504 | 0.002 | 0.011 | — | — | — | — |
| SN-VTA unique | GABA-A alpha-5-enriched benzodiazepine-site availability | RO15-4513 | lukow2022 | exploratory | 0.579 | $5.000 \times 10^{-4}$ | 0.008 | — | — | — | — |
| SN-VTA unique | Broad GABA-A benzodiazepine-site availability | flumazenil | norgaard2021 | exploratory | 0.031 | 0.830 | 0.886 | — | — | — | — |
| SN-VTA unique | Kappa-opioid receptor | LY2795050 | vijay2018 | exploratory | 0.242 | 0.525 | 0.599 | — | — | — | — |
| SN-VTA unique | Metabotropic glutamate receptor 5 | ABP688 | smart2019 | exploratory | 0.288 | 0.062 | 0.099 | — | — | — | — |
| SN-VTA unique | NMDA receptor open-channel-sensitive availability | GE-179 | galovic2021 | exploratory | 0.278 | 0.028 | 0.055 | — | — | — | — |
| SN-VTA unique | Mu-opioid receptor | carfentanil | kantonen2020 | exploratory | 0.474 | 0.007 | 0.023 | — | — | — | — |
| SN-VTA unique | Norepinephrine transporter | MRB | ding2010 | primary | −0.088 | 0.622 | 0.747 | — | — | — | — |
| SN-VTA unique | Norepinephrine transporter | MRB | hesse2017 | primary | 0.106 | 0.622 | 0.747 | — | — | — | — |

| Model type | PET target | PET tracer | PET source | Family | Spearman ρ | P | $P_{FDR}$ | Adjustment | Partial ρ | $P_{partial}$ | $P_{partial,FDR}$ |
|---|---|---|---|---|---|---|---|---|---|---|---|
| SN-VTA unique | Dopamine D1-like receptor | SCH23390 | kaller2017 | primary | 0.472 | $4.000 \times 10^{-4}$ | 0.002 | — | — | — | — |
| SN-VTA unique | Dopamine transporter | FE-PE2I | sasaki2012 | primary | −0.030 | 0.807 | 0.807 | — | — | — | — |
| SN-VTA unique | Dopamine D2-family receptor | FLB457 | smith2017 | primary | 0.584 | 0.002 | 0.006 | — | — | — | — |
| SN-VTA unique | Dopamine D2-family receptor | fallypride | jaworska2020 | primary | 0.543 | 0.003 | 0.006 | — | — | — | — |
| SN-VTA | Serotonin 1A receptor | WAY100635 | savli2012 | explorator y | 0.661 | $5.000 \times 10^{-4}$ | 0.004 | Joint | 0.301 | 0.029 | 0.446 |
| | | | | | | | | SCH23390 | 0.544 | 0.002 | 0.014 |
| | | | | | | | | FLB457 | 0.360 | 0.006 | 0.052 |
| | | | | | | | | fallypride | 0.427 | $1.00 \times 10^{-3}$ | 0.008 |
| SN-VTA | Serotonin 1B receptor | P943 | gallezot2010 | explorator y | 0.004 | 0.975 | 0.975 | Joint | −0.044 | 0.726 | 0.817 |
| | | | | | | | | SCH23390 | −0.180 | 0.248 | 0.617 |
| | | | | | | | | FLB457 | 0.069 | 0.442 | 0.544 |
| | | | | | | | | fallypride | 0.030 | 0.867 | 0.867 |
| SN-VTA | Serotonin 2A receptor | altanserin | savli2012 | explorator y | 0.327 | 0.013 | 0.027 | Joint | 0.188 | 0.168 | 0.698 |
| | | | | | | | | SCH23390 | 0.158 | 0.432 | 0.768 |
| | | | | | | | | FLB457 | 0.261 | 0.061 | 0.222 |
| | | | | | | | | fallypride | 0.217 | 0.110 | 0.295 |
| SN-VTA | Serotonin 4 receptor | SB207145 | beliveau2017 | explorator y | 0.453 | 0.002 | 0.008 | Joint | 0.127 | 0.174 | 0.698 |
| | | | | | | | | SCH23390 | 0.400 | 0.002 | 0.014 |
| | | | | | | | | FLB457 | 0.115 | 0.168 | 0.245 |
| | | | | | | | | fallypride | 0.084 | 0.355 | 0.516 |
| SN-VTA | Serotonin 6 receptor | GSK215083 | radnakrishnan2 018 | explorator y | 0.314 | 0.060 | 0.095 | Joint | 0.073 | 0.639 | 0.817 |
| | | | | | | | | SCH23390 | 0.065 | 0.734 | 0.909 |
| | | | | | | | | FLB457 | 0.191 | 0.112 | 0.222 |
| | | | | | | | | fallypride | 0.177 | 0.193 | 0.386 |

| Model type | PET target | PET tracer | PET source | Family | Spearman ρ | P | $P_{FDR}$ | Adjustment | Partial ρ | $P_{partial}$ | $P_{partial,FDR}$ |
|---|---|---|---|---|---|---|---|---|---|---|---|
| SN-VTA | Histamine H3 receptor | GSK189254 | gallezot2017 | exploratory | 0.302 | 0.091 | 0.132 | Joint | −0.085 | 0.429 | 0.817 |
| | | | | | | | | SCH23390 | −0.044 | 0.881 | 0.939 |
| | | | | | | | | FLB457 | 0.067 | 0.434 | 0.544 |
| | | | | | | | | fallypride | 0.147 | 0.152 | 0.346 |
| SN-VTA | Nicotinic acetylcholine alpha-4 beta-2 receptor | flubatine | hillmer2016 | exploratory | 0.164 | 0.443 | 0.545 | Joint | 0.094 | 0.408 | 0.817 |
| | | | | | | | | SCH23390 | 0.006 | 0.970 | 0.970 |
| | | | | | | | | FLB457 | 0.133 | 0.107 | 0.222 |
| | | | | | | | | fallypride | 0.226 | 0.039 | 0.124 |
| SN-VTA | Muscarinic M1 receptor | LSN3172176 | naganawa2020 | exploratory | 0.332 | 0.003 | 0.008 | Joint | 0.055 | 0.624 | 0.817 |
| | | | | | | | | SCH23390 | 0.114 | 0.308 | 0.617 |
| | | | | | | | | FLB457 | 0.175 | 0.126 | 0.222 |
| | | | | | | | | fallypride | 0.117 | 0.332 | 0.516 |
| SN-VTA | Vesicular acetylcholine transporter | FEOBV | Hansen et al. (2022) public subset | exploratory | 0.299 | 0.130 | 0.174 | Joint | −0.143 | 0.279 | 0.745 |
| | | | | | | | | SCH23390 | −0.078 | 0.666 | 0.909 |
| | | | | | | | | FLB457 | 0.072 | 0.489 | 0.558 |
| | | | | | | | | fallypride | 0.042 | 0.725 | 0.827 |
| SN-VTA | Cannabinoid receptor 1 | OMAR | normandin2015 | exploratory | 0.514 | 0.003 | 0.008 | Joint | 0.113 | 0.279 | 0.745 |
| | | | | | | | | SCH23390 | 0.375 | 0.005 | 0.025 |
| | | | | | | | | FLB457 | 0.127 | 0.139 | 0.222 |
| | | | | | | | | fallypride | 0.255 | 0.008 | 0.033 |
| SN-VTA | GABA-A alpha-5-enriched benzodiazepine-site availability | RO15-4513 | lukow2022 | exploratory | 0.611 | $3.000 \times 10^{-4}$ | 0.004 | Joint | 0.189 | 0.056 | 0.446 |
| | | | | | | | | SCH23390 | 0.426 | 0.038 | 0.152 |
| | | | | | | | | FLB457 | 0.322 | $6.00 \times 10^{-4}$ | 0.010 |
| | | | | | | | | fallypride | 0.362 | $2.00 \times 10^{-4}$ | 0.003 |
| SN-VTA | Broad GABA-A benzodiazepine-site availability | flumazenil | norgaard2021 | exploratory | −0.020 | 0.891 | 0.951 | Joint | 0.043 | 0.589 | 0.817 |
| | | | | | | | | SCH23390 | −0.063 | 0.738 | 0.909 |
| | | | | | | | | FLB457 | 0.067 | 0.549 | 0.586 |

| Model type | PET target | PET tracer | PET source | Family | Spearman ρ | P | $P_{FDR}$ | Adjustment | Partial ρ | $P_{partial}$ | $P_{partial,FDR}$ |
|---|---|---|---|---|---|---|---|---|---|---|---|
| | | | | | | | | fallypride | 0.089 | 0.423 | 0.564 |
| SN-VTA | Kappa-opioid receptor | LY2795050 | vijay2018 | exploratory | 0.261 | 0.547 | 0.625 | Joint | −0.057 | 0.664 | 0.817 |
| | | | | | | | | SCH23390 | 0.092 | 0.655 | 0.909 |
| | | | | | | | | FLB457 | $7.49 \times 10^{-4}$ | 0.993 | 0.993 |
| | | | | | | | | fallypride | 0.044 | 0.729 | 0.827 |
| SN-VTA | Metabotropic glutamate receptor 5 | ABP688 | smart2019 | exploratory | 0.312 | 0.041 | 0.072 | Joint | −0.043 | 0.783 | 0.817 |
| | | | | | | | | SCH23390 | −0.027 | 0.868 | 0.939 |
| | | | | | | | | FLB457 | 0.168 | 0.131 | 0.222 |
| | | | | | | | | fallypride | 0.048 | 0.776 | 0.827 |
| SN-VTA | NMDA receptor open-channel-sensitive availability | GE-179 | galovic2021 | exploratory | 0.308 | 0.013 | 0.027 | Joint | −0.031 | 0.817 | 0.817 |
| | | | | | | | | SCH23390 | −0.183 | 0.289 | 0.617 |
| | | | | | | | | FLB457 | 0.209 | 0.064 | 0.222 |
| | | | | | | | | fallypride | 0.149 | 0.250 | 0.445 |
| SN-VTA | Mu-opioid receptor | carfentanil | kantonen2020 | exploratory | 0.505 | 0.002 | 0.008 | Joint | 0.031 | 0.752 | 0.817 |
| | | | | | | | | SCH23390 | 0.271 | 0.183 | 0.587 |
| | | | | | | | | FLB457 | 0.145 | 0.099 | 0.222 |
| | | | | | | | | fallypride | 0.246 | 0.008 | 0.033 |
| SN-VTA | Norepinephrine transporter | MRB | ding2010 | primary | −0.055 | 0.767 | 0.767 | — | — | — | — |
| SN-VTA | Norepinephrine transporter | MRB | hesse2017 | primary | 0.174 | 0.444 | 0.666 | — | — | — | — |
| SN-VTA | Dopamine D1-like receptor | SCH23390 | kaller2017 | primary | 0.491 | $2.000 \times 10^{-4}$ | 0.001 | — | — | — | — |
| SN-VTA | Dopamine transporter | FE-PE2I | sasaki2012 | primary | −0.051 | 0.682 | 0.767 | — | — | — | — |
| SN-VTA | Dopamine D2-family receptor | FLB457 | smith2017 | primary | 0.597 | 0.001 | 0.003 | — | — | — | — |

| Model type | PET target | PET tracer | PET source | Family | Spearman ρ | P | $P_{FDR}$ | Adjustment | Partial ρ | $P_{partial}$ | $P_{partial,FDR}$ |
|---|---|---|---|---|---|---|---|---|---|---|---|
| SN-VTA | Dopamine D2-family receptor | fallypride | jaworska2020 | primary | 0.572 | 0.001 | 0.003 | — | — | — | — |

PET, positron emission tomography; MRB, methylreboxetine; partial ρ, partial Spearman correlation. P and $P_{FDR}$ denote the spatial P values and their FDR-adjusted values. $\mathbf{P_{partial}}$ and $\mathbf{P_{partial,FDR}}$ denote exploratory spatial P values for partial correlations and their FDR-adjusted values, respectively. MRB (Hesse) denotes the Hesse et al. NET map; Joint denotes simultaneous adjustment for SCH23390, FLB457, and fallypride. “Unique” denotes other-nucleus-CR-adjusted PLSC patterns. Dashes indicate that no analysis was performed for that row.

**Supplementary Table 18. Clinical progression**

| Outcome | Model type | N | Events | Region | β | SE | OR | CI lower | CI upper | P | $P_{FDR}$ |
|---|---|---|---|---|---|---|---|---|---|---|---|
| MCI-to-AD dementia | Primary separate LC and SN-VTA models | 232 | 166 | LC | 0.607 | 0.207 | 1.835 | 1.222 | 2.754 | 0.003 | 0.007 |
| MCI-to-AD dementia | Primary separate LC and SN-VTA models | 232 | 166 | SN-VTA | 0.201 | 0.186 | 1.223 | 0.849 | 1.761 | 0.279 | 0.279 |
| Any progression | Primary separate LC and SN-VTA models | 335 | 180 | LC | 0.378 | 0.158 | 1.460 | 1.072 | 1.989 | 0.016 | 0.033 |
| Any progression | Primary separate LC and SN-VTA models | 335 | 180 | SN-VTA | 0.312 | 0.162 | 1.366 | 0.994 | 1.877 | 0.054 | 0.054 |
| MCI-to-AD dementia | Both LC and SN-VTA included | 232 | 166 | LC | 0.603 | 0.208 | 1.828 | 1.217 | 2.747 | 0.004 | — |

| Outcome | Model type | N | Events | Region | β | SE | OR | CI lower | CI upper | P | $P_{FDR}$ |
|---|---|---|---|---|---|---|---|---|---|---|---|
| MCI-to-AD dementia | Both LC and SN-VTA included | 232 | 166 | SN-VTA | 0.172 | 0.192 | 1.187 | 0.814 | 1.731 | 0.373 | — |
| MCI-to-AD dementia | Separate LC and SN-VTA models using the same observations | 232 | 166 | LC | 0.612 | 0.207 | 1.843 | 1.229 | 2.764 | 0.003 | — |
| MCI-to-AD dementia | Separate LC and SN-VTA models using the same observations | 232 | 166 | SN-VTA | 0.200 | 0.186 | 1.221 | 0.848 | 1.758 | 0.282 | — |
| Any progression | Both LC and SN-VTA included | 335 | 180 | LC | 0.377 | 0.158 | 1.457 | 1.068 | 1.987 | 0.017 | — |
| Any progression | Both LC and SN-VTA included | 335 | 180 | SN-VTA | 0.311 | 0.164 | 1.364 | 0.989 | 1.882 | 0.059 | — |

| Outcome | Model type | N | Events | Region | β | SE | OR | CI lower | CI upper | P | $P_{FDR}$ |
|---|---|---|---|---|---|---|---|---|---|---|---|
| Any progression | Separate LC and SN-VTA models using the same observations | 335 | 180 | LC | 0.384 | 0.157 | 1.468 | 1.079 | 1.998 | 0.015 | — |
| Any progression | Separate LC and SN-VTA models using the same observations | 335 | 180 | SN-VTA | 0.319 | 0.162 | 1.375 | 1.002 | 1.888 | 0.049 | — |

SE, standard error. P, P value. CI, confidence interval.

OR, odds ratio.

Logistic β contrasts and odds ratios are expressed per one SD lower CR. P denotes the raw two-sided P value. $P_{FDR}$ applies to the primary two-nucleus family within each endpoint; thus no adjusted value is reported for the comparison models.

**Supplementary Table 19. Clinical progression, mediation and continuous-amyloid memory sensitivities**

| Model type | Outcome / Interaction | Region | Effect | β / OR | P | $P_{FDR}$ |
|---|---|---|---|---|---|---|
| Progression: LC and SN-VTA included | MCI-to-AD dementia | LC | OR | 1.828 | 0.004 | — |
| Progression: LC and SN-VTA included | MCI-to-AD dementia | SN-VTA | OR | 1.187 | 0.373 | — |
| Progression: LC and SN-VTA included | Any progression | LC | OR | 1.457 | 0.017 | — |
| Progression: LC and SN-VTA included | Any progression | SN-VTA | OR | 1.364 | 0.059 | — |
| Later-T1 subset: original PLSC | MCI-to-AD dementia | LC | Indirect | −0.184 | 0.019 | 0.038 |
| Later-T1 subset: original PLSC | MCI-to-AD dementia | SN-VTA | Indirect | −0.096 | 0.262 | 0.262 |
| Later-T1 subset: original PLSC | Any progression | LC | Indirect | −0.106 | 0.025 | 0.049 |
| Later-T1 subset: original PLSC | Any progression | SN-VTA | Indirect | −0.086 | 0.122 | 0.122 |
| Later-T1 subset: re-estimated PLSC | MCI-to-AD dementia | LC | Indirect | −0.213 | 0.012 | 0.025 |
| Later-T1 subset: re-estimated PLSC | MCI-to-AD dementia | SN-VTA | Indirect | −0.137 | 0.146 | 0.146 |
| Later-T1 subset: re-estimated PLSC | Any progression | LC | Indirect | −0.137 | 0.013 | 0.025 |
| Later-T1 subset: re-estimated PLSC | Any progression | SN-VTA | Indirect | −0.109 | 0.081 | 0.081 |
| Amyloid-memory: continuous amyloid (Centiloid) | LC CR × amyloid × time | LC | Interaction | 0.011 | 0.168 | — |
| Amyloid-memory: continuous amyloid (Centiloid) | SN-VTA CR × amyloid × time | SN-VTA | Interaction | 0.005 | 0.565 | — |
| Amyloid-memory: continuous amyloid (Centiloid) | LC-related brain pattern × amyloid × time | LC | Interaction | 0.015 | 0.148 | — |

| Model type | Outcome / Interaction | Region | Effect | β / OR | P | $P_{FDR}$ |
|---|---|---|---|---|---|---|
| Amyloid-memory: continuous amyloid (Centiloid) | SN-VTA-related brain pattern × amyloid × time | SN-VTA | Interaction | 0.045 | $9.743 \times 10^{-4}$ | — |

Progression ORs are per one SD lower CR, holding the other nucleus constant. Mediation effects used the original latent-probit scale; their $P_{FDR}$ values use the two-nucleus family within endpoint and variant.

**Supplementary Table 20. Primary mediation of five-year clinical progression through longitudinal brain volume patterns**

| Region | Outcome | TPV adjustment | N | Effect | β | SE | CI lower | CI upper | P | $P_{FDR}$ |
|---|---|---|---|---|---|---|---|---|---|---|
| LC | MCI-to-AD dementia | Without TPV adjustment | 232 | indirect | −0.194 | 0.057 | −0.306 | −0.082 | $6.787 \times 10^{-4}$ | 0.001 |
| LC | MCI-to-AD dementia | Without TPV adjustment | 232 | direct | −0.178 | 0.092 | −0.358 | 0.003 | 0.054 | — |
| LC | MCI-to-AD dementia | Without TPV adjustment | 232 | total | −0.372 | 0.110 | −0.587 | −0.157 | $7.066 \times 10^{-4}$ | — |
| LC | MCI-to-AD dementia | With TPV adjustment | 232 | indirect | −0.148 | 0.049 | −0.243 | −0.052 | 0.002 | 0.005 |
| LC | MCI-to-AD dementia | With TPV adjustment | 232 | direct | −0.214 | 0.100 | −0.410 | −0.018 | 0.033 | — |
| LC | MCI-to-AD dementia | With TPV adjustment | 232 | total | −0.362 | 0.108 | −0.573 | −0.150 | $7.958 \times 10^{-4}$ | — |
| LC | Any progression | Without TPV adjustment | 335 | indirect | −0.135 | 0.040 | −0.213 | −0.056 | $8.354 \times 10^{-4}$ | $9.061 \times 10^{-4}$ |

| Region | Outcome | TPV adjustment | N | Effect | β | SE | CI lower | CI upper | P | $P_{FDR}$ |
|---|---|---|---|---|---|---|---|---|---|---|
| LC | Any progression | Without TPV adjustment | 335 | direct | −0.090 | 0.068 | −0.223 | 0.043 | 0.183 | — |
| LC | Any progression | Without TPV adjustment | 335 | total | −0.225 | 0.078 | −0.378 | −0.071 | 0.004 | — |
| LC | Any progression | With TPV adjustment | 335 | indirect | −0.091 | 0.029 | −0.149 | −0.034 | 0.002 | 0.004 |
| LC | Any progression | With TPV adjustment | 335 | direct | −0.135 | 0.082 | −0.295 | 0.026 | 0.099 | — |
| LC | Any progression | With TPV adjustment | 335 | total | −0.226 | 0.089 | −0.400 | −0.053 | 0.011 | — |
| SN-VTA | MCI-to-AD dementia | Without TPV adjustment | 232 | indirect | −0.158 | 0.059 | −0.273 | −0.042 | 0.007 | 0.007 |
| SN-VTA | MCI-to-AD dementia | Without TPV adjustment | 232 | direct | 0.034 | 0.154 | −0.269 | 0.336 | 0.828 | — |
| SN-VTA | MCI-to-AD dementia | Without TPV adjustment | 232 | total | −0.124 | 0.142 | −0.402 | 0.154 | 0.381 | — |

| Region | Outcome | TPV adjustment | N | Effect | β | SE | CI lower | CI upper | P | $P_{FDR}$ |
|---|---|---|---|---|---|---|---|---|---|---|
| SN-VTA | MCI-to-AD dementia | With TPV adjustment | 232 | indirect | −0.085 | 0.042 | −0.168 | −0.002 | 0.044 | 0.044 |
| SN-VTA | MCI-to-AD dementia | With TPV adjustment | 232 | direct | 0.037 | 0.139 | −0.235 | 0.309 | 0.791 | — |
| SN-VTA | MCI-to-AD dementia | With TPV adjustment | 232 | total | −0.048 | 0.132 | −0.307 | 0.211 | 0.715 | — |
| SN-VTA | Any progression | Without TPV adjustment | 335 | indirect | −0.145 | 0.044 | −0.231 | −0.059 | $9.061 \times 10^{-4}$ | $9.061 \times 10^{-4}$ |
| SN-VTA | Any progression | Without TPV adjustment | 335 | direct | −0.045 | 0.133 | −0.305 | 0.215 | 0.733 | — |
| SN-VTA | Any progression | Without TPV adjustment | 335 | total | −0.190 | 0.126 | −0.436 | 0.056 | 0.130 | — |
| SN-VTA | Any progression | With TPV adjustment | 335 | indirect | −0.062 | 0.029 | −0.120 | −0.004 | 0.035 | 0.035 |
| SN-VTA | Any progression | With TPV adjustment | 335 | direct | −0.047 | 0.111 | −0.264 | 0.169 | 0.668 | — |

| Region | Outcome | TPV adjustment | N | Effect | β | SE | CI lower | CI upper | P | $P_{FDR}$ |
|---|---|---|---|---|---|---|---|---|---|---|
| SN-VTA | Any progression | With TPV adjustment | 335 | total | −0.109 | 0.106 | −0.317 | 0.098 | 0.301 | — |

SE, standard error. P, P value. CI, confidence interval.

CR, contrast ratio. TPV, total parenchymal volume.

Effects are conditional latent-probit SEM estimates. $P_{FDR}$ is the original two-nucleus indirect-effect family within each endpoint and model variant; direct and total effects retain raw P.

**Supplementary Table 21. Mediation using PLSC patterns estimated in CN and MCI participants**

| Region | Outcome | TPV adjustment | N | Effect | β | SE | CI lower | CI upper | P | $P_{FDR}$ |
|---|---|---|---|---|---|---|---|---|---|---|
| LC | MCI-to-AD dementia | Without TPV adjustment | 232 | indirect | −0.193 | 0.060 | −0.311 | −0.075 | 0.001 | 0.003 |
| LC | MCI-to-AD dementia | Without TPV adjustment | 232 | direct | −0.178 | 0.094 | −0.362 | 0.007 | 0.059 | — |
| LC | MCI-to-AD dementia | Without TPV adjustment | 232 | total | −0.371 | 0.110 | −0.586 | −0.157 | $7.008 \times 10^{-4}$ | — |
| LC | MCI-to-AD dementia | With TPV adjustment | 232 | indirect | −0.149 | 0.050 | −0.247 | −0.052 | 0.003 | 0.005 |
| LC | MCI-to-AD dementia | With TPV adjustment | 232 | direct | −0.209 | 0.100 | −0.406 | −0.013 | 0.037 | — |
| LC | MCI-to-AD dementia | With TPV adjustment | 232 | total | −0.359 | 0.107 | −0.568 | −0.149 | $7.875 \times 10^{-4}$ | — |
| LC | Any progression | Without TPV adjustment | 335 | indirect | −0.140 | 0.042 | −0.223 | −0.057 | $9.341 \times 10^{-4}$ | $9.341 \times 10^{-4}$ |
| LC | Any progression | Without TPV adjustment | 335 | direct | −0.086 | 0.068 | −0.220 | 0.047 | 0.204 | — |
| LC | Any progression | Without TPV adjustment | 335 | total | −0.226 | 0.079 | −0.381 | −0.071 | 0.004 | — |
| LC | Any progression | With TPV adjustment | 335 | indirect | −0.094 | 0.029 | −0.151 | −0.036 | 0.001 | 0.003 |
| LC | Any progression | With TPV adjustment | 335 | direct | −0.132 | 0.082 | −0.291 | 0.028 | 0.107 | — |
| LC | Any progression | With TPV adjustment | 335 | total | −0.225 | 0.088 | −0.398 | −0.052 | 0.011 | — |
| SN-VTA | MCI-to-AD dementia | Without TPV adjustment | 232 | indirect | −0.162 | 0.058 | −0.276 | −0.047 | 0.006 | 0.006 |
| SN-VTA | MCI-to-AD dementia | Without TPV adjustment | 232 | direct | 0.037 | 0.154 | −0.265 | 0.340 | 0.809 | — |

| Region | Outcome | TPV adjustment | N | Effect | β | SE | CI lower | CI upper | P | $P_{FDR}$ |
|---|---|---|---|---|---|---|---|---|---|---|
| SN-VTA | MCI-to-AD dementia | Without TPV adjustment | 232 | total | −0.124 | 0.142 | −0.403 | 0.154 | 0.381 | — |
| SN-VTA | MCI-to-AD dementia | With TPV adjustment | 232 | indirect | −0.089 | 0.042 | −0.171 | −0.007 | 0.034 | 0.034 |
| SN-VTA | MCI-to-AD dementia | With TPV adjustment | 232 | direct | 0.040 | 0.139 | −0.233 | 0.313 | 0.772 | — |
| SN-VTA | MCI-to-AD dementia | With TPV adjustment | 232 | total | −0.048 | 0.132 | −0.308 | 0.211 | 0.715 | — |
| SN-VTA | Any progression | Without TPV adjustment | 335 | indirect | −0.148 | 0.044 | −0.233 | −0.062 | $7.353 \times 10^{-4}$ | $9.341 \times 10^{-4}$ |
| SN-VTA | Any progression | Without TPV adjustment | 335 | direct | −0.042 | 0.133 | −0.303 | 0.218 | 0.750 | — |
| SN-VTA | Any progression | Without TPV adjustment | 335 | total | −0.190 | 0.126 | −0.436 | 0.056 | 0.130 | — |
| SN-VTA | Any progression | With TPV adjustment | 335 | indirect | −0.065 | 0.030 | −0.123 | −0.006 | 0.031 | 0.031 |
| SN-VTA | Any progression | With TPV adjustment | 335 | direct | −0.045 | 0.111 | −0.262 | 0.172 | 0.686 | — |
| SN-VTA | Any progression | With TPV adjustment | 335 | total | −0.109 | 0.106 | −0.317 | 0.098 | 0.302 | — |

SE, standard error. P, P value. CI, confidence interval. CR, contrast ratio. TPV, total parenchymal volume.

Effects are conditional latent-probit SEM estimates, not logistic odds ratios. $P_{FDR}$ is the original two-nucleus indirect-effect family within each endpoint and model variant; direct and total effects retain raw P.

**Supplementary Table 22. Temporal-order mediation sensitivity: T1 after LC/SN-VTA MRI**

| Region | Outcome | Model type | N | Effect | β | SE | CI lower | CI upper | P | $P_{FDR}$ |
|---|---|---|---|---|---|---|---|---|---|---|
| LC | MCI-to-AD dementia | Original PLSC | 150 | indirect | −0.184 | 0.078 | −0.337 | −0.030 | 0.019 | 0.038 |
| LC | MCI-to-AD dementia | Original PLSC | 150 | direct | −0.296 | 0.180 | −0.648 | 0.057 | 0.100 | — |
| LC | MCI-to-AD dementia | Original PLSC | 150 | total | −0.479 | 0.217 | −0.904 | −0.055 | 0.027 | — |
| LC | Any progression | Original PLSC | 235 | indirect | −0.106 | 0.047 | −0.198 | −0.014 | 0.025 | 0.049 |
| LC | Any progression | Original PLSC | 235 | direct | −0.228 | 0.137 | −0.497 | 0.040 | 0.096 | — |
| LC | Any progression | Original PLSC | 235 | total | −0.334 | 0.138 | −0.604 | −0.063 | 0.016 | — |
| SN-VTA | MCI-to-AD dementia | Original PLSC | 151 | indirect | −0.096 | 0.086 | −0.264 | 0.072 | 0.262 | 0.262 |
| SN-VTA | MCI-to-AD dementia | Original PLSC | 151 | direct | −0.071 | 0.228 | −0.518 | 0.376 | 0.757 | — |
| SN-VTA | MCI-to-AD dementia | Original PLSC | 151 | total | −0.167 | 0.225 | −0.609 | 0.275 | 0.459 | — |
| SN-VTA | Any progression | Original PLSC | 237 | indirect | −0.086 | 0.055 | −0.194 | 0.023 | 0.122 | 0.122 |
| SN-VTA | Any progression | Original PLSC | 237 | direct | 0.065 | 0.151 | −0.231 | 0.361 | 0.666 | — |
| SN-VTA | Any progression | Original PLSC | 237 | total | −0.021 | 0.143 | −0.300 | 0.259 | 0.886 | — |
| LC | MCI-to-AD dementia | Re-estimated PLSC | 150 | indirect | −0.213 | 0.085 | −0.379 | −0.046 | 0.012 | 0.025 |
| LC | MCI-to-AD dementia | Re-estimated PLSC | 150 | direct | −0.270 | 0.179 | −0.621 | 0.080 | 0.131 | — |

| Region | Outcome | Model type | N | Effect | β | SE | CI lower | CI upper | P | $P_{FDR}$ |
|---|---|---|---|---|---|---|---|---|---|---|
| LC | MCI-to-AD dementia | Re-estimated PLSC | 150 | total | −0.483 | 0.220 | −0.914 | −0.052 | 0.028 | — |
| LC | Any progression | Re-estimated PLSC | 235 | indirect | −0.137 | 0.055 | −0.245 | −0.030 | 0.013 | 0.025 |
| LC | Any progression | Re-estimated PLSC | 235 | direct | −0.202 | 0.144 | −0.484 | 0.079 | 0.159 | — |
| LC | Any progression | Re-estimated PLSC | 235 | total | −0.340 | 0.142 | −0.617 | −0.062 | 0.017 | — |
| SN-VTA | MCI-to-AD dementia | Re-estimated PLSC | 151 | indirect | −0.137 | 0.094 | −0.322 | 0.048 | 0.146 | 0.146 |
| SN-VTA | MCI-to-AD dementia | Re-estimated PLSC | 151 | direct | −0.033 | 0.232 | −0.488 | 0.421 | 0.886 | — |
| SN-VTA | MCI-to-AD dementia | Re-estimated PLSC | 151 | total | −0.170 | 0.231 | −0.622 | 0.282 | 0.461 | — |
| SN-VTA | Any progression | Re-estimated PLSC | 237 | indirect | −0.109 | 0.062 | −0.231 | 0.013 | 0.081 | 0.081 |
| SN-VTA | Any progression | Re-estimated PLSC | 237 | direct | 0.088 | 0.153 | −0.213 | 0.389 | 0.567 | — |
| SN-VTA | Any progression | Re-estimated PLSC | 237 | total | −0.021 | 0.145 | −0.304 | 0.262 | 0.886 | — |

SE, standard error. P, P value. CI, confidence interval. CR, contrast ratio. TPV, total parenchymal volume.

Effects are conditional latent-probit SEM estimates. $P_{FDR}$ is the original two-nucleus indirect-effect family within each endpoint and model variant; direct and total effects retain raw P.

**Supplementary Table 23. Amyloid-memory interaction**

| Region | Amyloid encoding | N | Observations | Interaction | β | SE | CI lower | CI upper | P |
|---|---|---|---|---|---|---|---|---|---|
| LC | Binary amyloid | 122 | 562 | LC CR × amyloid × time | 0.020 | 0.015 | −0.009 | 0.049 | 0.175 |
| LC | Binary amyloid | 122 | 562 | LC-related brain pattern × amyloid × time | 0.029 | 0.019 | −0.009 | 0.066 | 0.132 |
| LC | Binary amyloid | 122 | 562 | TPV × amyloid × time | 0.040 | 0.016 | 0.009 | 0.071 | 0.012 |
| SN-VTA | Binary amyloid | 122 | 562 | SN-VTA CR × amyloid × time | −0.021 | 0.015 | −0.050 | 0.008 | 0.153 |
| SN-VTA | Binary amyloid | 122 | 562 | SN-VTA-related brain pattern × amyloid × time | 0.088 | 0.025 | 0.038 | 0.138 | $5.292 \times 10^{-4}$ |
| SN-VTA | Binary amyloid | 122 | 562 | TPV × amyloid × time | 0.005 | 0.022 | −0.038 | 0.047 | 0.824 |
| LC | Continuous amyloid (Centiloid) | 122 | 562 | LC CR × amyloid × time | 0.011 | 0.008 | −0.005 | 0.026 | 0.168 |
| LC | Continuous amyloid (Centiloid) | 122 | 562 | LC-related brain pattern × amyloid × time | 0.015 | 0.011 | −0.005 | 0.036 | 0.148 |
| LC | Continuous amyloid (Centiloid) | 122 | 562 | TPV × amyloid × time | 0.003 | 0.008 | −0.013 | 0.020 | 0.694 |
| SN-VTA | Continuous amyloid (Centiloid) | 122 | 562 | SN-VTA CR × amyloid × time | 0.005 | 0.008 | −0.011 | 0.020 | 0.565 |
| SN-VTA | Continuous amyloid (Centiloid) | 122 | 562 | SN-VTA-related brain pattern × amyloid × time | 0.045 | 0.014 | 0.018 | 0.072 | $9.743 \times 10^{-4}$ |

| Region | Amyloid encoding | N | Observations | Interaction | β | SE | CI lower | CI upper | P |
|---|---|---|---|---|---|---|---|---|---|
| SN-VTA | Continuous amyloid (Centiloid) | 122 | 562 | TPV × amyloid × time | −0.010 | 0.011 | −0.031 | 0.011 | 0.351 |

SE, standard error. P, P value. CI, confidence interval.

TPV, total parenchymal volume.
